# Multisensory approach in Mental Imagery: ALE meta-analyses comparing Motor, Visual and Auditory Imagery

**DOI:** 10.1101/2024.09.02.610739

**Authors:** Elise E Van Caenegem, Marcos Moreno-Verdú, Baptiste M Waltzing, Gautier Hamoline, Siobhan M McAteer, Frahm Lennart, Robert M Hardwick

## Abstract

Mental Imagery is a topic of longstanding and widespread scientific interest. Individual studies have typically focused on a single modality (e.g. Motor, Visual, Auditory) of Mental Imagery. Relatively little work has considered directly comparing and contrasting the brain networks associated with these different modalities of Imagery. The present study integrates data from 439 neuroimaging experiments to identify both modality-specific and shared neural networks involved in Mental Imagery. Comparing the networks involved in Motor, Visual, and Auditory Imagery identified a pattern whereby each form of Imagery preferentially recruited ‘higher level’ associative brain regions involved in the associated ‘real’ experience. Results also indicate significant overlap in a left-lateralized network including the pre-supplementary motor area, ventral premotor cortex and inferior parietal lobule. This pattern of results supports the existence of a ‘core’ network that supports the attentional, spatial, and decision-making demands of Mental Imagery. Together these results offer new insights into the brain networks underlying human imagination.

**Highlights:**

- Meta-Analyses of Motor, Visual, and Auditory modalities of Imagery were compared
- Each modality recruited high level areas related to associated real-life experience
- A ‘core’ network of pre-SMA, PMv and IPL were recruited by all imagery modalities
- Results emphasize attentional, spatial, & executive aspects of Mental Imagery

## 1 Introduction

Mental Imagery is a concept that has been explored across a diverse range of fields such as sport performance (Ladda et al., 2021), motor skill learning (Lotze & Halsband, 2006; Williams & Gribble, 2012), rehabilitation (Ietswaart et al., 2011; Jackson et al., 2001; Malouin & Richards, 2010), music (Izadifar et al., 2022; Tanaka & Kirino, 2017; Tsai et al., 2018), people with disabilities (e.g. blind, deaf; Lazard et al., 2011; Nierhaus et al., 2023) and many other domains. It is a complex cognitive ability that allows an individual to generate internal sensory experiences, contributing to creativity, memory, planning, and other aspects of cognition (Kosslyn et al., 2001). Imagery can generally be defined as the mental faculty of creating sensory representations in the mind, without these stimuli being present in the real environment. Imagery can encompass all five senses, and may emphasize different sensory modalities, such as Auditory Imagery (imagining sounds or voices), Olfactory Imagery (imagining scents), Gustatory Imagery (imagining tastes), Tactile Imagery (imagining touch perceptions), Visual Imagery (imagining visual scenes, pictures, objects, colours), and Kinesthetic Imagery (imagining muscle tension and proprioception). There are also types of imagery that can involve several imagery modalities; for example, Motor Imagery (imagining performing movements) often focuses on combinations of Visual and Kinesthetic Imagery.

Previous research has established a close link between imagery and actual perception or execution, suggesting that these processes share similarities. For example, based on the long-established concept of ’Functional Equivalence’ (Finke, 1980), Motor Simulation Theory (Jeannerod, 2001) proposes that Motor Imagery uses the same brain networks and processes involved in the physical execution of movements. Motor Simulation Theory therefore argues that motor network plays a central role in Motor Imagery. Research exploring other modalities of imagery has identified similar relationships between imagery and perception (Kosslyn et al., 1997; Kosslyn & Thompson, 2003; Plailly et al., 2008; Small & Prescott, 2005; Zatorre & Halpern, 2005). Hétu et al. (2013) proposed that such similarities could result from evolutionary processes resulting from the anatomical and computational efficiency of sharing circuitry between imagery and real-life experiences. Extending this logic, it seems plausible that different modalities of imagery may themselves also share common neural circuitry. More specifically, Imagery may involve the recruitment of both ‘task-specific’ regions closely related to their associated physical experiences, alongside regions representing a globally shared ‘core network’ that is more directly responsible for the process of generating Mental Imagery itself.

To date, most previous studies have investigated different modalities of imagery individually, making it difficult to understand what might be unique or global to different modalities of imagery. For example, neuroimaging meta-analyses have generally focused on one modality of Imagery (Hardwick et al., 2018; Hétu et al., 2013; Spagna et al., 2021; Winlove et al., 2018). To the best of our knowledge, only McNorgan (2012) has previously investigated multiple different modalities of imagery, examining Auditory, Tactile, Motor, Gustatory, Olfactory, and different forms of Visual imagery. When combining data across all modalities into a single meta-analysis, the results identified a primarily left-lateralized network, with bilateral parietal involvement. While analysis of individual sensory modalities primarily identified left-localized networks, unfortunately, these results may have been affected by a relatively low sample size; a mean average of 10 experiments were available for each individual modality, and subsequent work has identified the minimum sample size to provide adequate power for the meta-analysis technique used in this study is 17 experiments (Eickhoff et al., 2016). Consequently, more detailed analyses, such as performing conjunctions to identify which regions may have been consistently recruited across different modalities of imagery, were not possible. Given that more than a decade has elapsed since the last synthesis, at the present time, there is an order of magnitude more papers available for several of the modalities of imagery examined, making it possible to examine both the different modalities of imagery, and the relationship between them, in greater depth.

The present study therefore examined the neural networks activated during different modalities of imagery, with the goal to examine both the specific neural networks activated for different modalities of imagery, and to identify regions that may be recruited more generally across multiple modalities of imagery. Using the Activation Likelihood Estimation (ALE) method, we obtained a quantitative summary of the available neuroimaging data examining Mental Imagery. We hypothesised that these individual networks may also share common activations, representing a ‘core’ network central to the process of generating Mental Imagery itself.

## 2 Methods

### 2.1 Literature searches

Relevant neuroimaging papers were identified via literature searches on PubMed, conducted up to June 2023. Specific search strings used are provided in Table 1. In addition to the search strings, we also included data identified in previous ALE meta-analyses where possible (e.g. data for Motor Imagery by Hardwick et al. (2018), and from McNorgan (2012)).

**Table 1:**
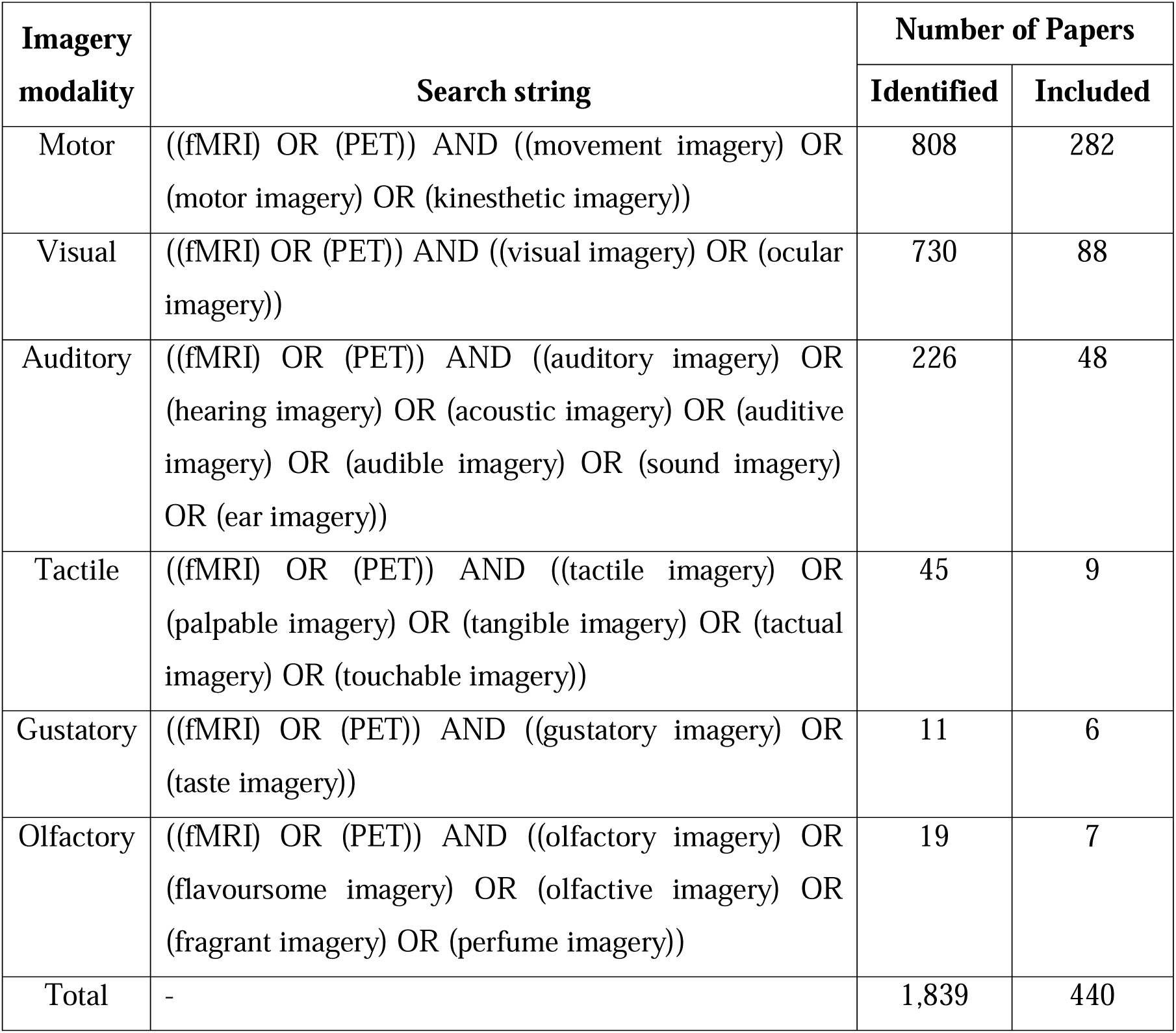
PubMed search strings and associated papers.

| Imagery modality | Search string | Number of Papers |  |
| --- | --- | --- | --- |
|  |  | Identified | Included |
| Motor | ((fMRI) OR (PET)) AND ((movement imagery) OR (motor imagery) OR (kinesthetic imagery)) | 808 | 282 |
| Visual | ((fMRI) OR (PET)) AND ((visual imagery) OR (ocular imagery)) | 730 | 88 |
| Auditory | ((fMRI) OR (PET)) AND ((auditory imagery) OR (hearing imagery) OR (acoustic imagery) OR (auditive imagery) OR (audible imagery) OR (sound imagery) OR (ear imagery)) | 226 | 48 |
| Tactile | ((fMRI) OR (PET)) AND ((tactile imagery) OR (palpable imagery) OR (tangible imagery) OR (tactual imagery) OR (touchable imagery)) | 45 | 9 |
| Gustatory | ((fMRI) OR (PET)) AND ((gustatory imagery) OR (taste imagery)) | 11 | 6 |
| Olfactory | ((fMRI) OR (PET)) AND ((olfactory imagery) OR (flavoursome imagery) OR (olfactive imagery) OR (fragrant imagery) OR (perfume imagery)) | 19 | 7 |
| Total | - | 1,839 | 440 |

### 2.2 Inclusion/Exclusion criteria

As is standard for ALE meta-analysis, the analyses included only experiments that involved coordinates from whole-brain analyses, excluding papers focusing only on specific regions of interest (ROI). The ALE method examines the clustering of peak coordinates, and as such only those studies reporting their results in the standardized Montreal Neurological Institute (MNI) or Talairach and Tornaux (TAL) space were included. The selected experiments had to report data from healthy individuals with normal or corrected-to-normal vision, and no history of neurological disease. In instances where available, data from healthy control groups from patient studies (but not data from the patient groups themselves) were also included. The meta-analysis concentrated on within-subject contrasts to avoid comparisons with patient groups/across groups of unequal size. Also, only papers providing explicit forms of imagery were included.

### 2.3 Data extraction and classification

Each paper was given a unique identifier based on the surname of the first author and the year of publication (letters a, b, c etc. were appended in cases where duplicates occurred). Data extracted from each paper included the number of participants in each experiment and the coordinates of the reported activations in MNI or Talairach space. Coordinates reported in Talairach space were converted to MNI space using the Lancaster transform (Lancaster et al., 2007). Each experiment was categorized as involving Motor, Visual, Auditory, Tactile, Gustatory or Olfactory Imagery. For ease of reference, we also identified the source of the data within the paper (e.g. Table number), and the DOI. A summary of the data included in each analysis is presented in Table 2. More detailed information on the individual experiments included in each analysis is presented in Supplementary Table S85.

**Table 2.** Data included in the meta-analyses.

| Imagery Modalities | Experiments | Participants | Foci |
| --- | --- | --- | --- |
| All | 439 | 7366 | 6386 |
| Motor | 284 | 4882 | 4309 |
| Visual | 92 | 1337 | 1310 |
| Auditory | 48 | 793 | 611 |
| *Tactile | 9 | 126 | 111 |
| *Gustatory | 7 | 102 | 53 |
| *Olfactory | 7 | 200 | 50 |

\* Note that these imagery modalities did not have enough experiments ( $n < 17$ ) for an adequately powered meta-analysis (Eickhoff et al., 2016). Exploratory analyses examining these modalities of imagery are presented in the supplementary materials associated with this paper.

### 2.4 Data analyses

#### 2.4.1 General Procedures

In a first step, separate meta-analyses identified the individual networks involved in each modality of imagery We note, however, that only Motor, Visual, and Auditory imagery provided enough studies to allow us to conduct suitably powered ALE analyses (according to Eickhoff et al. (2016), a minimum of 17 studies are required). Consequently, only three of the original six modalities will be presented in detail in the main manuscript (though note that exploratory analyses for Tactile, Gustatory, and Olfactory imagery are presented in the supplementary materials – see (Supplementary Tables S5-S6)We then used pairwise conjunction analyses to assess the convergence and divergence between the individual modality networks and combined their results to identify regions consistently recruited across all imagery modalities. Differences between the individual modalities were examined using pairwise contrast analyses. Finally, we conducted an ALE meta-analysis with all data combined, in order to identify the general brain network involved in the generation of Mental Imagery itself, regardless of the specific modality of imagery used. All the results tables can be found in Supplementary Materials, alongside a series of control analyses (e.g. contrasts that compared imagery vs rest conditions)

Given that the imagery modalities grouped together all the types of tasks, it was possible to find divergences within these categories. To analyse each modality of imagery in more detail, further sub-analyses were carried out and the results are presented in the Supplementary Tables S83-S84. For Visual Imagery, the separation into three sub-categories (shapes, colour, movement) made by McNorgan (2012) was followed. Visual shapes imagery refers to the imagination of objects, shapes and landscapes. Visual motion imagery, on the other hand, is about imagining objects or shapes in motion, excluding any imagery involving the human body. Finally, Visual colour imagery is the imagination of colour. For Motor Imagery, no sub-analyses were performed given the problem highlighted by Van Caenegem et al. (2022) regarding the lack of information in the description of Motor Imagery modalities. For Auditory Imagery, we did not split into categories because there were not enough papers to establish categories.

#### 2.4.2 ALE analyses

The analyses were carried out using the revised version of the ALE algorithm (Eickhoff et al., 2009; Turkeltaub et al., 2002, 2012). This ALE approach determines whether the convergence of activation coordinates (foci) across various experiments occurs at a level greater than what would be expected by chance. The reported foci are represented as the centres of 3D Gaussian probability distributions (Turkeltaub et al., 2002). In the revised algorithm, the width of these Gaussians is determined through empirical comparisons between subjects and templates, allowing the increased spatial reliability associated with larger sample sizes to be modelled by employing smaller Gaussian distributions (Eickhoff et al., 2009).

Foci from each experiment were aggregated across voxels to generate a modelled activation map (Turkeltaub et al., 2012). The combination of these modelled activation maps across experiments resulted in ALE scores, representing the convergence of coordinates for each location. These ALE scores were then compared to a non-linear histogram integration based on the frequency of distinct modelled activation maps (Eickhoff et al., 2012). This comparison determined areas where the convergence exceeded what would be expected by chance. ALE values were calculated exclusively for voxels with a probability of ≥10% of containing grey matter (Evans et al., 1994), as functional activations are primarily observed in grey matter regions. The results were thresholded at p < 0.05 (cluster-level family-wise error, corrected for multiple comparisons, with a cluster-forming threshold at voxel level p < 0.001) and reported at a voxel resolution of 2mm^3^. Results were reported with a minimum cluster volume of 100mm^3^ (i.e. >13 voxels, Beissner et al., 2013; Erickson et al., 2014; Turkeltaub et al., 2012).

#### 2.4.3 Conjunction analyses

Conjunction analyses were employed to synthesize the overlap between networks. These analyses adhered to the conjunction null hypothesis and were computed utilizing the minimum statistic method (Nichols et al., 2005), incorporating a minimum cluster volume of 100mm^3^ (Beissner et al., 2013; Erickson et al., 2014; Turkeltaub et al., 2012). In a final step we conducted a conjunction across each of the main analyses of different imagery modalities to identify neural substrates commonly recruited by all of them.

#### 2.4.4 Contrast analyses

To compare the resulting meta-analyses, random effects ALE subtraction analysis was employed (Eickhoff et al., 2012). In the initial step, voxel-wise differences between ALE maps were computed for each pool of experiments. Subsequently, experiments were randomly shuffled into two samples of equal size for the compared analyses, and voxel-wise differences between their ALE scores were recorded. This shuffling process was iterated 10,000 times to generate an empirical null distribution of ALE score differences between the conditions being compared. The map of differences obtained from this procedure was thresholded at a posterior probability for true differences of P > 0.95, and inclusively masked by the respective main effect of the minuend (Chase et al., 2011; Rottschy et al., 2012) with a minimum cluster volume of 100 mm^3^ (Beissner et al., 2013; Erickson et al., 2014; Turkeltaub et al., 2012).

#### 2.4.5 Labelling

The results were labelled anatomically based on their most likely macro-anatomical and cytoarchitectonic/tractographically assessed locations using the SPM Anatomy Toolbox 2 extension (Eickhoff et al., 2005, 2006, 2007). Furthermore, functional labels for motor regions were assigned using the human motor area template (HMAT) as defined by Mayka et al. (2006). Labels from Brodman area was added through a MRIcron Anatomical Template. The reported coordinates were based on peak maxima in MNI space.

## 3 Results

Our literature searches identified 439 experiments that could be included in one of our meta-analyses (see Table 2; details of the studies included are given in Supplementary Tables S85). Notably, of the six categories of imagery present, only three provided enough studies to allow us to conduct suitably powered ALE analyses (according to Eickhoff et al. (2016), a minimum of 17 studies is required to carry out analyses with enough power). We therefore focused our modality-specific analyses on those fields with suitable statistical power, i.e. Motor, Visual and Auditory Imagery. In order to provide a complete overview of the data available, exploratory analyses of results for other modalities of imagery (i.e. Tactile, Gustatory, and Olfactory) are presented in the supplementary materials.

### 3.1 Modality-Specific Analyses

#### 3.1.1 Motor Imagery

The analysis of Motor Imagery included 284 experiments reporting 4309 foci recorded from a total of 4882 participants. Motor Imagery identified the network with the greatest overall volume of the meta-analyses (a comparison of the volumes across networks is presented in Fig. 1). Motor Imagery mainly recruited bilateral premotor, parietal, and cerebellar regions, with left-lateralized recruitment of the dorsolateral prefrontal cortex (DLPFC). Two large clusters including bilateral ventral and dorsal premotor cortex were identified; both bilateral clusters extended to the supplementary motor area and the insula lobe, while the right cluster also extended to the putamen. While regions of this cluster extended into the primary motor cortex (M1), no peaks were identified in this area, suggesting activity did not originate from this region. Two bilateral parietal clusters spanned the inferior and superior parietal lobules, with right lateralized activations in the superior parietal lobule and the primary somatosensory cortex (S1). A left-lateralized activation of the Temporal lobe was also observed in the same cluster. Further bilateral clusters were identified in the cerebellum (lobule VI).

**Fig. 1.**
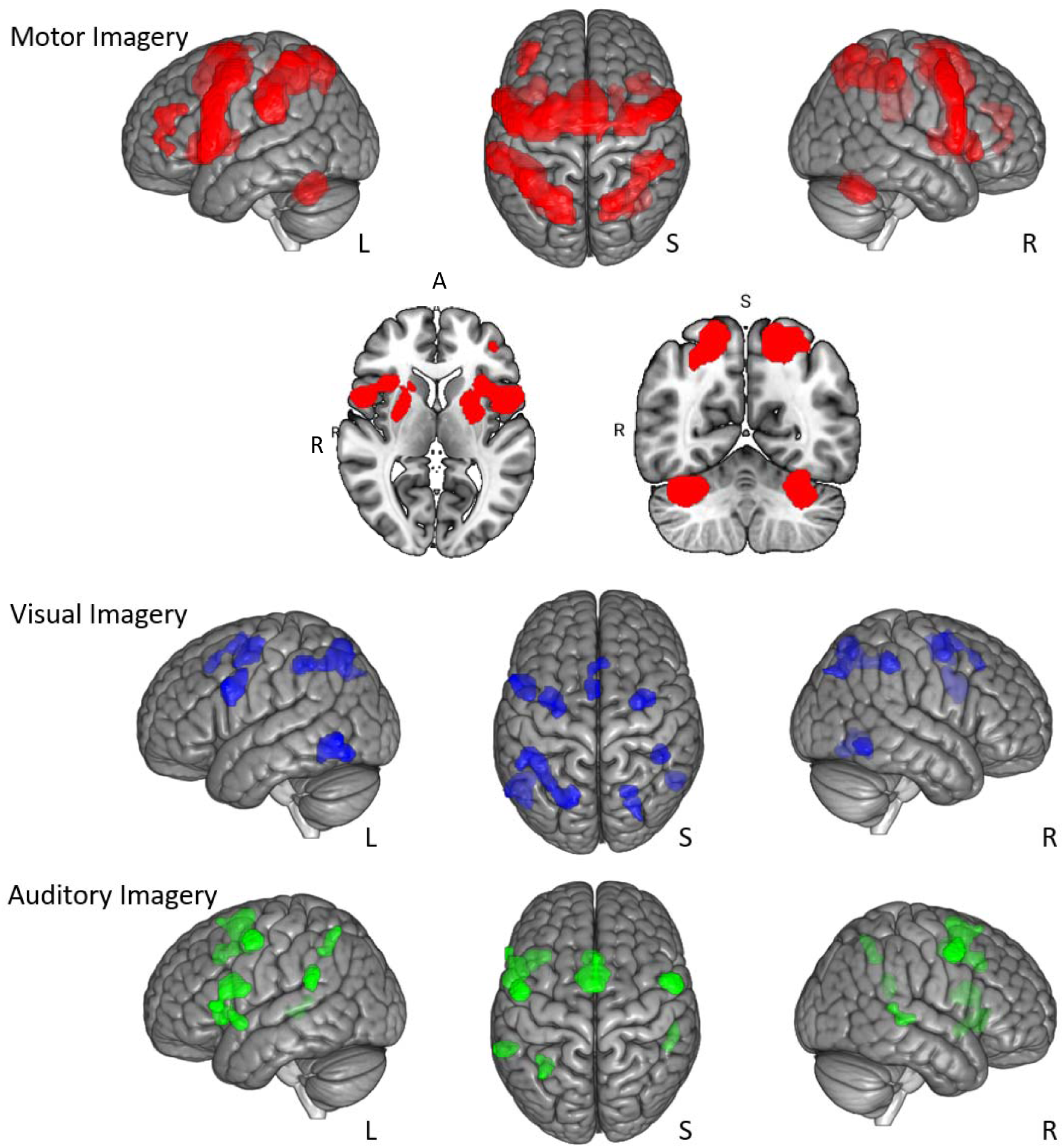
Quantitative meta-analyses of the three modalities.

#### 3.1.2 Visual Imagery

Visual Imagery included 92 experiments reporting 1310 foci recorded from a total of 1337 participants. Visual imagery recruited multiple parietal and frontal clusters. We observed a left-lateralized activation for the superior and inferior parietal lobule. Two small clusters spanned the dorsal and ventral premotor cortices on the left, while a third one included the right dorsal premotor cortex. A smaller cluster also identified activation in the pre-SMA region. Finally, we also identified a bilateral activation of the inferior temporal gyrus.

#### 3.1.3 Auditory Imagery

The analysis of Auditory Imagery involved 48 experiments reporting 611 foci recorded from a total of 793 participants. Auditory Imagery identified the smallest overall volume of the three meta-analyses. The largest cluster spanned the SMA and pre-SMA on both sides with maximum peaks over the left side. A second large cluster encompassed left-lateralized activation of the ventral premotor cortex, insula lobe, temporal pole, and primary somatosensory cortex. Two smaller clusters demonstrated bilateral activation of the dorsal premotor cortex. A final cluster identified activation of the left inferior parietal lobule.

#### 3.1.4 Volume comparison

We assessed the similarity and differences among each network by measuring the number of voxels within each network that coincided with other analyses, as well as the number of unique voxels in each analysis (Fig. 2.).

**Fig. 2.**
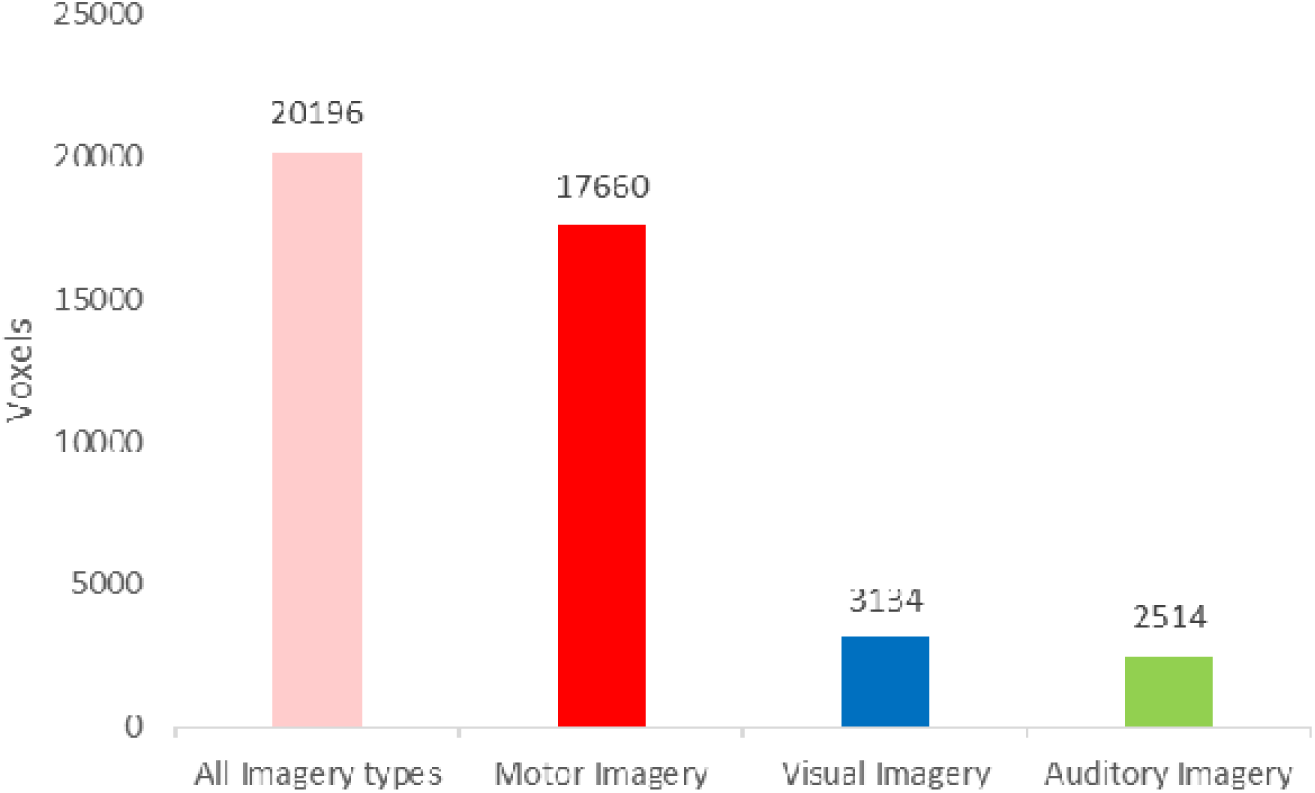
Volume Comparisons. Bar chart illustrates the number of voxels contributing to the volume for each modality.

#### 3.2 Conjunction analyses

Minimum-statistic conjunction analyses were conducted to identify regions consistently recruited across the different imagery modalities (Fig. 3; see also Supplementary Tables S8– S19).

**Fig. 3.**
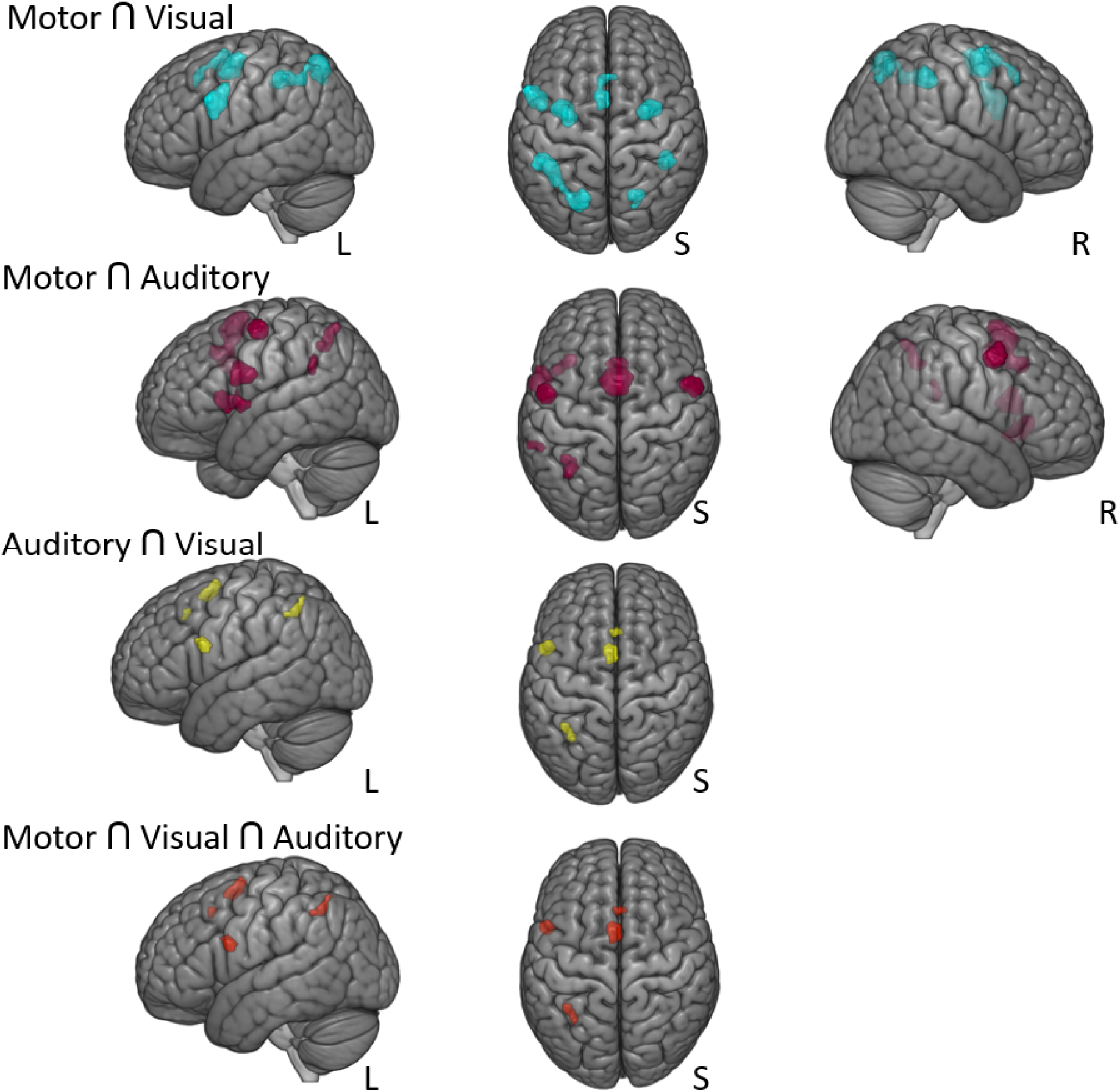
Conjunction analyses conducted across combinations of the imagery modalities.

#### 3.2.1 Motor Imagery ⍰ Visual Imagery

A conjunction between Motor Imagery and Visual Imagery identified a large cluster including the left superior and inferior parietal lobule. Three smaller clusters were found in the premotor cortex (left ventral and dorsal premotor cortex and right dorsal premotor cortex). A common activation of the pre-SMA was also demonstrated. Two relatively small clusters that encompass the right supra-marginal gyrus and the right cuneus were identified across both imagery conditions.

#### 3.2.2 Motor Imagery ⍰ Auditory Imagery

A conjunction between Motor Imagery and Auditory Imagery identified a network including the premotor cortex, parietal lobule, and temporal gyrus. In the premotor regions, one cluster included the left pre-SMA and SMA proper and two smaller clusters included the left and the right ventral premotor cortex. The second largest cluster encompassed left-lateralized regions such as the ventral premotor cortex, the insula lobe, the temporal pole, and the primary somatosensory cortex. Two further clusters were identified, one including the left inferior parietal lobule, while the second included the left superior temporal gyrus.

#### 3.2.3 Visual Imagery ⍰ Auditory Imagery

Consistent activations across Visual Imagery and Auditory Imagery were identified mainly in the premotor regions. Two small clusters included the pre-SMA on the left side. Another small cluster included the left ventral premotor cortex. The final cluster included the left inferior parietal lobule.

#### 3.2.4 Motor Imagery ⍰ Visual Imagery ⍰ Auditory Imagery

Finally, we computed a grand conjunction across Motor Imagery, Visual Imagery and Auditory Imagery. This analysis identified four small left-lateralized clusters. Separate premotor clusters spanned the pre-SMA (one more posterior-medial and the other one more superior-medial) and the ventral premotor cortex. In the parietal lobule, the cluster included a small part of the inferior parietal lobe.

### 3.3 Contrast analyses

Contrast analyses revealed regions that showed more consistent involvement with one modality compared to another (Fig. 4; see also Supplementary Tables S20–S38).

**Fig. 4.**
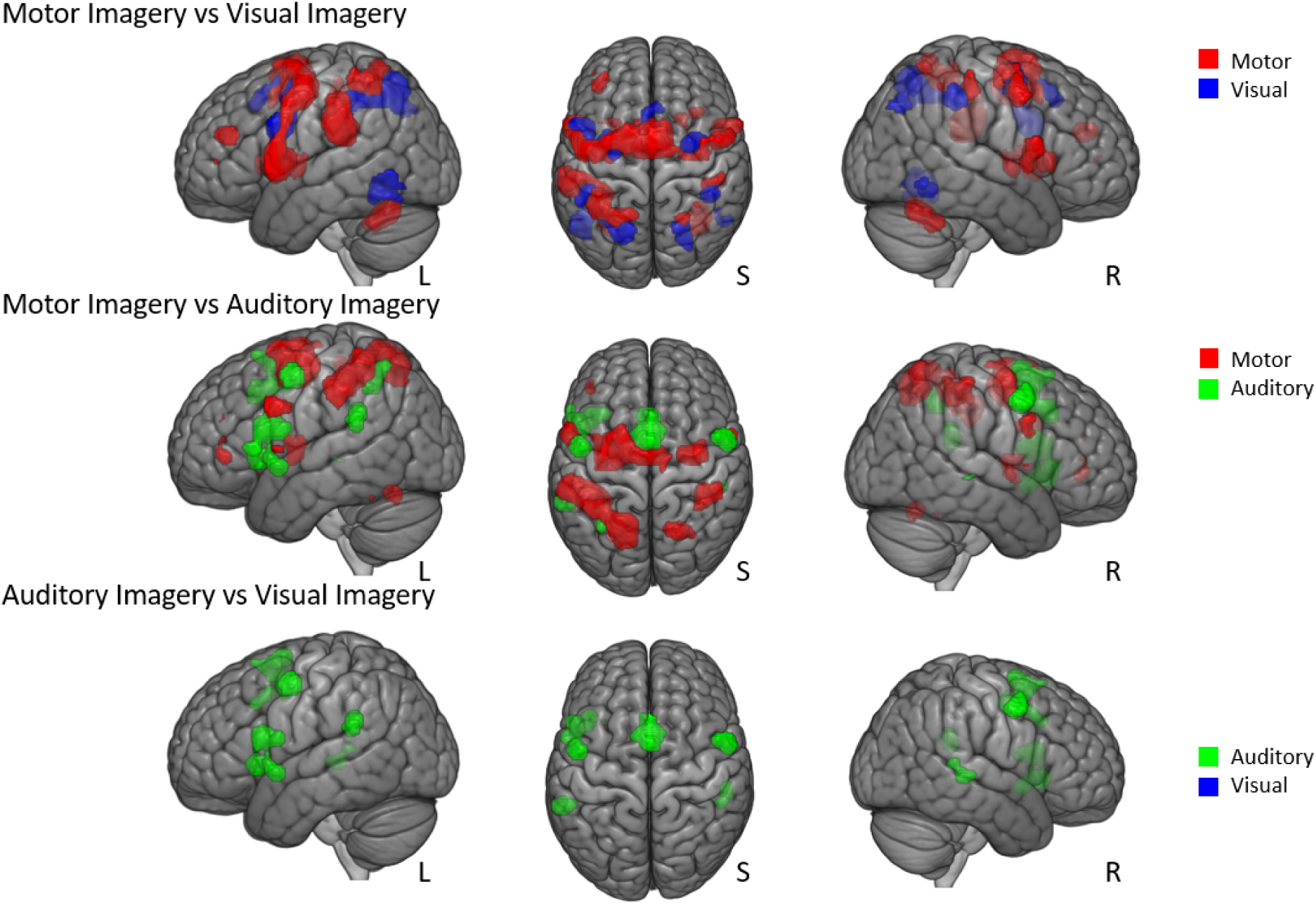
Contrast analyses.

#### 3.3.1 Motor Imagery vs Visual Imagery

Motor Imagery, when compared to Visual Imagery was more strongly associated with bilateral premotor regions such as pre-SMA, ventral and dorsal premotor cortex, and M1. Motor Imagery also recruited more parietal regions including the left inferior parietal lobule, the left supramarginal gyrus and the left superior parietal lobe. The left DLPFC, in the upper part of the frontal lobe, is also activated during Motor Imagery. Subcortically, the bilateral putamen and cerebellum were both more related to Motor Imagery.

Fewer voxels were more activated during Visual Imagery compared to Motor Imagery. These areas included more bilateral temporal and occipital regions, left pre-SMA and some smaller clusters around the left and right dorsal premotor cortex, and left ventral premotor cortex.

#### 3.3.2 Motor Imagery vs Auditory Imagery

Motor Imagery, compared to Auditory Imagery, was more consistently associated with recruiting premotor regions, including bilateral dorsal and ventral premotor cortex and bilateral SMA. A large volume spanning the left inferior and bilateral superior parietal cortex was also more consistently associated with Motor Imagery. There is also more activation in the primary somatosensory area and in the cerebellum during Motor Imagery compared to Auditory Imagery. In the frontal cortex, activation of the inferior frontal gyrus was noted.

By contrast, Auditory Imagery recruited more left lateralized pre-SMA area and a little part of the left inferior parietal lobule. Auditory Imagery also recruited more little parts of the bilateral dorsal premotor cortex and small temporal regions.

#### 3.3.3 Visual Imagery vs Auditory Imagery

Finally, Auditory Imagery, in comparison to Visual Imagery revealed a greater activation in several small clusters in premotor regions such as left pre-SMA, left ventral premotor cortex, bilateral dorsal premotor cortex, bilateral temporal regions and primary somatosensory area. In the opposite, Visual Imagery, compared to Auditory Imagery revealed no greater activation.

### 3.4 General Network for Mental Imagery

A final analysis considered the general brain network involved in Imagery regardless of the specific modality. This analysis therefore combined data from all available studies, combining data from Motor, Visual, Auditory, Tactile, Gustatory, and Olfactory modalities (Fig. 5; see also Supplementary Tables S1–S7). The analysis of all imagery modalities involved 439 experiments reporting 6386 foci recorded from a total of 7366 participants. This analysis revealed bilateral activations in the premotor cortex, parietal lobe, and temporal and occipital regions, with left-lateralized recruitment of the prefrontal cortex. The largest cluster recruited spanned several regions of the left hemisphere, including the temporal pole, the insula lobe, and the dorsolateral prefrontal cortex. Another large cluster, still left-lateralized, encompassed the inferior parietal lobule, the superior temporal pole, the supramarginal gyrus and the postcentral gyrus. In the right hemisphere, a cluster included the premotor cortex including the right dorsal and ventral premotor cortex extending to the right insula lobe, but also subcortical regions such as the right putamen and the cerebellum (right lobule VIIa and left lobule VI). A further cluster also included the right superior and inferior parietal lobule. Contrary to what may have been observed previously, a posterior left-lateralized activation was noted, including the inferior occipital gyrus, the inferior temporal gyrus, and the fusiform gyrus.

**Fig. 5.**
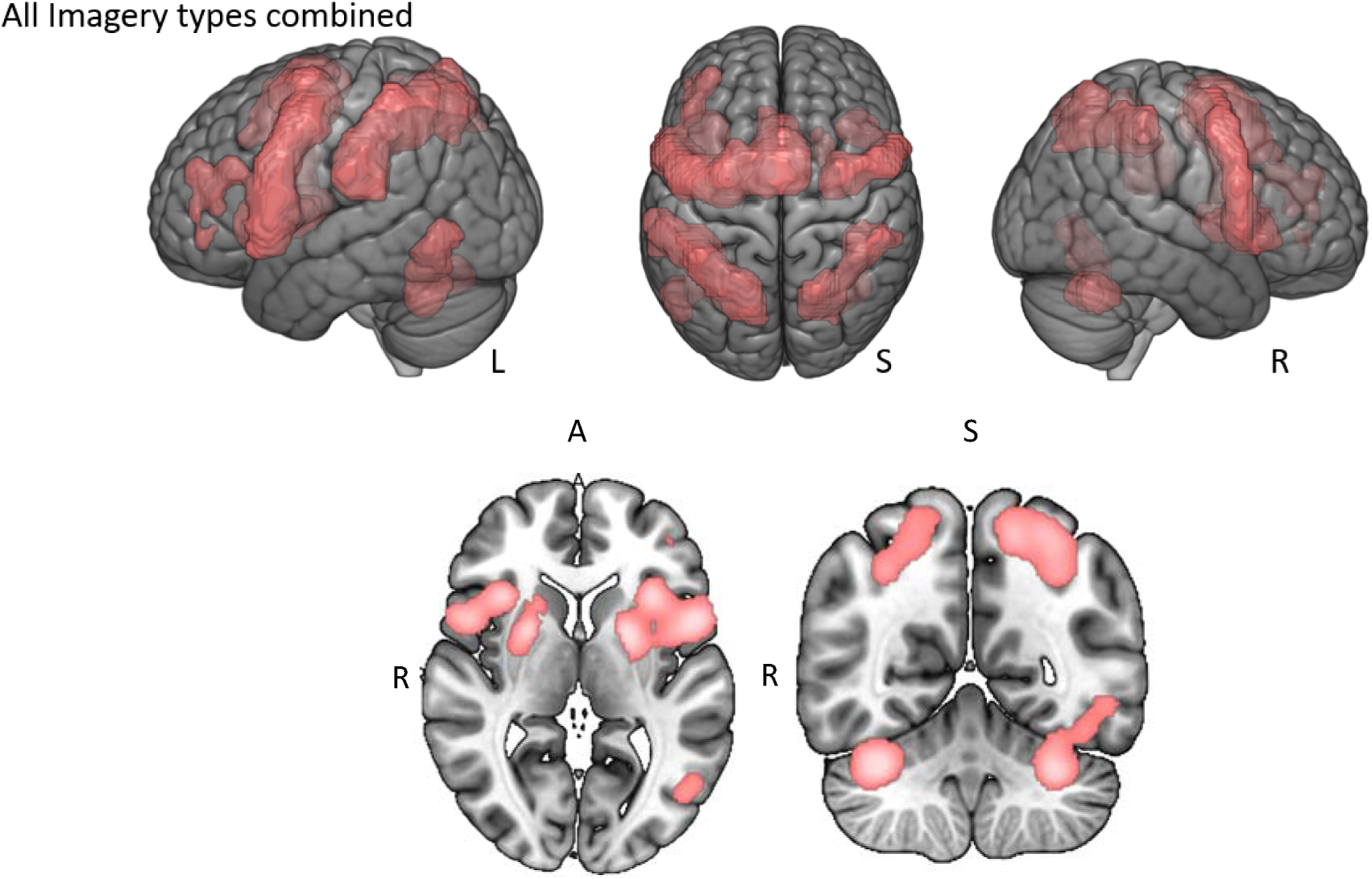
Quantitative meta-analyses of the combination of all imagery modalities (including Motor, Visual, Auditory, Tactile, Gustatory, and Olfactory).

## 4 Discussion

Imagery has been the subject of numerous studies, often focussing on a specific modality (e.g. Motor Imagery, Visual Imagery, Olfactory Imagery, etc). Most studies have proposed that imagery recruits similar brain regions to those involved in actual real-life experiences for that modality of imagery; relatively few studies have examined the possibility that different modalities of imagery may recruit common brain regions that are consistently involved in the generation of imagery itself. The present study therefore reviewed the available literature in order to consider imagery in the broadest sense of the term, across multiple different modalities. This question was examined using large-scale meta-analysis techniques, overcoming the limited sample sizes of previous individual studies, and providing quantitative and statistically principled results. Firstly, the network specific to each imagery modality was identified, compared, and contrasted. Examining these individual networks identified several important points, for example, showing that imagery appears to primarily recruit regions involved in ‘higher-level’ associative processing, as opposed to ‘lower-level’ primary sensorimotor brain functions, that recruitment of the prefrontal cortex was specific to Motor Imagery, and that activity of the temporal cortex was more related to Visual Imagery than other modalities. Conjunction analyses across multiple modality networks identified consistent activation of the pre-SMA, the ventral premotor cortex and the inferior parietal lobule across Motor, Visual, and Auditory Imagery. These results suggest these regions could represent a ‘core’ network that is consistently recruited during multiple modalities of Mental Imagery. Finally, by pooling data from all modalities of imagery available, a large neural network contributing to Imagery as a whole was identified. This network includes areas of the frontal, parietal and temporal cortex, as well as the cerebellum.

Alongside the novel analyses presented in this manuscript, our results replicate, update, and extend previous meta-analyses of the individual networks for Motor Imagery (Hardwick et al., 2018) and Visual Imagery (Spagna et al., 2021; Winlove et al., 2018). Since the outcomes of these individual analyses closely align with those of prior meta-analyses, we direct readers to these previous papers for a detailed examination of these individual networks. Our discussion will therefore concentrate on the novel insights derived from considering multiple different modalities of imagery.

### 4.1 Key observations

#### 4.1.1 Imagery preferentially recruits areas involved in higher-level associative brain functions

The present results provide considerable evidence of clusters originating in ‘higher-order’ associative areas for each sensory modality examined; for example, Motor Imagery recruited premotor and prefrontal regions implicated in motor planning, while Visual Imagery recruited premotor and parietal areas consistent with those of the dorsal and ventral visual systems (Goodale & Milner, 1992). By contrast, we found relatively little recruitment of ‘low-level’ or principle sensorimotor brain areas; more specifically, we found no clusters originating in M1 during Motor Imagery, nor was there evidence of clusters originating in the primary visual cortex (V1) during Visual Imagery. While our analysis of Auditory Imagery presented a notable exception to this finding by identifying clusters originating in the primary auditory cortex (A1), we note that previous research has argued A1 itself functions more like a higher-order region when considered relative to other brain networks (King & Nelken, 2009). The present results therefore provide evidence that imagery generally preferentially recruits regions involved in higher-order associative processing, though further discussion of these results is presented in the sections outlining the individual networks for each modality of imagery examined.

We note that this pattern of higher order associative recruitment appears to be relatively consistent across studies using whole-brain neuroimaging techniques such as fMRI and PET. For example, Hétu et al. (2013) found that only 22 of 122 studies examined in their meta-analysis of Motor Imagery reported recruitment of M1, and there was no activation of M1 in their meta-analysis. By contrast, studies that have delivered Transcranial Magnetic Stimulation (TMS) over M1 demonstrated modulation of corticospinal excitability during Motor Imagery (Fadiga et al., 1999; Grosprêtre et al., 2016). However, as the measurement technique used in such studies provides a general measurement of corticospinal excitability, whether this facilitation represents a direct activation of M1, or is mediated by a parallel pathway (e.g. premotor cortex to brainstem) remains an open question (for further discussion see Fadiga et al. (2005) and Barhoun et al. (2022)). Similarly, studies examining Motor Imagery using electroencephalography typically focus on electrodes that are considered to correspond to the position of M1 (although the exact location of these electrodes relative to underlying brain structure can vary, e.g. Silva et al. (2020)). As such, we caution against using our present results to argue strongly *against* the recruitment of ‘low order’ regions during imagery. It is, for example, plausible that lower-level regions could be recruited in a manner that traditional whole brain neuroimaging is not optimal to detect, and this question may be best addressed using more advanced analyses (e.g. multi-voxel pattern analysis (MVPA); see Albers et al., 2013), or techniques that complement whole brain neuroimaging such as (magneto)encephalogram and/or TMS studies. By contrast, the present results do strongly emphasize a pattern whereby different modalities of imagery consistently recruit higher-level associative brain regions involved in associated ‘real’ sensory experiences.

#### 4.1.2 Only Motor Imagery consistently recruited prefrontal regions

Mental Imagery has often been considered to be a complex cognitive process. Given the key role of the prefrontal cortex in cognitive control and executive function (Friedman & Robbins, 2022), and that recent work has proposed similarities between Motor Imagery and executive function (Glover et al., 2020; Glover & Baran, 2017; Martel & Glover, 2023), it seemed plausible that imagery may generally recruit prefrontal resources. However, we found that that consistent recruitment of the prefrontal cortex was unique to Motor Imagery. We found no evidence of prefrontal clusters in analyses examining other modalities of Imagery, including a control analysis in which we pooled the data from all studies except those using Motor Imagery in order to optimize statistical power (see Supplementary Materials Table S7). This lack of consistent prefrontal activation across analyses meant that the prefrontal cortex was also absent in conjunction analyses. Together these data suggest that consistent recruitment of the prefrontal cortex is unique to Motor Imagery (see also the specific section for further discussion of the possible roles of this structure during Motor Imagery).

### 4.2 Motor Imagery

Results of the present study are very similar to those observed in previous meta-analyses of Motor Imagery by Hardwick et al. (2018) and Hétu et al. (2013). Primary differences included larger overall volumes in the dorsolateral prefrontal cortex (DLPFC) and the cerebellum. Conjunction and contrast analyses highlighted that these two regions were specifically associated with Motor Imagery when compared to the other modalities of imagery examined, making them worthy of further discussion.

In our present study, the DLPFC was consistently recruited during Motor Imagery, but not during the other modalities of imagery examined. This is of note as the DLPFC plays an important role in executive functions such as working memory (Rodríguez-Nieto et al., 2022), which has been associated with the vividness of both Visual and Auditory Imagery (Baddeley & Andrade, 2000). However, neuroimaging evidence examining visual working memory and Visual Imagery indicates they share representations in more perceptual areas (i.e. visual cortex; Albers et al., 2013). This shared perceptual processing may therefore render the need for higher-level (DLPFC) activation unnecessary. By contrast, recruitment of the DLPFC during Motor Imagery is consistent with existing models. Motor Simulation Theory (Jeannerod, 2001) suggests that DLPFC is recruited for short-term storage of information supporting the simulated action. Similarly, the Motor-Cognitive model (Glover et al., 2020; Glover & Baran, 2017; Martel & Glover, 2023) proposes that DLPFC may contribute to a pool of central executive resources used during Motor Imagery in a similar fashion to working memory. The proposals of both these models are therefore in line with recent work highlighting the existence of ‘motor working memory’ (Hillman et al., 2024). While further work is required to identify the neural correlates of motor working memory, the recruitment of DLPFC is highly plausible in this context.

The cerebellum has classically been associated with motor control (Holmes, 1917), and plays a key role in error-based learning (Sokolov et al., 2017). More specifically, the cerebellum is considered the primary candidate site for ‘forward models’ used to predict the sensorimotor consequences of actions (Miall et al., 1993; Wolpert et al., 1998). Prior behavioural work indicates sensorimotor prediction also occurs during Motor Imagery (Kilteni et al., 2018), providing a plausible role for the cerebellum in Motor Imagery (Lebon, 2024; Miall, 2024; Rieger et al., 2023). More recent work has also indicated that the cerebellum is active in higher cognitive functions (McDougle et al., 2022). Studies in patients with ‘cerebellar cognitive affective syndrome’, resulting from damage or impairment of the cerebellum, link it with deficits in executive functioning, and more specifically with working memory (Beuriat et al., 2020; Hayter et al., 2007; Ramnani, 2006). In particular, patients with lesions in lobule VI (the site of the peak cerebellar coordinate in the present meta-analysis), have disorders of working memory, while other executive functions are unaffected (Beuriat et al., 2020). This is consistent with meta-analytic studies that have identified consistent recruitment of the cerebellum for tasks involving working memory, but not other domains of executive function (Rodríguez-Nieto et al., 2022). This again re-iterates the association between Motor Imagery and working memory as discussed above in relation to the DLPFC. These differing motor and cognitive roles of the cerebellum could be linked to the differing models of Motor Imagery; the ‘motoric’ account of cerebellar activity during Motor Imagery would be more in line with Motor Simulation Theory, while the potential role of the cerebellum in working memory would be more consistent with the account of the Motor-Cognitive model.

### 4.3 Visual Imagery

In contrast to Winlove et al. (2018) but in agreement with Spagna et al. (2021), our meta-analysis of Visual Imagery did not reveal activation of V1. This non-activation is consistent with behavioural studies of patients with lesions in visual areas. Moro et al. (2008) found that Visual Imagery deficits were present in patients where V1 was intact, complementing studies indicating that occipital damage is neither necessary nor sufficient to produce deficits in visual imagery (Bartolomeo, 2002). Rather, these studies propose that the temporal lobe plays a crucial role in Visual Imagery. Indeed, a lesion in the left temporal lobe can lead to Visual Imagery deficits, especially for the mental generation of object form or colour, while Visual Imagery of faces more often results of a bilateral damage (Bartolomeo, 2002). Neuropsychological work has also highlighted the importance of the temporal lobe in the perceptual identification of objects (Goodale & Milner, 1992). These results are therefore consistent with the temporal activation in the present meta-analysis.

A notable result was that during Visual Imagery, our analysis revealed activation at the pre-SMA level which had not been shown in the two previous meta-analyses (Spagna et al., 2021; Winlove et al., 2018). Relatively few papers on Visual Imagery have considered the role of the pre-SMA in Visual Imagery, though work by Kosslyn & Thompson (2003) mention the region relating to the pre-SMA in the areas of activation during a Visual Imagery task. Although this area of the brain has often been associated with movement, its activation during Visual Imagery is not surprising if we consider its potential functions. For example, a study by Tanaka et al. (2005) showed that Brodman’s area 6, which corresponds to pre-SMA, plays a critical role in spatial representations, which is a possible aspect of Visual Imagery.

### 4.4 Auditory Imagery

The largest activation cluster observed in the meta-analysis of Auditory Imagery was in supplementary motor cortex, spanning the pre-SMA and SMA proper. This activation is not surprising given the studies carried out by Lima et al. (2016) on the association between Auditory Imagery and activation of the pre-SMA and SMA proper. This activation is thought to be due to their role in generating and controlling the motor programmes associated with auditory perception. These brain regions are involved in the coordination of movements and actions, including those linked to vocal production and the manipulation of musical instruments. During Auditory Imagery, SMA and pre-SMA are involved in retrieving and exploiting sensory expectations based on potential motor actions, thereby optimising perceptual processes, and generating a subjective experience of ’listening’.

Similarly to the SMA, we observed activation of the ventral and dorsal premotor cortices. In a study carried out in musicians (Schürmann et al., 2002), this activation was associated with the visualization of notes before imagining the associated sound. Similar activation is also observed in auditory verbal imagery studies (McGuire et al., 1996) where voices/sounds are often associated with actions. Compared to imagining your own voice, imagining someone else’s voice is proposed to require more covert articulation, engagement of auditory attention, and verbal control.

Recruitment of the left inferior parietal lobule during auditory imagery is consistent with previous reviews on Auditory Imagery (Hubbard, 2010; Kosslyn et al., 2001). This area is activated in both voluntary Auditory Imagery and when hearing auditory hallucinations (Shergill et al., 2001). Although this area is not part of the auditory cortex as such, it is as much activated when imagining sounds as when actually perceiving them (Zatorre et al., 1996).

As noted previously, Auditory Imagery was unique in recruiting the associated ‘primary’ cortex (i.e. while Auditory Imagery recruited A1, Motor Imagery did not appear to directly recruit M1, nor did Visual Imagery seem to directly recruit V1). We note, however, that the processing in A1 could also be considered more associative than for other equivalent regions. For example, King & Nelken (2009) note that while neurons in V1 show a preferential response for highly specific stimuli, to date no analogous response to stimuli has been found in A1, which instead appear to respond to a varied range of stimuli. This broad processing has led to the proposal that A1 may be functionally analogous to higher areas of other sensory systems (King & Nelken, 2009). This result is therefore broadly consistent with the general observation that Mental Imagery appears to preferentially involve regions involved in higher order, associative processing.

Notably, most of the studies used in the present meta-analysis used fMRI scanning procedures. The MRI scanner could be considered a challenging environment to study auditory processing due to the inherent background noises associated with scanning sequences. While the traditional contrast analysis approach should in theory minimize activations related to background noise, it could be argued that subtle differences in this background noise could result in activation of A1. We note, however, that if this were the case, we might expect A1 to be recruited in all of the present meta-analyses, which was not the case. Furthermore, work using PET (which uses a procedure comparable to MRI, but without the associated environmental noise) has implicated the involvement of A1, but not V1, in Auditory Imagery (for a review see Zvyagintsev et al., 2013). We therefore conclude that the recruitment of A1 in the present study is unlikely to represent an artefact of the noise associated with the scanner environment, and instead is more likely to represent true consistent recruitment of A1 during Auditory Imagery itself.

### 4.5 Consistent sub-network for Motor, Visual and Auditory Imagery

A conjunction across Motor, Visual, and Auditory Imagery identified a left-lateralized network of brain areas including the pre-SMA, ventral premotor cortex, and inferior parietal lobule. A common feature of these regions is their implication in broadly defined networks for attention, spatial processing, and decision-making (Bouchard et al., 2023). The pre-SMA is specifically recruited when participants focus their attention on their intentions (Lau et al., 2004); the consistent recruitment of the pre-SMA across multiple modalities of imagery could therefore arise from focussing attention on the intentional generation of an imagined experience. The IPL is implicated in attention related to spatial processing, including spatial, visuospatial, and motor attention (Corbetta et al., 2008; Corbetta & Shulman, 2002, Kraeutner et al., 2016) as well as lexical decisions (Numssen et al., 2021). By comparison, PMv is also implicated in spatial perception (Rizzolatti et al., 2002), while also playing a key role in decision making, processing sensory information used to choose, decide, act, and evaluate the results of previous decisions (Acuña et al., 2010). IPL and PMv may therefore support the generation of the mental image in space relative to the individual (e.g. spatial kinematics of imagined movements, spatial aspects of imagined visual scenes, spatial location of imagined sounds), with PMv involved in a more executive role.

The existence of a neural network that is common to all three imagery modalities could potentially explain the results of recent work by Dawes et al. (2024) on multisensory subtypes of aphantasia. This study demonstrated a range of different types of aphantasia; some aphantasics only had problems during visual imagery, others had problems with all imagery modalities except one, while those with the most severe form of aphantasia were unable to imagine all modalities of imagery. Differences in specific forms of imagery could be explained by deficits in the specific brain networks identified for those modalities in the present paper, with the possibility that multimodal imagery deficits are linked to the ‘core’ network.

### 4.6 Strengths and limitations

Individual neuroimaging studies typically include a small sample size of 12–20 participants, resulting in limited statistical power. To address this issue, meta-analytic techniques aggregate data across multiple studies, allowing for the examination of data from thousands of participants. Unlike review papers, which offer subjective evaluations, meta-analysis employs statistical methods to provide an objective summary of research findings. Nonetheless, meta-analyses are constrained by the existing literature, which may limit their scope, and they encounter various limitations such as age, sex, and task-related effects. Also, only papers from the PubMed database were collected. Searches in other databases could have provided more papers but only using PubMed is a standard in ALE meta-analyses.

Our main objective was therefore to compare all modalities of Imagery. Given the insufficient number of studies in three (Tactile, Gustatory, Olfactory) of the six modalities, we were unable to carry out the desired analyses. We hope that these areas will be studied more in the future to allow a more complete and detailed analysis of Imagery as a whole. However, we can still observe a link between the three modalities of Imagery (Auditory, Visual, Motor) that we were able to study, and future work will verify whether this pattern is consistent across other modalities of imagery.

Another limitation in the literature in general and that could explain the divergence in the results is the methodological differences observed. For example, in the field of Motor Imagery, it has already been pointed out that there is a recurrent lack of information about the methods used regarding the modality and the perspective of imagery (Van Caenegem et al., 2022). To remedy this, guidelines have been created to standardise methods in this field and avoid any confusion in future studies (Moreno-Verdú et al., 2024). This problem observed with Motor Imagery may be the same for Auditory and Visual Imagery and would therefore explain the divergence in the literature regarding the results observed. Also, this lack of information in the methods prevented certain sub-analyses that could have been carried out, such as Motor visual Imagery with Visual Imagery itself.

In the present manuscript Motor Imagery is broadly considered a combination of the use of visual-motor and kinesthetic-motor imagery. While we had hoped to perform separate analyses of these sub-modalities of Motor Imagery, unfortunately there were too few studies that examined them separately for us to run meaningful analyses. Extrapolating from the present results, we speculate that visual-motor imagery may be more likely to recruit visual areas similar to those identified in our present meta-analysis of Visual Imagery as well as regions involved in visuomotor integration (e.g. parietal cortex), whereas kinesthetic-motor imagery may be more closely linked to motor regions (e.g. premotor cortex). On a similar note, Auditory Imagery included studies where participants were asked to imagine hearing music. Previous work has suggested that hearing the sound of music can lead to the activation of motor programs associated with playing musical instruments (Schürmann et al., 2002). Again, unfortunately too few studies were present in the current manuscript to perform meaningful analyses of this sub-category of Auditory Imagery. Future work in this field may consider using different imagery tasks (e.g. imagining hearing a musical instrument compared to hearing sounds that are not directly associated with performing movements) to examine this question further.

In relation to this lack of homogeneity in the methods used for Motor Imagery studies, another bias that could be taken into consideration is the fact that the ability to imagine was assessed or not. In the analyses carried out, assessing the ability to imagine was not an inclusion or exclusion criterion. In the results provided, there may therefore be discrepancies between the data where the ability to imagine was assessed and those where it was not. Indeed, the ability of an individual to use Motor Imagery can significantly affect their effectiveness in achieving intended outcomes, highlighting the importance of assessing these abilities before interventions (Cumming & Ramsey, 2008; Martin et al., 1999).

## 5 Conclusion

In conclusion, the present results identify notable similarities between the brain regions involved in different modality of Mental Imagery. Consistent with previous models, we found that Mental Imagery generally recruited brain networks similar to those involved in the associated real-life experiences, with the exception that Mental Imagery appears to preferentially recruit ‘higher order’ associative brain regions as opposed to more ‘lower-level’ direct sensorimotor areas. Conjunction analyses also identified a common network including the pre-SMA, PMv, and IPL. We propose this ‘core’ network is associated with attentional, spatial, and decision-making aspects of Mental Imagery.

## Author Roles (CRediT)

| Author | Conceptualization | Methodology | Software | Validation | Formal Analysis | Investigation | Resources | Data Curation | Writing – original draft | Writing – reviewing & editing | Visualization | Supervision | Project Administration | Funding Acquisition |
| --- | --- | --- | --- | --- | --- | --- | --- | --- | --- | --- | --- | --- | --- | --- |
| Elise Van Caenegem | x | x |  | x | x | x | x | x | x | x | x | x | x | x |
| Robert Hardwick | x | x |  | x |  | x | x | x |  | x |  | x | x | x |
| Marcos Moreno-Verdu |  |  |  |  |  | x |  |  |  | x |  |  |  |  |
| Baptiste Waltzing |  |  |  |  |  | x |  |  |  | x |  |  |  |  |
| Siobhan McAteer |  |  |  |  |  | x |  |  |  |  |  |  |  |  |
| Gautier Hamoline |  |  |  |  |  | x |  |  |  |  |  |  |  |  |
| Frahm Lenart |  |  | x | x |  |  |  |  |  |  |  |  |  |  |

