## Supplementary Tables for "Multisensory approach in Mental Imagery: ALE meta-analyses comparing Motor, Visual and Auditory Imagery"

**Supplementary Materials**

**Table of Content**

### ***Information about supplementary tables***

In this file, you will find all the analyses carried out and the detailed results for each of these analyses. Firstly, you will find the main analyses that were the subject of the results in the article, as well as the exploratory data, which includes data on tactile, gustatory, and olfactory imagery. Each time one of these types of imagery is analysed, the table showing the results is shown in grey.

Also, for all the analyses that included a type of imaging, the number of experiments that were the subject of these analyses was indicated.

To help you understand the tables, here is an indication of the figures between brackets in the ‘Microanatomical location’ column. These correspond to Brodmann areas.

In addition to the contrast analyses, you will find the balanced contrasts. This means that the minimum requirement of 20 experiments to carry out analyses is not necessary as the threshold level is different and allows a comparison of a very large number of studies with a smaller one.

Also, in addition to the analyses grouping all the articles collected; in order to avoid any bias linked to comparisons between subjects on different tasks, we carried out additional analyses of all the papers comparing only the imaging group and the resting baseline of these people, as well as those that made a comparison between the imaging and a control condition that was close to a resting baseline but not entirely so. These additional analyses enabled us to see whether the comparisons that could have been made in other papers did not remove areas of the brain that might have been of interest to us because of the contrast analyses that could have been carried out. As the results of the control analyses did not reveal any new areas of activation, only the principal results were analysed in greater depth in the paper.

Also, if you don't find an analysis or a conjunction, or a contrast between two types of imagery, or a sub-analysis, this means that there are no voxels activated for the analysis in question.

### Section 1: Main analyses (all)

#### Table S1: Meta-analysis of All Imagery combined (n=439)

| **Cluster Voxels** | **Z-score** | **Macroanatomical Location** | **Cytoarchitectonic/ Tractographic Label** | **MNI Coordinates** | | |
| --- | --- | --- | --- | --- | --- | --- |
|  |  |  |  | **x** | **y** | **z** |
| 9028 | 8.13 | L Temporal Pole (48) |  | -52 | 8 | 0 |
|  | 7.97 | L Insula Lobe (48) |  | -34 | 20 | 4 |
|  | 5.69 | L IFG (p. Triangularis) (45) |  | -44 | 28 | 26 |
|  | 4.84 | L Middle Frontal Gyrus (DLPFC) |  | -34 | 46 | 26 |
|  | 4.60 | L ACC (32) |  | -6 | 26 | 36 |
|  | 4.32 | L IFG (p. Triangularis) (45) |  | -42 | 36 | 16 |
|  | 3.92 | L IFG (p. Orbitalis) (47) |  | -44 | 42 | -4 |
|  | 3.67 | L Middle Frontal Gyrus (45) |  | -40 | 46 | 8 |
| 3956 | 8.13 | N/A (40) | L Area hIP1 (IPS) | -34 | -50 | 42 |
|  | 6.78 | L Inferior Parietal Lobule (40) | L Area PFt (IPL) | -52 | -32 | 42 |
|  | 6.48 | L Superior Temporal Gyrus (48) | L Area PFcm (IPL) | -54 | -36 | 24 |
|  | 5.61 | L SupraMarginal Gyrus (48) |  | -52 | -38 | 30 |
|  | 5.20 | L Postcentral Gyrus (48) | L Area PFop (IPL) | -62 | -26 | 30 |
| 3325 | 8.13 | R Precentral Gyrus (PMd) |  | 54 | 2 | 44 |
|  | 7.90 | R Putamen |  | 24 | 4 | 4 |
|  | 7.82 | R Insula Lobe (48) |  | 36 | 20 | 4 |
|  | 7.63 | R IFG (p. Opercularis) (PMv) | R Area 44 | 56 | 10 | 22 |
|  | 7.52 | R IFG (p. Opercularis) (PMv) | R Area 44 | 56 | 12 | 12 |
|  | 7.28 | R IFG (p. Opercularis) (PMv) |  | 54 | 8 | 36 |
|  | 7.12 | R IFG (p. Opercularis) (48) |  | 50 | 10 | 4 |
| 2118 | 8.13 | N/A (40) | R Area hIP2 (IPS) | 42 | -40 | 44 |
|  | 7.49 | R Superior Parietal Lobule (7) |  | 18 | -62 | 56 |
|  | 6.02 | R Inferior Parietal Lobule (7) | R Area hIP3 (IPS) | 32 | -54 | 46 |
|  | 5.71 | R Inferior Parietal Lobule (40) | R Area hIP3 (IPS) | 36 | -46 | 52 |
| 1101 | 8.08 | N/A | L Lobule VI (Hem) | -32 | -56 | -32 |
|  | 5.02 | L Inferior Occipital Gyrus (37) |  | -48 | -70 | 2 |
|  | 4.90 | L Inferior Temporal Gyrus (37) |  | -52 | -60 | -6 |
|  | 4.45 | L Fusiform Gyrus (37) | L Area FG4 | -42 | -54 | -16 |
|  | 4.39 | L Fusiform Gyrus (37) | L Area FG4 | -44 | -58 | -16 |
|  | 3.92 | L Inferior Temporal Gyrus (37) | L Area FG2 | -50 | -66 | -16 |
| 668 | 8.13 | R Cerebelum (Crus 1) | R Lobule VIIa crusI (Hem) | 36 | -56 | -32 |

#### Table S2: Meta-analysis of Motor Imagery (n=284)

| **Cluster Voxels** | **Z-score** | **Macroanatomical Location** | **Cytoarchitectonic/ Tractographic Label** | **MNI Coordinates** | | |
| --- | --- | --- | --- | --- | --- | --- |
|  |  |  |  | **x** | **y** | **z** |
| 10550 | 8.13 | L Temporal Pole (48) |  | -52 | 8 | 0 |
|  | 7.81 | R IFG (p. Opercularis) (PMv) | R Area 44 | 56 | 12 | 12 |
|  | 7.25 | R IFG (p. Opercularis) (PMv) | R Area 44 | 56 | 10 | 20 |
|  | 7.18 | L Precentral Gyrus (PMd) |  | -44 | -4 | 44 |
|  | 7.05 | R IFG (p. Opercularis) (PMv) |  | 54 | 8 | 36 |
|  | 7.01 | R Precentral Gyrus (PMd) |  | 52 | 2 | 46 |
|  | 6.94 | R Insula Lobe (48) |  | 36 | 20 | 4 |
|  | 6.39 | R Putamen |  | 26 | 0 | 2 |
|  | 6.12 | L Insula Lobe (48) |  | -34 | 16 | 6 |
|  | 4.89 | R Putamen |  | 20 | 14 | 2 |
|  | 4.16 | L IFG (p. Orbitalis) (47) |  | -32 | 20 | -8 |
| 3665 | 8.13 | N/A (40) |  | -36 | -42 | 42 |
|  | 6.85 | L Inferior Parietal Lobule (7) | L Area 7PC (SPL) | -28 | -54 | 60 |
|  | 6.76 | L Inferior Parietal Lobule (40) | L Area PFt (IPL) | -52 | -32 | 42 |
|  | 6.31 | L Superior Temporal Gyrus (48) | L Area PFcm (IPL) | -54 | -36 | 24 |
|  | 5.34 | L SupraMarginal Gyrus (2) | L Area PFop (IPL) | -62 | -28 | 32 |
| 1832 | 8.13 | R SupraMarginal Gyrus (40) | R Area 2 | 42 | -38 | 46 |
|  | 6.71 | R Superior Parietal Lobule (7) | R Area 7A (SPL) | 18 | -62 | 62 |
|  | 5.93 | R Inferior Parietal Lobule (40) | R Area hIP3 (IPS) | 36 | -46 | 52 |
|  | 4.79 | R Inferior Parietal Lobule (40) | R Area hIP3 (IPS) | 34 | -52 | 46 |
|  | 4.75 | R Precuneus (7) |  | 14 | -68 | 50 |
|  | 4.56 | R Angular Gyrus (7) |  | 28 | -62 | 50 |
|  | 4.21 | R Postcentral Gyrus (S1) | R Area 2 | 50 | -30 | 52 |
|  | 4.20 | R Postcentral Gyrus (S1) | R Area 2 | 48 | -32 | 56 |
| 674 | 8.13 | R Cerebelum (VI) | Lobule VIIa crusI (Hem) | 36 | -58 | -30 |
| 556 | 7.57 | L Cerebelum (VI) | L Lobule VI (Hem) | -30 | -60 | -26 |
|  | 7.28 | L Cerebelum (VI) | L Lobule VI (Hem) | -32 | -58 | -30 |
| 383 | 5.04 | L Middle Frontal Gyrus (DLPFC) |  | -38 | 40 | 22 |
|  | 4.07 | L Middle Frontal Gyrus (45) |  | -40 | 46 | 6 |
|  | 3.96 | L IFG (p. Triangularis) (45) |  | -42 | 30 | 26 |
|  | 3.95 | L IFG (p. Triangularis) (45) |  | -42 | 38 | 14 |
|  | 3.72 | L IFG (p. Orbitalis) (47) |  | -42 | 36 | 0 |

#### Table S3: Meta-analysis of Visual Imagery (n=92)

| **Cluster Voxels** | **Z-score** | **Macroanatomical Location** | **Cytoarchitectonic/ Tractographic Label** | **MNI Coordinates** | | |
| --- | --- | --- | --- | --- | --- | --- |
|  |  |  |  | **x** | **y** | **z** |
| 801 | 5.37 | L Superior Parietal Lobule (7) |  | -20 | -68 | 52 |
|  | 5.32 | L Inferior Parietal Lobule (7) | L Area hIP3 (IPS) | -40 | -42 | 46 |
|  | 4.34 | L Inferior Parietal Lobule (7) | L Area hIP3 (IPS) | -32 | -58 | 46 |
|  | 3.88 | L Inferior Parietal Lobule (40) | L Area hIP2 (IPS) | -52 | -44 | 46 |
| 559 | 5.65 | L Inferior Temporal Gyrus (37) |  | -52 | -58 | -8 |
|  | 4.92 | N/A (37) | L Area FG2 | -48 | -68 | -14 |
|  | 3.83 | L Fusiform Gyrus (37) | L Area FG4 | -42 | -52 | -16 |
| 416 | 5.71 | L IFG (p. Opercularis) (PMv) | L Area 44 | -46 | 6 | 28 |
| 325 | 5.31 | L Precentral Gyrus (PMd) |  | -28 | -4 | 52 |
| 255 | 6.21 | R Middle Frontal Gyrus (PMd) |  | 28 | -4 | 58 |
| 231 | 4.32 | L Posterior-Medial Frontal (preSMA) |  | -2 | 4 | 58 |
|  | 4.03 | L Superior Medial Gyrus (preSMA) |  | 2 | 22 | 44 |
|  | 3.84 | L Superior Medial Gyrus (preSMA) |  | -4 | 18 | 48 |
|  | 3.83 | L Superior Medial Gyrus (preSMA) |  | -2 | 20 | 46 |
| 227 | 4.44 | N/A (7) |  | 20 | -64 | 54 |
|  | 4.26 | R Superior Occipital Gyrus (7) |  | 24 | -72 | 40 |
| 161 | 5.21 | R Supramarginal Gyrus (40) | R Area 2 | 44 | -38 | 46 |
| 159 | 5.07 | R Inferior Temporal Gyrus (37) |  | 50 | -56 | -12 |

#### Table S4: Meta-analysis of Auditory Imagery (n=48)

| **Cluster Voxels** | **Z-score** | **Macroanatomical Location** | **Cytoarchitectonic/ Tractographic Label** | **MNI Coordinates** | | |
| --- | --- | --- | --- | --- | --- | --- |
|  |  |  |  | **x** | **y** | **z** |
| 873 | 6.24 | L Posterior-Medial Frontal (pre-SMA) |  | -4 | 2 | 66 |
|  | 5.77 | L Posterior-Medial Frontal (pre-SMA) |  | 0 | 10 | 58 |
|  | 4.61 | L MCC (SMA) |  | 2 | 16 | 42 |
|  | 4.47 | L MCC (24) |  | -2 | 20 | 38 |
|  | 4.03 | L MCC (SMA) |  | -6 | 10 | 44 |
|  | 3.51 | L Posterior-Medial Frontal (pre-SMA) |  | 0 | 20 | 62 |
| 766 | 6.25 | L IFG (p. Opercularis) (PMv) | L Area 44 | -50 | 10 | 20 |
|  | 5.19 | L Temporal Pole (38) | L Area TE 3 | -54 | 6 | -6 |
|  | 5.08 | L IFG (p. Orbitalis) (47) |  | -40 | 20 | 0 |
|  | 5.06 | L Insula Lobe (48) |  | -34 | 20 | 4 |
|  | 4.04 | N/A (45) | L Area 45 | -58 | 24 | 6 |
|  | 3.27 | L IFG (p. Triangularis) (S1) | L Area 44 | -50 | 12 | 6 |
| 215 | 5.03 | L Superior Temporal Gyrus (A1-42) |  | -60 | -42 | 24 |
| 212 | 7.15 | R Precentral Gyrus (PMd) |  | 54 | 2 | 46 |
| 175 | 6.75 | L Precentral Gyrus (PMd) |  | -50 | -2 | 50 |
| 153 | 4.61 | L Inferior Parietal Lobule (7) |  | -32 | -54 | 52 |
|  | 4.47 | L Inferior Parietal Lobule (40) |  | -32 | -52 | 44 |
|  | 4.09 | N/A (40) | L Area hIP1 (IPS) | -36 | -48 | 42 |
| 120 | 4.42 | R Middle Temporal Gyrus (21) |  | 52 | -32 | 2 |
|  | 3.72 | R Superior Temporal Gyrus (A1-42) |  | 50 | -38 | 10 |

#### Table S5: Meta-analysis of Tactile Imagery (n=9)

| **Cluster Voxels** | **Z-score** | **Macroanatomical Location** | **Cytoarchitectonic/ Tractographic Label** | **MNI Coordinates** | | |
| --- | --- | --- | --- | --- | --- | --- |
|  |  |  |  | **x** | **y** | **z** |
| 114 | 4.85 | N/A (40) | L Area hIP1 (IPS) | -34 | -46 | 42 |
| 100 | 5.28 | L Posterior-Medial Frontal (pre-SMA) |  | -8 | 12 | 56 |

#### Table S6: Meta-analysis of Gustatory Imagery (n=7)

| **Cluster Voxels** | **Z-score** | **Macroanatomical Location** | **Cytoarchitectonic/ Tractographic Label** | **MNI Coordinates** | | |
| --- | --- | --- | --- | --- | --- | --- |
|  |  |  |  | **x** | **y** | **z** |
| 230 | 5.57 | L Insula lobe (48) |  | -38 | -4 | 12 |

#### Table S7: Meta-analysis of All Imagery but not MI (n=155)

| **Cluster Voxels** | **Z-score** | **Macroanatomical Location** | **Cytoarchitectonic/ Tractographic Label** | **MNI Coordinates** | | |
| --- | --- | --- | --- | --- | --- | --- |
|  |  |  |  | **x** | **y** | **z** |
| 1180 | 6.96 | L Posterior-Medial Frontal (pre-SMA) |  | -6 | 8 | 58 |
|  | 6.55 | L Posterior-Medial Frontal (pre-SMA) |  | -4 | 2 | 66 |
|  | 5.17 | R MCC (SMA) |  | 2 | 20 | 42 |
|  | 5.05 | L Superior Medial Gyrus (pre-SMA) |  | -4 | 20 | 46 |
|  | 3.84 | L MCC (M1) |  | -6 | 8 | 44 |
|  | 3.71 | L ACC (24) |  | -4 | 28 | 32 |
| 1076 | 7.34 | L IFG (p. Opercularis) (PMv) | L Area 44 | -52 | 10 | 20 |
|  | 5.85 | L Precentral Gyrus (PMd) |  | -50 | -2 | 50 |
|  | 4.30 | L IFG (p. Triangularis) (45) |  | -44 | 28 | 26 |
| 715 | 6.15 | L Inferior Parietal Lobule (40) | L Area hIP1 (IPS) | -36 | -46 | 44 |
|  | 5.23 | L Superior Parietal Lobule (18) |  | -22 | -68 | 50 |
| 424 | 8.13 | L Inferior Temporal Gyrus (37) |  | -54 | -60 | -6 |
|  | 7.49 | L Inferior Occipital Gyrus (37) | L Area FG2 | -48 | -68 | -12 |
| 418 | 6.70 | L IFG (p. Triangularis) (48) |  | -34 | 22 | 4 |
|  | 4.01 | L Temporal Pole (38) |  | -52 | 6 | -6 |
|  | 3.78 | L IFG (p. Orbitalis) (48) |  | -46 | 12 | -6 |
|  | 3.38 | N/A (38) | L Area FG4 | -52 | 22 | -6 |
| 346 | 5.69 | R Middle Frontal Gyrus (PMd) |  | 30 | -2 | 54 |
|  | 5.59 | R Precentral Gyrus (PMd) |  | 54 | 2 | 46 |
|  | 5.53 | R Precentral Gyrus (PMd) |  | 52 | 4 | 48 |
| 321 | 4.75 | R Insula Lobe (48) |  | 48 | 10 | 0 |
|  | 3.86 | N/A (48) |  | 56 | 14 | 2 |
|  | 3.72 | R Insula Lobe (48) |  | 36 | 20 | 4 |
| 249 | 5.09 | L Precentral Gyrus (PMd) |  | -28 | -4 | 52 |
| 217 | 5.17 | R SupraMarginal Gyrus (40) | R Area hIP2 (IPS) | 44 | -40 | 46 |
| 154 | 5.14 | N/A (7) |  | 18 | -64 | 54 |
|  | 3.27 | R Precuneus (7) |  | 12 | -68 | 42 |

### Section 2: Conjunction analyses (all)

#### Table S8: Meta-analysis of Conjunction between Auditory and Visual Imagery

| **Cluster Voxels** | **Z-score** | **Macroanatomical Location** | **Cytoarchitectonic/ Tractographic Label** | **MNI Coordinates** | | |
| --- | --- | --- | --- | --- | --- | --- |
|  |  |  |  | **x** | **y** | **z** |
| 116 | 4.32 | L Posterior-Medial Frontal (pre-SMA) |  | -2 | 4 | 58 |
| 82 | 4.83 | L IFG (p. Opercularis) (PMv) |  | -48 | 10 | 24 |
| 62 | 3.99 | L Inferior Parietal Lobule (7) | L Area hIP1 (IPS) | -32 | -54 | 46 |
|  | 3.71 | L Inferior Parietal Lobule (40) | L Area hIP1 (IPS) | -36 | -48 | 44 |
| 33 | 3.71 | L Superior Medial Gyrus (pre-SMA) |  | 2 | 20 | 44 |

#### Table S9: Meta-analysis of Conjunction between Motor and Auditory Imagery

| **Cluster Voxels** | **Z-score** | **Macroanatomical Location** | **Cytoarchitectonic/ Tractographic Label** | **MNI Coordinates** | | |
| --- | --- | --- | --- | --- | --- | --- |
|  |  |  |  | **x** | **y** | **z** |
| 778 | 6.24 | L Posterior-Medial Frontal (pre-SMA) |  | -4 | 2 | 66 |
|  | 5.77 | L Posterior-Medial Frontal (pre-SMA) |  | 0 | 10 | 58 |
|  | 4.61 | L MCC (SMA) |  | 2 | 16 | 42 |
|  | 4.03 | L MCC (pre-SMA) |  | -6 | 10 | 44 |
| 496 | 5.99 | L IFG (p. Opercularis) (PMv) | L Area 44 | -52 | 10 | 18 |
|  | 5.06 | L Insula Lobe (48) |  | -34 | 20 | 4 |
|  | 4.70 | L Temporal Pole (38) |  | -52 | 8 | -4 |
|  | 3.64 | L IFG (p. Orbitalis) (48) |  | -40 | 16 | -4 |
|  | 3.27 | L IFG (p. Triangularis) (S1) | L Area 44 | -50 | 12 | 6 |
| 206 | 6.68 | R Precentral Gyrus (PMd) |  | 52 | 2 | 46 |
| 171 | 6.40 | L Precentral Gyrus (PMd) |  | -48 | -4 | 50 |
| 137 | 4.61 | L Inferior Parietal Lobule (7) | L Area hIP3 (IPS) | -32 | -54 | 52 |
|  | 4.47 | L Inferior Parietal Lobule (40) | L Area hIP1 (IPS) | -32 | -52 | 44 |
|  | 4.09 | N/A (40) | L Area hIP1 (IPS) | -36 | -48 | 42 |
| 78 | 4.47 | L Superior Temporal Gyrus (48) | L Area PFcm (IPL) | -54 | -40 | 24 |
|  | 4.28 | L Superior Temporal Gyrus | L Area PFcm (IPL) | -60 | -38 | 24 |

#### Table S10: Meta-analysis of Conjunction between Motor and Visual Imagery

| **Cluster Voxels** | **Z-score** | **Macroanatomical Location** | **Cytoarchitectonic/ Tractographic Label** | **MNI Coordinates** | | |
| --- | --- | --- | --- | --- | --- | --- |
|  |  |  |  | **x** | **y** | **z** |
| 604 | 5.37 | L Superior Parietal Lobule (7) |  | -20 | -68 | 52 |
|  | 5.32 | L Inferior Parietal Lobule (40) | L Area hIP3 (IPS) | -40 | -42 | 46 |
|  | 4.06 | L Inferior Parietal Lobule (7) | L Area hIP1 (IPS) | -34 | -54 | 46 |
|  | 3.51 | L Inferior Parietal Lobule (7) | L Area hIP3 (IPS) | -30 | -58 | 52 |
| 365 | 5.70 | L IFG (p. Opercularis) (PMv) | L Area 44 | -48 | 6 | 28 |
| 309 | 5.31 | L Precentral Gyrus (PMd) |  | -28 | -4 | 52 |
| 255 | 6.21 | R Middle Frontal Gyrus (PMd) |  | 28 | -4 | 58 |
| 207 | 4.32 | L Posterior-Medial Frontal (pre-SMA) |  | -2 | 4 | 58 |
|  | 4.03 | L Superior Medial Gyrus (pre-SMA) |  | 2 | 22 | 44 |
|  | 3.84 | L Superior Medial Gyrus (pre-SMA) |  | -4 | 18 | 48 |
|  | 3.83 | L Superior Medial Gyrus (pre-SMA) |  | -2 | 20 | 46 |
| 157 | 5.21 | R SupraMarginal Gyrus (40) | R Area 2 | 44 | -38 | 46 |
| 124 | 4.44 | N/A (7) |  | 20 | -64 | 54 |
|  | 3.56 | R Cuneus (7) |  | 20 | -70 | 44 |

#### Table S11: Meta-analysis of Conjunction between Auditory and Tactile Imagery

| **Cluster Voxels** | **Z-score** | **Macroanatomical Location** | **Cytoarchitectonic/ Tractographic Label** | **MNI Coordinates** | | |
| --- | --- | --- | --- | --- | --- | --- |
|  |  |  |  | **x** | **y** | **z** |
| 46 | 4.38 | L Posterior-Medial Frontal (pre-SMA) |  | -6 | 10 | 58 |
| 41 | 4.08 | N/A | L Area hIP1 (IPS) | -34 | -50 | 42 |

#### Table S12: Meta-analysis of Conjunction between Motor and Gustatory Imagery

| **Cluster Voxels** | **Z-score** | **Macroanatomical Location** | **Cytoarchitectonic/ Tractographic Label** | **MNI Coordinates** | | |
| --- | --- | --- | --- | --- | --- | --- |
|  |  |  |  | **x** | **y** | **z** |
| 37 | 4.07 | L Insula Lobe (PMd) |  | -42 | 2 | 8 |
| 21 | 3.99 | L Putamen (48) |  | -32 | -4 | 10 |

#### Table S13: Meta-analysis of Conjunction between Motor and Tactile Imagery

| **Cluster Voxels** | **Z-score** | **Macroanatomical Location** | **Cytoarchitectonic/ Tractographic Label** | **MNI Coordinates** | | |
| --- | --- | --- | --- | --- | --- | --- |
|  |  |  |  | **x** | **y** | **z** |
| 101 | 4.85 | N/A (40) | L Area hIP1 (IPS) | -34 | -46 | 42 |
| 98 | 5.28 | L Posterior-Medial Frontal (pre-SMA) |  | -8 | 12 | 56 |

#### Table S14: Meta-analysis of Conjunction between Visual and Tactile Imagery

| **Cluster Voxels** | **Z-score** | **Macroanatomical Location** | **Cytoarchitectonic/ Tractographic Label** | **MNI Coordinates** | | |
| --- | --- | --- | --- | --- | --- | --- |
|  |  |  |  | **x** | **y** | **z** |
| 34 | 3.94 | L Inferior Parietal Lobule (40) | L Area hIP1 (IPS) | -36 | -46 | 44 |
| 33 | 3.98 | L Posterior-Medial Frontal (pre-SMA) |  | -6 | 8 | 56 |

#### Table S15: Meta-analysis of Conjunction between Motor, Visual, Auditory Imagery

| **Cluster Voxels** | **Z-score** | **Macroanatomical Location** | **Cytoarchitectonic/ Tractographic Label** | **MNI Coordinates** | | |
| --- | --- | --- | --- | --- | --- | --- |
|  |  |  |  | **x** | **y** | **z** |
| 122 | 4.37 | L Posterior-Medial Frontal (pre-SMA) |  | -2 | 4 | 58 |
| 64 | 4.66 | L IFG (p. Opercularis) (PMv) | L Area 44 | -50 | 8 | 24 |
| 56 | 3.99 | L Inferior Parietal Lobule (7) | L Area hIP1 (IPS) | -32 | -54 | 46 |
|  | 3.71 | L Inferior Parietal Lobule (40) | L Area hIP1 (IPS) | -36 | -48 | 44 |
| 29 | 3.71 | L Superior Medial Gyrus (pre-SMA) |  | 2 | 20 | 44 |

#### Table S16: Meta-analysis of Conjunction between Motor, Visual and Tactile Imagery

| **Cluster Voxels** | **Z-score** | **Macroanatomical Location** | **Cytoarchitectonic/ Tractographic Label** | **MNI Coordinates** | | |
| --- | --- | --- | --- | --- | --- | --- |
|  |  |  |  | **x** | **y** | **z** |
| 34 | 3.94 | L Inferior Parietal Lobule (40) | L Area hIP1 (IPS) | -36 | -46 | 44 |
| 33 | 3.98 | L Posterior-Medial Frontal (pre-SMA) |  | -6 | 8 | 56 |

#### Table S17: Meta-analysis of Conjunction between Motor, Auditory and Tactile Imagery

| **Cluster Voxels** | **Z-score** | **Macroanatomical Location** | **Cytoarchitectonic/ Tractographic Label** | **MNI Coordinates** | | |
| --- | --- | --- | --- | --- | --- | --- |
|  |  |  |  | **x** | **y** | **z** |
| 46 | 4.38 | L Posterior-Medial Frontal (pre-SMA) |  | -6 | 10 | 58 |
| 41 | 4.08 | N/A | L Area hIP1 (IPS) | -34 | -50 | 42 |

#### Table S18: Meta-analysis of Conjunction between Visual, Auditory and Tactile Imagery

| **Cluster Voxels** | **Z-score** | **Macroanatomical Location** | **Cytoarchitectonic/ Tractographic Label** | **MNI Coordinates** | | |
| --- | --- | --- | --- | --- | --- | --- |
|  |  |  |  | **x** | **y** | **z** |
| 28 | 3.98 | L Posterior-Medial Frontal (pre-SMA) |  | -6 | 8 | 56 |
| 19 | 3.71 | L Inferior Parietal Lobule (40) | L Area hIP1 (IPS) | -36 | -48 | 44 |

#### Table S19: Meta-analysis of Conjunction between Motor, Visual, Auditory and Tactile Imagery

| **Cluster Voxels** | **Z-score** | **Macroanatomical Location** | **Cytoarchitectonic/ Tractographic Label** | **MNI Coordinates** | | |
| --- | --- | --- | --- | --- | --- | --- |
|  |  |  |  | **x** | **y** | **z** |
| 28 | 3.98 | L Posterior-Medial Frontal (pre-SMA) |  | -6 | 8 | 56 |
| 19 | 3.71 | L Inferior Parietal Lobule (40) | L Area hIP1 (IPS) | -36 | -48 | 44 |

### Section 3: Contrast results (all)

#### Table S20: Meta-analysis of Contrast_Auditory_greater_than_Visual

| **Cluster Voxels** | **Z-score** | **Macroanatomical Location** | **Cytoarchitectonic/ Tractographic Label** | **MNI Coordinates** | | |
| --- | --- | --- | --- | --- | --- | --- |
|  |  |  |  | **x** | **y** | **z** |
| 413 | 4.47 | L Posterior-Medial Frontal (6) |  | 0 | 8 | 64 |
| 378 | 3.54 | L Temporal Pole (38) | L Area TE 3 | -50 | 10 | 20 |
|  | 3.35 | L IFG (p. Triangularis) (48) | L Area 44 | -52 | 6 | -6 |
|  | 3.29 | L IFG (p. Triangularis) (48) |  | -40 | 20 | 0 |
|  | 3.01 | L IFG (p. Triangularis) (48) | L Area 44 | -34 | 20 | 4 |
|  | 2.49 | L IFG (p. Orbitalis) (48) | L Area 44 | -58 | 20 | 4 |
|  | 2.22 | L IFG (p. Opercularis) (6) | L Area 44 | -50 | 12 | 6 |
| 209 | 7.15 | R Precentral Gyrus (PMd) |  | 54 | 2 | 46 |
| 198 | 5.03 | L Superior Temporal Gyrus (A1-42) |  | -60 | -42 | 24 |
| 163 | 6.11 | L Precentral Gyrus (6) |  | -50 | -4 | 52 |
|  | 3.72 | L Precentral Gyrus (6) |  | -50 | -2 | 48 |
| 141 | 3.24 | L IFG (p. Orbitalis) (48) |  | -44 | 16 | 0 |
|  | 1.68 | L Insula Lobe (48) |  | -32 | 18 | 6 |
| 120 | 2.66 | L MCC (24) |  | -2 | 14 | 40 |
| 111 | 3.66 | R Middle Temporal Gyrus (21) |  | 50 | -30 | 0 |
|  | 3.09 | R Superior Temporal Gyrus (A1) |  | 54 | -38 | 12 |

#### Table S21: Meta-analysis of Contrast_Auditory_greater_than_Motor

| **Cluster Voxels** | **Z-score** | **Macroanatomical Location** | **Cytoarchitectonic/ Tractographic Label** | **MNI Coordinates** | | |
| --- | --- | --- | --- | --- | --- | --- |
|  |  |  |  | **x** | **y** | **z** |
| 873 | 6.24 | L Posterior-Medial Frontal (pre-SMA) |  | -4 | 2 | 66 |
|  | 5.77 | L Posterior-Medial Frontal (pre-SMA) |  | 0 | 10 | 58 |
|  | 4.61 | L MCC (SMA) |  | 2 | 16 | 42 |
|  | 4.47 | L MCC (24) |  | -2 | 20 | 38 |
|  | 4.03 | L MCC (pre-SMA) |  | -6 | 10 | 44 |
|  | 3.51 | L Posterior-Medial Frontal (pre-SMA) |  | 0 | 20 | 62 |
| 764 | 6.25 | L IFG (p. Opercularis) (PMv) | L Area 44 | -50 | 10 | 20 |
|  | 5.09 | L Temporal Pole (38) |  | -52 | 6 | -6 |
|  | 5.08 | L IFG (p. Orbitalis) (47) |  | -40 | 20 | 0 |
|  | 5.06 | L Insula Lobe (48) |  | -34 | 20 | 4 |
|  | 3.54 | N/A (48) | L Area 45 | -58 | 20 | 4 |
|  | 3.27 | L IFG (p. Triangularis) (S1) | L Area 44 | -50 | 12 | 6 |
| 215 | 5.03 | L Superior Temporal Gyrus (A1-42) |  | -60 | -42 | 24 |
| 212 | 7.15 | R Precentral Gyrus (PMd) |  | 54 | 2 | 46 |
| 175 | 6.75 | L Precentral Gyrus (PMd) |  | -50 | -2 | 50 |
| 153 | 4.61 | L Inferior Parietal Lobule (7) |  | -32 | -54 | 52 |
|  | 4.47 | L Inferior Parietal Lobule (40) |  | -32 | -52 | 44 |
|  | 4.08 | N/A (40) |  | -36 | -48 | 42 |

#### Table S22: Meta-analysis of Contrast_Motor_greater_than_Auditory

| **Cluster Voxels** | **Z-score** | **Macroanatomical Location** | **Cytoarchitectonic/ Tractographic Label** | **MNI Coordinates** | | |
| --- | --- | --- | --- | --- | --- | --- |
|  |  |  |  | **x** | **y** | **z** |
| 2007 | 8.13 | N/A (PMd) |  | -22 | -10 | 50 |
|  | 3.72 | N/A (PMd) |  | -30 | -8 | 46 |
|  | 3.54 | L Posterior-Medial Frontal (SMA) |  | -4 | -12 | 68 |
|  | 3.18 | R Posterior-Medial Frontal (SMA) |  | 4 | -14 | 64 |
|  | 2.95 | R Posterior-Medial Frontal (SMA) |  | 8 | -12 | 62 |
|  | 2.88 | R Posterior-Medial Frontal (SMA) |  | 8 | -16 | 68 |
| 1939 | 8.13 | L Inferior Parietal Lobule (40) | L Area PFt (IPL) | -40 | -36 | 46 |
|  | 7.84 | L Superior Parietal Lobule (7) | L Area 7A (SPL) | -18 | -62 | 58 |
|  | 3.72 | L Inferior Parietal Lobule (40) | L Area PFt (IPL) | -54 | -34 | 50 |
|  | 3.19 | L Superior Parietal Lobule (7) | L Area 5L (SPL) | -30 | -50 | 66 |
|  | 2.97 | L SupraMarginal Gyrus (2) | L Area PFt (IPL) | -60 | -26 | 46 |
| 433 | 3.29 | R Precentral Gyrus (PMd) |  | 30 | -10 | 62 |
|  | 3.24 | R Precentral Gyrus (SMA) |  | 36 | -10 | 60 |
|  | 3.04 | N/A (PMd) |  | 20 | -8 | 58 |
|  | 2.89 | N/A (PMd) |  | 20 | -10 | 54 |
|  | 2.42 | N/A (PMd) |  | 34 | -4 | 46 |
|  | 2.30 | R Middle Frontal Gyrus (PMd) |  | 36 | -2 | 48 |
| 352 | 3.35 | R Postcentral Gyrus (S1) | R Area 3b | 42 | -32 | 58 |
|  | 3.17 | R Postcentral Gyrus (S1) | R Area 4p | 38 | -34 | 58 |
|  | 3.16 | R Postcentral Gyrus (S1) | R Area 4p | 34 | -34 | 56 |
|  | 3.12 | R Postcentral Gyrus (S1) | R Area 4p | 36 | -30 | 56 |
|  | 2.21 | R Postcentral Gyrus (2) | R Area 2 | 32 | -44 | 60 |
| 274 | 3.04 | R Superior Parietal Lobule | R Area 7A (SPL) | 22 | -64 | 66 |
|  | 2.91 | R Superior Parietal Lobule | R Area hIP3 (IPS) | 26 | -62 | 60 |
|  | 2.89 | R Superior Parietal Lobule | R Area 7A (SPL) | 16 | -62 | 68 |
|  | 2.67 | R Superior Parietal Lobule (5) |  | 16 | -60 | 62 |
|  | 2.64 | R Precuneus (5) | R Area 5L (SPL) | 10 | -58 | 64 |
| 171 | 2.89 | L Putamen |  | -26 | -8 | 12 |
| 162 | 2.85 | L IFG (p. Opercularis) (PMv) | L Area 44 | -58 | 8 | 30 |
| 88 | 2.49 | R IFG (p. Opercularis) (PMv) | R Area 44 | 56 | 4 | 32 |
| 78 | 2.56 | L Cerebelum (VI) | L Lobule VI (Hem) | -22 | -64 | -22 |
| 47 | 2.41 | L IFG (p. Orbitalis) (47) |  | -40 | 38 | 2 |
| 31 | 2.10 | L Temporal Pole (PMv) |  | -56 | 6 | 4 |
| 27 | 2.08 | R Putamen (48) |  | 30 | -4 | -2 |
|  | 1.98 | N/A (48) |  | 26 | -4 | -4 |

#### Table S23: Meta-analysis of Contrast_Motor_greater_than_Visual

| **Cluster Voxels** | **Z-score** | **Macroanatomical Location** | **Cytoarchitectonic/ Tractographic Label** | **MNI Coordinates** | | |
| --- | --- | --- | --- | --- | --- | --- |
|  |  |  |  | **x** | **y** | **z** |
| 3332 | 8.13 | L Temporal Pole (PMv) |  | -52 | 6 | 2 |
|  | 7.05 | L Superior Frontal Gyrus (PMd) |  | -20 | 0 | 62 |
|  | 6.92 | N/A (PMd) |  | -16 | -4 | 56 |
|  | 5.84 | L Precentral Gyrus (M1) |  | -46 | -8 | 54 |
|  | 3.72 | L IFG (p. Opercularis) (S1) | L Area 44 | -60 | 12 | 10 |
|  | 3.54 | L Precentral Gyrus (M1) |  | -52 | -8 | 48 |
|  | 3.29 | L MCC (pre-SMA) |  | -6 | 10 | 48 |
|  | 3.24 | L MCC (pre-SMA) |  | -2 | 10 | 48 |
|  | 3.04 | L Precentral Gyrus (PMd) |  | -38 | -4 | 64 |
|  | 2.93 | L Precentral Gyrus (PMd) |  | -34 | -2 | 66 |
|  | 2.88 | L Precentral Gyrus (PMv) |  | -56 | -2 | 42 |
| 1450 | 8.13 | L Inferior Parietal Lobule (40) |  | -36 | -46 | 54 |
|  | 3.24 | L Inferior Parietal Lobule (S1) | L Area 2 | -56 | -28 | 46 |
|  | 3.15 | L Superior Temporal Gyrus (A1-41) | L Area OP1 [SII] | -52 | -32 | 18 |
|  | 3.09 | L SupraMarginal Gyrus | L Area PFt (IPL) | -66 | -30 | 36 |
|  | 3.04 | L SupraMarginal Gyrus (40) | L Area PF (IPL) | -64 | -34 | 30 |
|  | 2.99 | L Postcentral Gyrus (M1) | L Area PFt (IPL) | -62 | -24 | 38 |
|  | 2.91 | L Superior Parietal Lobule (5) | L Area 5L (SPL) | -16 | -54 | 68 |
|  | 2.74 | L Superior Parietal Lobule | L Area 5L (SPL) | -24 | -54 | 66 |
|  | 2.64 | L Superior Temporal Gyrus (48) | L Area PFcm (IPL) | -56 | -40 | 28 |
|  | 2.34 | L Inferior Parietal Lobule (40) |  | -34 | -34 | 42 |
|  | 2.33 | N/A (40) |  | -32 | -36 | 40 |
| 444 | 8.13 | R Cerebelum (Crus 1) | R Lobule VI (Hem) | 34 | -58 | -28 |
|  | 3.72 | R Cerebelum (VI) | R Lobule VI (Hem) | 28 | -60 | -26 |
|  | 3.19 | N/A | R Lobule VI (Hem) | 32 | -46 | -34 |
| 424 | 3.24 | R Precentral Gyrus (PMd) |  | 48 | -4 | 44 |
|  | 3.24 | R Precentral Gyrus (PMd) |  | 48 | -6 | 48 |
|  | 3.16 | R Precentral Gyrus (PMd) |  | 52 | -4 | 46 |
|  | 3.06 | R Precentral Gyrus (M1) |  | 34 | -14 | 58 |
|  | 3.01 | R Precentral Gyrus (M1) |  | 36 | -12 | 60 |
|  | 2.82 | R Middle Frontal Gyrus (M1) |  | 38 | -8 | 60 |
|  | 2.59 | R Precentral Gyrus (M1) |  | 40 | -12 | 56 |
|  | 2.44 | R Middle Frontal Gyrus (PMd) |  | 40 | -2 | 62 |
|  | 2.17 | N/A (PMd) |  | 20 | -10 | 54 |
| 366 | 7.11 | L Cerebelum (VI) | L Lobule VI (Hem) | -28 | -60 | -24 |
|  | 3.72 | L Cerebelum (VI) | L Lobule VI (Hem) | -24 | -64 | -24 |
| 333 | 3.72 | R IFG (p. Opercularis) (PMv) | R Area 44 | 56 | 6 | 12 |
|  | 3.24 | R IFG (p. Opercularis) (SMA) | R Area 44 | 64 | 10 | 14 |
| 144 | 2.83 | L Middle Frontal Gyrus (DLPFC) |  | -34 | 40 | 24 |
| 132 | 2.46 | N/A (48) |  | 30 | 16 | 8 |
|  | 2.43 | R Putamen (48) |  | 26 | 16 | 4 |
|  | 1.85 | R Putamen (48) |  | 26 | 8 | 10 |
| 125 | 3.04 | R Postcentral Gyrus (3) | R Area 2 | 48 | -26 | 52 |
|  | 2.83 | R Postcentral Gyrus (3) | R Area 1 | 46 | -28 | 56 |
|  | 2.60 | R Postcentral Gyrus (3) | R Area 3b | 40 | -30 | 52 |
| 75 | 2.45 | L Putamen (48) |  | -32 | -6 | 6 |
|  | 1.85 | L Putamen |  | -26 | -10 | 10 |
| 64 | 2.99 | R Pallidum |  | 28 | -8 | 0 |
| 52 | 2.47 | R Superior Parietal Lobule (5) | R Area 7PC (SPL) | 22 | -56 | 62 |
| 50 | 2.22 | N/A (48) |  | -26 | 12 | 6 |
|  | 1.84 | L Putamen |  | -24 | 6 | 12 |
|  | 1.76 | L Insula Lobe (48) |  | -34 | 12 | 8 |

#### Table S24: Meta-analysis of Contrast_Visual_greater_than_Motor

| **Cluster Voxels** | **Z-score** | **Macroanatomical Location** | **Cytoarchitectonic/ Tractographic Label** | **MNI Coordinates** | | |
| --- | --- | --- | --- | --- | --- | --- |
|  |  |  |  | **x** | **y** | **z** |
| 801 | 5.37 | L Superior Parietal Lobule (7) |  | -20 | -68 | 52 |
|  | 5.32 | L Inferior Parietal Lobule (40) | L Area hIP3 (IPS) | -40 | -42 | 46 |
|  | 4.32 | L Inferior Parietal Lobule (7) | L Area hIP1 (IPS) | -32 | -56 | 46 |
|  | 3.75 | L Inferior Parietal Lobule (40) | L Area hIP2 (IPS) | -50 | -44 | 46 |
|  | 3.64 | L Superior Occipital Gyrus (7) |  | -20 | -68 | 42 |
| 416 | 5.71 | L IFG (p. Opercularis) (PMv) | L Area 44 | -46 | 6 | 28 |
| 325 | 3.78 | L Inferior Temporal Gyrus (37) |  | -52 | -62 | -2 |
|  | 3.72 | L Fusiform Gyrus (37) | L Area FG4 | -42 | -54 | -16 |
|  | 3.51 | L Fusiform Gyrus (37) | L Area FG4 | -38 | -52 | -18 |
|  | 3.43 | L Inferior Temporal Gyrus (37) |  | -56 | -62 | 0 |
|  | 3.35 | L Inferior Temporal Gyrus (37) |  | -48 | -62 | -4 |
|  | 3.19 | L Fusiform Gyrus (19) | L Area FG2 | -46 | -64 | -16 |
|  | 3.09 | L Inferior Occipital Gyrus (19) | L Area FG2 | -48 | -70 | -8 |
|  | 2.19 | L Inferior Temporal Gyrus (37) |  | -52 | -52 | -4 |
|  | 1.98 | L Inferior Temporal Gyrus (20) |  | -50 | -50 | -8 |
|  | 1.84 | L Fusiform Gyrus (19) | L Area FG2 | -42 | -70 | -14 |
| 324 | 5.31 | L Precentral Gyrus (PMd) |  | -28 | -4 | 52 |
| 255 | 6.21 | R Middle Frontal Gyrus (PMd) |  | 28 | -4 | 58 |
| 231 | 4.32 | L Posterior-Medial Frontal (pre-SMA) |  | -2 | 4 | 58 |
|  | 4.03 | L Superior Medial Gyrus (pre-SMA) |  | 2 | 22 | 44 |
|  | 3.84 | L Superior Medial Gyrus (pre-SMA) |  | -4 | 18 | 48 |
|  | 3.83 | L Superior Medial Gyrus (pre-SMA) |  | -2 | 20 | 46 |
| 214 | 4.44 | N/A (7) |  | 20 | -64 | 54 |
|  | 3.56 | R Cuneus (7) |  | 20 | -70 | 44 |
|  | 3.43 | R Superior Occipital Gyrus (7) |  | 24 | -68 | 42 |
|  | 2.56 | R Middle Occipital Gyrus (19) |  | 28 | -74 | 36 |
| 161 | 5.21 | R SupraMarginal Gyrus (40) | R Area 2 | 44 | -38 | 46 |
| 65 | 3.06 | R Inferior Temporal Gyrus (37) |  | 54 | -58 | -6 |
|  | 2.71 | R Inferior Temporal Gyrus (37) |  | 52 | -62 | -8 |
|  | 2.15 | R Inferior Temporal Gyrus (20) | R Area FG4 | 48 | -52 | -16 |
|  | 2.14 | R Inferior Temporal Gyrus (37) | R Area FG4 | 46 | -52 | -12 |
|  | 2.07 | R Inferior Temporal Gyrus (37) | R Area FG4 | 48 | -60 | -12 |

#### Table S25: Meta-analysis of Conjunction Motor greater than Visual and Auditory Imagery

| **Cluster Voxels** | **Z-score** | **Macroanatomical Location** | **Cytoarchitectonic/ Tractographic Label** | **MNI Coordinates** | | |
| --- | --- | --- | --- | --- | --- | --- |
|  |  |  |  | **x** | **y** | **z** |
| 1137 | 8.13 | L Posterior-Medial Frontal (SMA) |  | -6 | -6 | 56 |
|  | 7.05 | L Superior Frontal Gyrus (PMd) |  | -20 | 0 | 62 |
|  | 6.92 | N/A (PMd) |  | -16 | -4 | 56 |
|  | 2.95 | R Posterior-Medial Frontal (SMA) |  | 8 | -12 | 62 |
|  | 2.36 | L Middle Frontal Gyrus (PMd) |  | -32 | -2 | 62 |
|  | 2.11 | L Precentral Gyrus (M1) |  | -30 | -14 | 52 |
|  | 2.09 | L Precentral Gyrus (PMd) |  | -34 | -8 | 60 |
|  | 2.06 | L Precentral Gyrus (M1) |  | -36 | -12 | 54 |
| 829 | 8.13 | L Inferior Parietal Lobule (40) |  | -40 | -46 | 54 |
|  | 2.97 | L SupraMarginal Gyrus (2) | L Area PFt (IPL) | -60 | -26 | 46 |
|  | 2.91 | L Superior Parietal Lobule (5) | L Area 5L (SPL) | -16 | -54 | 68 |
|  | 2.74 | L Superior Parietal Lobule | L Area 5L (SPL) | -24 | -54 | 66 |
|  | 2.34 | L Inferior Parietal Lobule (40) |  | -34 | -34 | 42 |
|  | 2.33 | N/A (40) |  | -32 | -36 | 40 |
| 113 | 3.01 | R Precentral Gyrus (M1) |  | 36 | -12 | 60 |
|  | 2.82 | R Middle Frontal Gyrus (pre-SMA) |  | 38 | -8 | 60 |
|  | 2.17 | N/A (PMd) |  | 20 | -10 | 54 |
| 76 | 2.56 | L Cerebelum (VI) | L Lobule VI (Hem) | -22 | -64 | -22 |
| 73 | 2.45 | L Putamen (48) |  | -32 | -6 | 6 |
|  | 1.85 | L Putamen |  | -26 | -10 | 10 |
| 68 | 2.76 | R Postcentral Gyrus (S1) | R Area 1 | 44 | -28 | 58 |
|  | 2.60 | R Postcentral Gyrus (S1) | R Area 3b | 40 | -30 | 52 |
| 32 | 2.21 | N/A (PMv) | L Area 44 | -62 | 4 | 30 |
| 30 | 2.10 | L Temporal Pole (PMv) |  | -56 | 6 | 4 |
| 26 | 2.08 | R Putamen (48) |  | 30 | -4 | -2 |
|  | 1.98 | N/A (48) |  | 26 | -4 | -4 |
| 19 | 1.88 | R Superior Parietal Lobule (7) | R Area 7A (SPL) | 24 | -60 | 64 |
|  | 1.85 | R Superior Parietal Lobule (5) | R Area 5L (SPL) | 20 | -58 | 62 |

#### Table S26: Meta-analysis of Contrast_Auditory_greather_than_Gustatory

| **Cluster Voxels** | **Z-score** | **Macroanatomical Location** | **Cytoarchitectonic/ Tractographic Label** | **MNI Coordinates** | | |
| --- | --- | --- | --- | --- | --- | --- |
|  |  |  |  | **x** | **y** | **z** |
| 342 | 3.92 | L MCC (32) |  | -6 | 20 | 40 |
|  | 3.67 | R MCC (pre-SMA) |  | 2 | 14 | 46 |
|  | 3.51 | L ACC (24) |  | 0 | 22 | 36 |
|  | 3.37 | L Posterior-Medial Frontal (pre-SMA) |  | 0 | 8 | 50 |
|  | 3.35 | L MCC (24) |  | -4 | 12 | 40 |
|  | 3.32 | R MCC (pre-SMA) |  | 4 | 10 | 50 |
|  | 3.29 | R MCC (32) |  | 6 | 18 | 40 |
|  | 3.19 | L MCC (pre-SMA) |  | 0 | 10 | 44 |
|  | 3.16 | R MCC (24) |  | 4 | 22 | 38 |
|  | 3.15 | L Superior Medial Gyrus (pre-SMA) |  | -4 | 14 | 46 |
|  | 2.95 | R MCC (24) |  | 2 | 16 | 38 |
| 155 | 5.03 | L Superior Temporal Gyrus (48) |  | -62 | -42 | 24 |
|  | 3.64 | L Superior Temporal Gyrus (48) | L Area PF (IPL) | -60 | -46 | 28 |
|  | 3.63 | L Superior Temporal Gyrus | L Area PF (IPL) | -66 | -40 | 24 |
|  | 3.57 | L Superior Temporal Gyrus (A1-42) |  | -58 | -42 | 18 |
|  | 3.38 | L Superior Temporal Gyrus (A1-42) | L Area PFcm (IPL) | -52 | -42 | 20 |
|  | 3.16 | L Superior Temporal Gyrus (48) | L Area PFcm (IPL) | -52 | -42 | 26 |
|  | 3.12 | L Superior Temporal Gyrus (22) |  | -64 | -42 | 18 |
|  | 2.99 | L Superior Temporal Gyrus (A1-42) | L Area PFcm (IPL) | -56 | -42 | 28 |
|  | 2.97 | L SupraMarginal Gyrus (48) | L Area PFcm (IPL) | -56 | -44 | 32 |
|  | 2.66 | L SupraMarginal Gyrus (48) | L Area PFcm (IPL) | -52 | -40 | 30 |
|  | 2.47 | L Superior Temporal Gyrus (48) | L Area PF (IPL) | -64 | -44 | 28 |
| 101 | 3.43 | R Superior Temporal Gyrus (22) |  | 54 | -30 | 4 |
|  | 3.23 | R Middle Temporal Gyrus (21) |  | 48 | -40 | 6 |
|  | 2.99 | R Middle Temporal Gyrus (21) |  | 48 | -42 | 10 |
|  | 2.93 | R Middle Temporal Gyrus (21) |  | 46 | -38 | 8 |
|  | 2.71 | R Superior Temporal Gyrus (22) |  | 54 | -24 | 4 |
|  | 2.68 | R Superior Temporal Gyrus (A1-42) |  | 54 | -38 | 10 |
|  | 2.51 | R Middle Temporal Gyrus (21) |  | 50 | -34 | 0 |
| 72 | 3.72 | L Inferior Parietal Lobule (7) | L Area hIP3 (IPS) | -32 | -56 | 50 |
|  | 3.35 | L Inferior Parietal Lobule (7) | L Area hIP3 (IPS) | -32 | -60 | 54 |
|  | 3.29 | N/A (40) | L Area hIP1 (IPS) | -36 | -46 | 40 |
|  | 3.23 | N/A (7) |  | -28 | -54 | 42 |
|  | 2.97 | L Inferior Parietal Lobule (7) | L Area hIP1 (IPS) | -30 | -52 | 46 |
|  | 2.91 | L Inferior Parietal Lobule (40) | L Area hIP1 (IPS) | -32 | -54 | 44 |
|  | 2.81 | L Inferior Parietal Lobule (40) | L Area hIP3 (IPS) | -34 | -56 | 58 |
|  | 2.73 | L Inferior Parietal Lobule (7) | L Area hIP3 (IPS) | -32 | -54 | 54 |
|  | 2.51 | L Inferior Parietal Lobule (40) | L Area hIP1 (IPS) | -38 | -48 | 44 |
|  | 2.39 | N/A (40) |  | -30 | -48 | 42 |
|  | 2.28 | L Inferior Parietal Lobule (7) | L Area hIP3 (IPS) | -36 | -60 | 58 |
| 30 | 4.46 | R Middle Frontal Gyrus (PMd) |  | 50 | 6 | 52 |
|  | 3.34 | R Precentral Gyrus (PMd) |  | 46 | 4 | 56 |
|  | 3.21 | R Precentral Gyrus (PMd) |  | 50 | 2 | 54 |
|  | 2.60 | R Precentral Gyrus (PMd) |  | 52 | 0 | 52 |

#### Table S27: Meta-analysis of Contrast_Gustatory_greather_than_Auditory

| **Cluster Voxels** | **Z-score** | **Macroanatomical Location** | **Cytoarchitectonic/ Tractographic Label** | **MNI Coordinates** | | |
| --- | --- | --- | --- | --- | --- | --- |
|  |  |  |  | **x** | **y** | **z** |
| 46 | 2.91 | L Insula Lobe |  | -40 | 6 | 8 |
|  | 2.79 | L Insula Lobe (48) |  | -40 | 4 | 4 |
|  | 2.70 | L Insula Lobe (pre-SMA) |  | -44 | 2 | 6 |

#### Table S28: Meta-analysis of Contrast_Auditory_greather_than_Tactile

| **Cluster Voxels** | **Z-score** | **Macroanatomical Location** | **Cytoarchitectonic/ Tractographic Label** | **MNI Coordinates** | | |
| --- | --- | --- | --- | --- | --- | --- |
|  |  |  |  | **x** | **y** | **z** |
| 97 | 3.32 | R MCC (24) |  | 4 | 22 | 38 |
|  | 3.12 | R MCC (24) |  | 2 | 16 | 38 |
|  | 2.19 | R MCC (SMA) |  | 6 | 20 | 42 |
|  | 2.19 | R MCC (pre-SMA) |  | 4 | 16 | 44 |
| 39 | 4.07 | R Precentral Gyrus (PMd) |  | 58 | 6 | 44 |
|  | 2.26 | R Precentral Gyrus (PMv) |  | 60 | 2 | 42 |
|  | 2.26 | R Precentral Gyrus (PMd) |  | 58 | 0 | 44 |
|  | 1.88 | R Precentral Gyrus (PMd) |  | 54 | 4 | 42 |
|  | 1.83 | R IFG (p. Opercularis) (PMd) |  | 50 | 8 | 44 |
| 26 | 2.03 | R Posterior-Medial Frontal (pre-SMA) |  | 6 | 14 | 62 |

#### Table S29: Meta-analysis of Contrast_Tactile_greather_than_Auditory

| **Cluster Voxels** | **Z-score** | **Macroanatomical Location** | **Cytoarchitectonic/ Tractographic Label** | **MNI Coordinates** | | |
| --- | --- | --- | --- | --- | --- | --- |
|  |  |  |  | **x** | **y** | **z** |
| 91 | 4.38 | L Posterior-Medial Frontal (pre-SMA) |  | -6 | 10 | 58 |
|  | 2.93 | L Posterior-Medial Frontal (pre-SMA) |  | -8 | 12 | 50 |
|  | 2.62 | L Posterior-Medial Frontal (pre-SMA) |  | -6 | 16 | 52 |
|  | 2.20 | L Posterior-Medial Frontal (pre-SMA) |  | -10 | 16 | 52 |
| 73 | 3.35 | L Inferior Parietal Lobule (40) | L Area hIP1 (IPS) | -34 | -52 | 44 |
|  | 3.16 | N/A (40) | L Area hIP1 (IPS) | -36 | -48 | 40 |

#### Table S30: Meta-analysis of Contrast_Gustatory_greather_than_Olfactory

| **Cluster Voxels** | **Z-score** | **Macroanatomical Location** | **Cytoarchitectonic/ Tractographic Label** | **MNI Coordinates** | | |
| --- | --- | --- | --- | --- | --- | --- |
|  |  |  |  | **x** | **y** | **z** |
| 230 | 5.52 | L Insula Lobe (48) |  | -36 | -4 | 12 |
|  | 4.94 | L Insula Lobe (PMd) |  | -40 | -4 | 14 |
|  | 4.20 | L Insula Lobe (48) |  | -38 | 0 | 10 |
|  | 3.85 | L Insula Lobe (48) |  | -44 | -4 | 8 |
|  | 3.73 | L Insula Lobe (48) |  | -34 | -6 | 16 |
|  | 3.69 | L Insula Lobe (48) |  | -40 | 2 | 4 |
|  | 3.54 | L Insula Lobe (48) |  | -42 | 4 | 6 |

#### Table S31: Meta-analysis of Contrast_Gustatory_greather_than_Visual

| **Cluster Voxels** | **Z-score** | **Macroanatomical Location** | **Cytoarchitectonic/ Tractographic Label** | **MNI Coordinates** | | |
| --- | --- | --- | --- | --- | --- | --- |
|  |  |  |  | **x** | **y** | **z** |
| 230 | 5.57 | L Insula Lobe (48) |  | -38 | -4 | 12 |

#### Table S32: Meta-analysis of Contrast_Gustatory_greather_than_Motor

| **Cluster Voxels** | **Z-score** | **Macroanatomical Location** | **Cytoarchitectonic/ Tractographic Label** | **MNI Coordinates** | | |
| --- | --- | --- | --- | --- | --- | --- |
|  |  |  |  | **x** | **y** | **z** |
| 230 | 5.57 | L Insula Lobe (48) |  | -38 | -4 | 12 |

#### Table S33: Meta-analysis of Contrast_Motor_greather_than_Gustatory

| **Cluster Voxels** | **Z-score** | **Macroanatomical Location** | **Cytoarchitectonic/ Tractographic Label** | **MNI Coordinates** | | |
| --- | --- | --- | --- | --- | --- | --- |
|  |  |  |  | **x** | **y** | **z** |
| 3312 | 8.13 | L MCC (pre-SMA) |  | -4 | 12 | 44 |
|  | 7.61 | L Precentral Gyrus (PMd) |  | -32 | -2 | 50 |
|  | 7.27 | L Middle Frontal Gyrus (PMd) |  | -28 | 2 | 66 |
|  | 7.23 | N/A (PMd) |  | -28 | 2 | 50 |
|  | 7.10 | R Superior Frontal Gyrus (PMd) |  | 26 | -6 | 62 |
|  | 6.63 | N/A (PMd) |  | -22 | 0 | 50 |
|  | 6.21 | L Middle Frontal Gyrus (PMd) |  | -24 | 4 | 54 |
|  | 5.35 | R MCC (pre-SMA) |  | 8 | 14 | 50 |
|  | 5.14 | L MCC (S1) |  | -4 | 0 | 48 |
|  | 4.71 | L Middle Frontal Gyrus (PMd) |  | -36 | 4 | 54 |
|  | 4.56 | R MCC (pre-SMA) |  | 6 | 6 | 50 |
| 2562 | 8.13 | L Inferior Parietal Lobule (40) |  | -36 | -42 | 46 |
|  | 7.51 | L Inferior Parietal Lobule (40) | L Area 2 | -40 | -46 | 58 |
|  | 7.49 | L Inferior Parietal Lobule (40) |  | -38 | -36 | 42 |
|  | 6.71 | L Inferior Parietal Lobule (40) | L Area hIP3 (IPS) | -32 | -52 | 54 |
|  | 5.95 | L Inferior Parietal Lobule (40) |  | -44 | -44 | 56 |
|  | 5.81 | N/A (40) | L Area hIP1 (IPS) | -40 | -46 | 42 |
|  | 5.47 | L Superior Parietal Lobule (7) | L Area 7A (SPL) | -26 | -58 | 64 |
|  | 5.37 | L Superior Parietal Lobule (7) | L Area 7PC (SPL) | -32 | -54 | 62 |
|  | 5.04 | L Inferior Parietal Lobule (40) | L Area hIP2 (IPS) | -46 | -44 | 48 |
|  | 4.89 | L Superior Parietal Lobule (7) |  | -22 | -58 | 52 |
|  | 4.65 | L Inferior Parietal Lobule (7) | L Area hIP1 (IPS) | -30 | -52 | 48 |
| 915 | 5.65 | R Postcentral Gyrus (M1) | R Area 3b | 38 | -36 | 52 |
|  | 3.72 | R Postcentral Gyrus (S1) | R Area 3A | 34 | -34 | 50 |
|  | 3.61 | R SupraMarginal Gyrus (S1) | R Area PFt (IPL) | 52 | -30 | 44 |
|  | 3.54 | R SupraMarginal Gyrus (40) |  | 58 | -34 | 44 |
|  | 3.43 | N/A (40) |  | 34 | -34 | 42 |
|  | 3.35 | R Superior Occipital Gyrus (7) |  | 26 | -66 | 50 |
|  | 3.26 | R SupraMarginal Gyrus (S1) | R Area 2 | 54 | -30 | 48 |
|  | 3.19 | R Postcentral Gyrus (S1) | R Area 2 | 44 | -34 | 54 |
|  | 3.18 | R Inferior Parietal Lobule (2) | R Area 2 | 48 | -38 | 56 |
|  | 3.16 | N/A (7) |  | 28 | -56 | 48 |
|  | 3.13 | R Precuneus (7) |  | 12 | -72 | 54 |
| 319 | 3.54 | N/A (48) |  | 30 | 20 | 8 |
|  | 3.43 | R Putamen |  | 28 | 6 | 4 |
|  | 3.29 | R Putamen (48) |  | 22 | 14 | 4 |
|  | 3.24 | R Putamen |  | 22 | 2 | 4 |
|  | 3.19 | R Insula Lobe (45) |  | 42 | 20 | 4 |
|  | 3.16 | R Putamen |  | 22 | 2 | 10 |
|  | 3.09 | R IFG (p. Orbitalis) (47) |  | 32 | 20 | 0 |
|  | 3.06 | R Putamen (48) |  | 22 | 14 | -2 |
|  | 2.93 | R Putamen (48) |  | 26 | 8 | 0 |
|  | 2.88 | R IFG (p. Orbitalis) (47) |  | 34 | 24 | 0 |
|  | 2.86 | R IFG (p. Opercularis) (PMd) | R Area 45 | 62 | 12 | 12 |
| 224 | 6.12 | L Cerebelum (VI) | L Lobule VI (Hem) | -28 | -60 | -30 |
|  | 3.54 | L Cerebelum (VIII) |  | -28 | -58 | -36 |
|  | 3.49 | N/A |  | -24 | -56 | -28 |
|  | 3.35 | L Cerebelum (VI) | L Lobule VI (Hem) | -28 | -66 | -20 |
|  | 3.19 | N/A |  | -32 | -56 | -36 |
|  | 3.09 | L Cerebelum (Crus 1) | L Lobule VIIa crusI (Hem) | -36 | -58 | -30 |
|  | 3.06 | L Cerebelum (VII) |  | -36 | -52 | -36 |
|  | 3.01 | L Cerebelum (VI) | L Lobule VI (Hem) | -24 | -62 | -26 |
|  | 2.83 | L Cerebelum (VI) | L Lobule VI (Hem) | -36 | -60 | -20 |
|  | 2.82 | L Cerebelum (VI) | L Lobule VI (Hem) | -30 | -56 | -24 |
|  | 2.74 | L Cerebelum (VI) | L Lobule VI (Hem) | -24 | -62 | -20 |
| 127 | 3.72 | R Cerebelum (VI) | R Lobule VI (Hem) | 26 | -68 | -26 |
|  | 3.28 | N/A |  | 30 | -64 | -34 |
|  | 3.19 | R Cerebelum (Crus 1) | R Lobule VIIa crusI (Hem) | 32 | -64 | -30 |
|  | 3.04 | N/A |  | 38 | -54 | -36 |
|  | 3.01 | R Cerebelum (Crus 1) | R Lobule VIIa crusI (Hem) | 42 | -62 | -28 |
|  | 2.95 | R Cerebelum (VI) | R Lobule VI (Hem) | 32 | -64 | -22 |
|  | 2.85 | R Cerebelum (VI) | R Lobule VI (Hem) | 34 | -62 | -24 |
|  | 2.82 | R Cerebelum (VI) | R Lobule VI (Hem) | 24 | -64 | -26 |
|  | 2.77 | R Cerebelum (Crus 1) | R Lobule VIIa crusI (Hem) | 44 | -60 | -22 |
|  | 2.72 | R Cerebelum (Crus 1) | R Lobule VIIa crusI (Hem) | 38 | -62 | -26 |
|  | 2.71 | N/A |  | 36 | -60 | -36 |

#### Table S34: Meta-analysis of Contrast_Motor_greather_than_Olfactory

| **Cluster Voxels** | **Z-score** | **Macroanatomical Location** | **Cytoarchitectonic/ Tractographic Label** | **MNI Coordinates** | | |
| --- | --- | --- | --- | --- | --- | --- |
|  |  |  |  | **x** | **y** | **z** |
| 7856 | 8.13 | L Posterior-Medial Frontal (pre-SMA) |  | -8 | 6 | 50 |
|  | 7.94 | L Middle Frontal Gyrus (PMd) |  | -22 | 2 | 58 |
|  | 7.81 | L Superior Frontal Gyrus (PMd) |  | -22 | -6 | 64 |
|  | 7.61 | L Precentral Gyrus (PMd) |  | -32 | -2 | 50 |
|  | 7.46 | L Precentral Gyrus (PMv) |  | -52 | 2 | 38 |
|  | 7.23 | N/A (PMd) |  | -28 | 2 | 50 |
|  | 7.12 | L Posterior-Medial Frontal (SMA) |  | 0 | -2 | 70 |
|  | 7.11 | L IFG (p. Opercularis) (PMv) |  | -52 | 6 | 26 |
|  | 7.04 | L Precentral Gyrus (PMd) |  | -46 | -6 | 46 |
|  | 6.93 | L Superior Frontal Gyrus (PMd) |  | -20 | 0 | 54 |
|  | 6.82 | L Posterior-Medial Frontal (pre-SMA) |  | -4 | 8 | 64 |
| 555 | 3.72 | R Superior Parietal Lobule (5) |  | 18 | -58 | 62 |
|  | 3.70 | R Superior Occipital Gyrus (pre-SMA) |  | 26 | -66 | 50 |
|  | 3.65 | R Angular Gyrus (7) |  | 30 | -64 | 46 |
|  | 3.54 | R Postcentral Gyrus (S1) | R Area 3a | 34 | -34 | 50 |
|  | 3.44 | N/A (7) |  | 22 | -64 | 46 |
|  | 3.43 | R Superior Parietal Lobule (7) | R Area 7A (SPL) | 20 | -62 | 60 |
|  | 3.35 | N/A (40) |  | 34 | -34 | 42 |
|  | 3.29 | R Precuneus (7) |  | 16 | -66 | 52 |
|  | 3.19 | R Inferior Parietal Lobule (40) | R Area hIP3 (IPS) | 38 | -52 | 50 |
|  | 3.13 | R Precuneus (7) |  | 12 | -72 | 54 |
|  | 3.12 | N/A (40) |  | 50 | -34 | 40 |
| 543 | 8.13 | L Inferior Parietal Lobule (40) |  | -36 | -42 | 46 |
|  | 3.43 | L Inferior Parietal Lobule (40) |  | -36 | -34 | 44 |
|  | 3.35 | L Postcentral Gyrus (S1) | L Area 3a | -38 | -32 | 50 |
|  | 3.24 | L Inferior Parietal Lobule (40) | L Area 2 | -34 | -44 | 56 |
|  | 3.16 | L Postcentral Gyrus (40) | L Area 2 | -36 | -40 | 56 |
|  | 3.06 | N/A (40) |  | -38 | -38 | 38 |
|  | 3.04 | N/A (40) | L Area hIP2 (IPS) | -42 | -46 | 40 |
|  | 2.99 | N/A |  | -32 | -40 | 38 |
|  | 2.97 | N/A (40) | L Area hIP1 (IPS) | -36 | -42 | 40 |
|  | 2.93 | L Inferior Parietal Lobule (S1) | L Area 3a | -38 | -30 | 44 |
|  | 2.83 | N/A (40) | L Area hIP1 (IPS) | -38 | -52 | 40 |
| 294 | 7.13 | R Cerebelum (VI) | R Lobule VI (Hem) | 30 | -58 | -26 |
|  | 3.72 | R Cerebelum (Crus 1) | R Lobule VIIa crusI (Hem) | 32 | -60 | -32 |
|  | 3.54 | R Cerebelum (Crus 1) | R Lobule VIIa crusI (Hem) | 32 | -64 | -30 |
|  | 3.28 | N/A |  | 30 | -64 | -34 |
|  | 3.19 | R Cerebelum (VI) | R Lobule VI (Hem) | 26 | -68 | -26 |
|  | 3.06 | N/A | R Lobule VIIa crusI (Hem) | 36 | -58 | -34 |
|  | 3.01 | N/A |  | 26 | -58 | -32 |
|  | 2.99 | N/A |  | 38 | -54 | -36 |
|  | 2.89 | R Cerebelum (VI) | R Lobule VI (Hem) | 26 | -62 | -24 |
|  | 2.88 | R Cerebelum (Crus 1) | R Lobule VIIa crusI (Hem) | 36 | -58 | -26 |
|  | 2.85 | R Cerebelum (Crus 1) | R Lobule VIIa crusI (Hem) | 42 | -62 | -28 |
| 201 | 4.68 | L Superior Temporal Gyrus (48) | L Area OP1 [SII] | -50 | -34 | 26 |
|  | 4.17 | L Superior Temporal Gyrus (A1-42) |  | -56 | -40 | 22 |
|  | 3.54 | L Superior Temporal Gyrus (48) | L Area PFop (IPL) | -58 | -34 | 26 |
|  | 3.24 | L SupraMarginal Gyrus (48) | L Area PFop (IPL) | -62 | -30 | 28 |
|  | 3.19 | L Superior Temporal Gyrus (48) | L Area PFcm (IPL) | -48 | -38 | 26 |
|  | 3.01 | L SupraMarginal Gyrus (48) | L Area OP1 [SII] | -54 | -30 | 24 |
|  | 2.91 | L Superior Temporal Gyrus (A1-42) | L Area PFcm (IPL) | -54 | -38 | 18 |
|  | 2.81 | L Superior Temporal Gyrus (48) | L Area PFcm (IPL) | -54 | -36 | 22 |
|  | 2.75 | L Postcentral Gyrus (48) | L Area OP1 [SII] | -60 | -24 | 22 |
|  | 2.69 | L Postcentral Gyrus (48) | L Area PFop (IPL) | -64 | -26 | 24 |
|  | 2.51 | L Superior Temporal Gyrus (48) | L Area PF (IPL) | -62 | -36 | 26 |
| 159 | 3.72 | L Cerebelum (VI) | L Lobule VI (Hem) | -28 | -60 | -30 |
|  | 3.54 | L Cerebelum (VIII) |  | -28 | -58 | -36 |
|  | 3.43 | N/A |  | -24 | -56 | -28 |
|  | 3.24 | L Cerebelum (VII) |  | -36 | -52 | -36 |
|  | 3.12 | L Cerebelum (Crus 1) | L Lobule VIIa crusI (Hem) | -38 | -60 | -32 |
|  | 3.06 | L Cerebelum (VI) | L Lobule VI (Hem) | -26 | -58 | -26 |
|  | 2.97 | L Cerebelum (VI) | L Lobule VI (Hem) | -34 | -54 | -24 |
|  | 2.95 | L Cerebelum (Crus 1) | L Lobule VI (Hem) | -34 | -58 | -30 |
|  | 2.91 | L Cerebelum (VI) | L Lobule VI (Hem) | -30 | -56 | -24 |
|  | 2.86 | L Cerebelum (Crus 1) | L Lobule VIIa crusI (Hem) | -38 | -52 | -28 |
|  | 2.59 | L Cerebelum (Crus 1) |  | -36 | -58 | -36 |

#### Table S35: Meta-analysis of Contrast_Tactile_greather_than__Gustatory

| **Cluster Voxels** | **Z-score** | **Macroanatomical Location** | **Cytoarchitectonic/ Tractographic Label** | **MNI Coordinates** | | |
| --- | --- | --- | --- | --- | --- | --- |
|  |  |  |  | **x** | **y** | **z** |
| 113 | 4.51 | N/A (40) |  | -32 | -46 | 44 |
|  | 4.16 | N/A (40) |  | -34 | -44 | 38 |
|  | 3.94 | L Inferior Parietal Lobule (40) | L Area hIP1 (IPS) | -36 | -48 | 44 |
|  | 3.60 | N/A (40) | L Area hIP1 (IPS) | -34 | -52 | 40 |
|  | 3.16 | L Inferior Parietal Lobule (40) |  | -34 | -40 | 42 |
| 27 | 3.35 | L Posterior-Medial Frontal (pre-SMA) | L Area hIP1 (IPS) | -8 | 12 | 50 |
|  | 3.16 | L Posterior-Medial Frontal (pre-SMA) | L Area hIP1 (IPS) | -6 | 8 | 52 |

#### Table S36: Meta-analysis of Contrast_Tactile_greather_than_Olfactory

| **Cluster Voxels** | **Z-score** | **Macroanatomical Location** | **Cytoarchitectonic/ Tractographic Label** | **MNI Coordinates** | | |
| --- | --- | --- | --- | --- | --- | --- |
|  |  |  |  | **x** | **y** | **z** |
| 114 | 4.16 | N/A (40) |  | -34 | -44 | 38 |
|  | 3.38 | L Inferior Parietal Lobule (40) |  | -34 | -40 | 42 |
|  | 3.04 | N/A (40) | L Area hIP1 (IPS) | -34 | -44 | 44 |
|  | 2.30 | N/A (40) | L Area hIP1 (IPS) | -34 | -54 | 40 |
| 71 | 4.35 | L Posterior-Medial Frontal (pre-SMA) |  | -8 | 8 | 54 |
|  | 3.36 | L Superior Frontal Gyrus (pre-SMA) |  | -12 | 8 | 56 |
|  | 3.31 | L Posterior-Medial Frontal (pre-SMA) |  | -8 | 8 | 60 |
|  | 2.29 | L Posterior-Medial Frontal (pre-SMA) |  | -4 | 12 | 58 |
|  | 1.85 | L Posterior-Medial Frontal (pre-SMA) |  | -4 | 12 | 52 |

#### Table S37: Meta-analysis of Contrast_Tactile_greather_than_Visual

| **Cluster Voxels** | **Z-score** | **Macroanatomical Location** | **Cytoarchitectonic/ Tractographic Label** | **MNI Coordinates** | | |
| --- | --- | --- | --- | --- | --- | --- |
|  |  |  |  | **x** | **y** | **z** |
| 112 | 3.72 | N/A (40) |  | -34 | -46 | 38 |
| 93 | 3.19 | L Posterior-Medial Frontal (pre-SMA) |  | -10 | 10 | 52 |

#### Table S38: Meta-analysis of Contrast_Motor_greather_than_Tactile

| **Cluster Voxels** | **Z-score** | **Macroanatomical Location** | **Cytoarchitectonic/ Tractographic Label** | **MNI Coordinates** | | |
| --- | --- | --- | --- | --- | --- | --- |
|  |  |  |  | **x** | **y** | **z** |
| 670 | 5.74 | R Superior Frontal Gyrus (PMd) |  | 26 | -8 | 64 |
|  | 5.55 | N/A (PMd) |  | 24 | -12 | 60 |
|  | 5.23 | R Superior Frontal Gyrus (PMd) |  | 22 | -8 | 62 |
|  | 3.43 | N/A (PMd) |  | 20 | -8 | 56 |
|  | 3.34 | L Posterior-Medial Frontal (SMA) |  | -8 | -14 | 66 |
|  | 3.29 | N/A (PMd) |  | 20 | -12 | 58 |
|  | 3.25 | R Superior Frontal Gyrus (SMA) |  | 16 | -10 | 68 |
|  | 3.24 | L Posterior-Medial Frontal (SMA) |  | -4 | -14 | 64 |
|  | 3.15 | R Posterior-Medial Frontal (SMA) |  | 2 | -14 | 64 |
|  | 3.12 | R Posterior-Medial Frontal (SMA) |  | 8 | -14 | 66 |
|  | 3.01 | R Posterior-Medial Frontal (SMA) |  | 10 | -18 | 68 |
| 477 | 5.55 | R IFG (p. Opercularis) (PMv) | R Area 44 | 52 | 8 | 18 |
|  | 5.09 | R IFG (p. Opercularis) (PMv) | R Area 44 | 54 | 6 | 22 |
|  | 3.54 | R IFG (p. Opercularis) (PMv) |  | 50 | 4 | 34 |
|  | 3.51 | R IFG (p. Opercularis) (PMv) | R Area 45 | 48 | 6 | 26 |
|  | 3.43 | R IFG (p. Opercularis) (PMv) | R Area 45 | 54 | 16 | 18 |
|  | 3.29 | R IFG (p. Opercularis) (PMv) | R Area 45 | 60 | 10 | 34 |
|  | 3.19 | R IFG (p. Triangularis) (PMv) | R Area 45 | 56 | 12 | 28 |
|  | 3.16 | R IFG (p. Opercularis) (PMv) |  | 48 | 12 | 32 |
|  | 3.04 | R IFG (p. Opercularis) (PMv) | R Area 44 | 50 | 12 | 18 |
|  | 2.99 | R IFG (p. Opercularis) (PMv) | R Area 44 | 56 | 10 | 16 |
|  | 2.91 | R IFG (p. Opercularis) (PMv) | R Area 44 | 56 | 8 | 26 |
| 62 | 2.89 | R MCC (24) |  | 2 | 16 | 38 |
|  | 1.86 | R MCC (S1) |  | 6 | 12 | 42 |
| 32 | 2.97 | R Putamen (48) |  | 30 | -4 | -2 |
|  | 2.88 | R Putamen |  | 28 | -6 | 4 |
|  | 2.70 | R Putamen |  | 28 | -2 | 4 |
| 27 | 1.81 | L Superior Temporal Gyrus (48) |  | -52 | -2 | 4 |
|  | 1.71 | N/A (48) |  | -62 | 8 | 4 |

### Section 4: Balanced contrast results (all)

#### Table S39: Meta-analysis of Imagery_Motor_vs_Visual_Balanced_Contrast

| **Cluster Voxels** | **Z-score** | **Macroanatomical Location** | **Cytoarchitectonic/ Tractographic Label** | **MNI Coordinates** | | |
| --- | --- | --- | --- | --- | --- | --- |
|  |  |  |  | **x** | **y** | **z** |
| 840 | 0.37 | L Posterior-Medial Frontal (SMA) |  | -4 | 0 | 60 |
|  | 0.00 | R Posterior-Medial Frontal (pre-SMA) |  | 12 | 4 | 64 |
| 394 | 0.02 | L Middle Frontal Gyrus (PMd) |  | -26 | -6 | 54 |
| 243 | 0.00 | L Inferior Parietal Lobule (40) |  | -36 | -40 | 44 |
|  | 0.00 | N/A (40) | L Area hIP1 (IPS) | -36 | -48 | 40 |

#### Table S40: Meta-analysis of Imagery_Motor_vs_Auditory_Balanced_Contrast

| **Cluster Voxels** | **Z-score** | **Macroanatomical Location** | **Cytoarchitectonic/ Tractographic Label** | **MNI Coordinates** | | |
| --- | --- | --- | --- | --- | --- | --- |
|  |  |  |  | **x** | **y** | **z** |
| 186 | 0.00 | L Posterior-Medial Frontal (pre-SMA) |  | -2 | 2 | 56 |

#### Table S41: Meta-analysis of Imagery_Auditory_vs_Visual_Balanced_Contrast

| **Cluster Voxels** | **Z-score** | **Macroanatomical Location** | **Cytoarchitectonic/ Tractographic Label** | **MNI Coordinates** | | |
| --- | --- | --- | --- | --- | --- | --- |
|  |  |  |  | **x** | **y** | **z** |
| 868 | 0.94 | L Posterior-Medial Frontal (pre-SMA) |  | -4 | 4 | 64 |
|  | 0.92 | L Posterior-Medial Frontal (pre-SMA) |  | 0 | 8 | 60 |
|  | 0.27 | L MCC (SMA) |  | 2 | 16 | 42 |
|  | 0.17 | L Posterior-Medial Frontal (pre-SMA) |  | 2 | 20 | 60 |
| 763 | 0.83 | L IFG (p. Opercularis) (PMv) | L Area 44 | -50 | 12 | 18 |
|  | 0.39 | L IFG (p. Orbitalis) (48) |  | -40 | 18 | 0 |
|  | 0.07 | L Temporal Pole (38) |  | -52 | 4 | -8 |
|  | 0.04 | N/A |  | -58 | 22 | 2 |
| 215 | 0.36 | L Superior Temporal Gyrus (48) |  | -58 | -42 | 24 |
| 212 | 0.77 | R Precentral Gyrus (PMd) |  | 56 | 0 | 46 |
| 175 | 0.49 | L Precentral Gyrus (PMd) |  | -48 | -2 | 46 |
| 153 | 0.20 | N/A (40) |  | -30 | -52 | 40 |
| 120 | 0.04 | R Middle Temporal Gyrus (21) |  | 52 | -32 | -2 |
| 93 | 0.02 | R Putamen (48) |  | 24 | 6 | -2 |

#### Table S42: Meta-analysis of Imagery_Tactile_vs_Visual_Balanced_Contrast

| **Cluster Voxels** | **Z-score** | **Macroanatomical Location** | **Cytoarchitectonic/ Tractographic Label** | **MNI Coordinates** | | |
| --- | --- | --- | --- | --- | --- | --- |
|  |  |  |  | **x** | **y** | **z** |
| 114 | 0.28 | N/A (40) |  | -32 | -44 | 36 |
| 100 | 0.42 | L Posterior-Medial Frontal (pre-SMA) |  | -8 | 12 | 50 |

#### Table S43: Meta-analysis of Imagery_Gustatory_vs_Visual_Balanced_Contrast

| **Cluster Voxels** | **Z-score** | **Macroanatomical Location** | **Cytoarchitectonic/ Tractographic Label** | **MNI Coordinates** | | |
| --- | --- | --- | --- | --- | --- | --- |
|  |  |  |  | **x** | **y** | **z** |
| 230 | 0.76 | N/A (48) |  | -38 | -2 | 8 |
|  | 0.62 | L Insula Lobe (48) |  | -40 | 2 | 4 |
|  | 0.57 | L Insula Lobe (48) |  | -42 | -8 | 6 |
|  | 0.48 | L Insula Lobe (PMd) |  | -42 | 6 | 8 |

#### Table S44: Meta-analysis of Imagery_Tactile_vs_Olfactory_Balanced_Contrast

| **Cluster Voxels** | **Z-score** | **Macroanatomical Location** | **Cytoarchitectonic/ Tractographic Label** | **MNI Coordinates** | | |
| --- | --- | --- | --- | --- | --- | --- |
|  |  |  |  | **x** | **y** | **z** |
| 114 | 0.12 | N/A (40) |  | -32 | -44 | 36 |
|  | 0.04 | N/A (40) | L Area hIP1 (IPS) | -34 | -52 | 38 |
| 100 | 0.12 | L Posterior-Medial Frontal (pre-SMA) |  | -8 | 12 | 50 |

#### Table S45: Meta-analysis of Imagery_Tactile_vs_Gustatory_Balanced_Contrast_

| **Cluster Voxels** | **Z-score** | **Macroanatomical Location** | **Cytoarchitectonic/ Tractographic Label** | **MNI Coordinates** | | |
| --- | --- | --- | --- | --- | --- | --- |
|  |  |  |  | **x** | **y** | **z** |
| 114 | 0.12 | N/A (40) |  | -32 | -44 | 36 |
|  | 0.04 | N/A (40) | L Area hIP1 (IPS) | -34 | -52 | 38 |
| 100 | 0.12 | L Posterior-Medial Frontal (pre-SMA) |  | -8 | 12 | 50 |

#### Table S46: Meta-analysis of Imagery_Gustatory_vs_Olfactory_Balanced_Contrast

| **Cluster Voxels** | **Z-score** | **Macroanatomical Location** | **Cytoarchitectonic/ Tractographic Label** | **MNI Coordinates** | | |
| --- | --- | --- | --- | --- | --- | --- |
|  |  |  |  | **x** | **y** | **z** |
| 230 | 0.76 | N/A (48) |  | -38 | -2 | 8 |
|  | 0.62 | L Insula Lobe (48) |  | -40 | 2 | 4 |
|  | 0.57 | L Insula Lobe (48) |  | -42 | -8 | 6 |
|  | 0.48 | L Insula Lobe (PMd) |  | -42 | 6 | 8 |

#### Table S47: Meta-analysis of Imagery_ Auditory_vs_Tactile_Balanced_Contrast

| **Cluster Voxels** | **Z-score** | **Macroanatomical Location** | **Cytoarchitectonic/ Tractographic Label** | **MNI Coordinates** | | |
| --- | --- | --- | --- | --- | --- | --- |
|  |  |  |  | **x** | **y** | **z** |
| 123 | 0.00 | R Precentral Gyrus (PMd) |  | -56 | -2 | 44 |

### Section 5: Main results (img>rest)

#### Table S48: Meta-analysis of Auditory Imagery (n=26)

| **Cluster Voxels** | **Z-score** | **Macroanatomical Location** | **Cytoarchitectonic/ Tractographic Label** | **MNI Coordinates** | | |
| --- | --- | --- | --- | --- | --- | --- |
|  |  |  |  | **x** | **y** | **z** |
| 541 | 5.82 | L Posterior-Medial Frontal (pre-SMA) |  | -4 | 2 | 66 |
|  | 5.73 | L Posterior-Medial Frontal (pre-SMA) |  | 0 | 10 | 58 |
| 285 | 5.97 | L IFG (p. Opercularis) (PMv) | L Area 44 | -50 | 10 | 20 |
|  | 4.14 | L IFG (p. Opercularis) (PMv) | L Area 44 | -58 | 4 | 18 |
| 280 | 5.18 | L IFG (p. Orbitalis) (47) |  | -40 | 18 | -2 |
|  | 4.53 | L Temporal Pole (38) |  | -52 | 6 | -6 |
| 257 | 7.18 | R Precentral Gyrus (PMd) |  | 54 | 2 | 46 |
|  | 3.53 | R Middle Frontal Gyrus (PMd) |  | 42 | 4 | 58 |
| 219 | 4.73 | L Superior Temporal Gyrus (48) | L Area PFcm (IPL) | -54 | -40 | 24 |
|  | 4.72 | L Superior Temporal Gyrus (A1-42) |  | -62 | -40 | 22 |
|  | 3.55 | L SupraMarginal Gyrus (40) | L Area PFm (IPL) | -62 | -50 | 32 |
| 209 | 7.29 | L Precentral Gyrus (PMd) |  | -50 | -2 | 50 |
| 162 | 4.87 | L Inferior Parietal Lobule (40) | L Area hIP1 (IPS) | -32 | -52 | 44 |
|  | 4.50 | N/A (40) | L Area hIP1 (IPS) | -36 | -48 | 42 |
|  | 4.18 | L Inferior Parietal Lobule (7) | L Area hIP3 (IPS) | -34 | -58 | 56 |
| 160 | 4.30 | L MCC (SMA) |  | 2 | 16 | 42 |
|  | 4.27 | L MCC (24) |  | -4 | 20 | 38 |
|  | 3.40 | L MCC (pre-SMA) |  | -4 | 10 | 44 |
|  | 3.33 | L Superior Medial Gyrus (pre-SMA) |  | -4 | 22 | 46 |
| 109 | 4.09 | R Insula Lobe (48) |  | 48 | 8 | 0 |
|  | 4.00 | R Temporal Pole (48) |  | 58 | 14 | 0 |

#### Table S49: Meta-analysis of Visual Imagery (n=29)

| **Cluster Voxels** | **Z-score** | **Macroanatomical Location** | **Cytoarchitectonic/ Tractographic Label** | **MNI Coordinates** | | |
| --- | --- | --- | --- | --- | --- | --- |
|  |  |  |  | **x** | **y** | **z** |
| 157 | 4.55 | N/A (PMd) |  | 34 | -2 | 50 |
| 119 | 4.50 | L Superior Parietal Lobule (7) |  | -16 | -66 | 54 |

#### Table S50: Meta-analysis of Motor Imagery (n=130)

| **Cluster Voxels** | **Z-score** | **Macroanatomical Location** | **Cytoarchitectonic/ Tractographic Label** | **MNI Coordinates** | | |
| --- | --- | --- | --- | --- | --- | --- |
|  |  |  |  | **x** | **y** | **z** |
| 6440 | 8.13 | L IFG (p. Opercularis) (PMv) | L Area 44 | -52 | 6 | 10 |
|  | 8.08 | N/A (48) | L Area 44 | -54 | 10 | 4 |
|  | 7.53 | L Precentral Gyrus (PMd) |  | -48 | -4 | 46 |
|  | 7.01 | L Putamen |  | -22 | 2 | 6 |
|  | 6.26 | N/A (PMd) |  | -16 | -4 | 56 |
|  | 5.78 | L Insula Lobe (48) |  | -32 | 18 | 8 |
| 2811 | 8.13 | L Inferior Parietal Lobule (40) |  | -36 | -42 | 46 |
|  | 6.54 | L Superior Temporal Gyrus (48) | L Area PFcm (IPL) | -52 | -36 | 24 |
|  | 6.14 | L SupraMarginal Gyrus (40) | L Area PFt (IPL) | -54 | -34 | 40 |
|  | 5.76 | L Inferior Parietal Lobule (7) |  | -28 | -54 | 58 |
|  | 4.09 | L SupraMarginal Gyrus (48) | L Area PFop (IPL) | -62 | -26 | 26 |
| 1964 | 7.99 | R Precentral Gyrus (PMd) |  | 34 | -2 | 54 |
|  | 7.97 | R IFG (p. Opercularis) (PMv) | R Area 44 | 54 | 12 | 12 |
|  | 7.44 | R Precentral Gyrus (PMd) |  | 50 | 2 | 48 |
|  | 7.35 | R IFG (p. Opercularis) (PMv) | R Area 44 | 56 | 10 | 20 |
| 1319 | 7.08 | R SupraMarginal Gyrus (40) | R Area 2 | 40 | -36 | 46 |
|  | 6.11 | R Superior Temporal Gyrus (48) | R Area PFcm (IPL) | 60 | -32 | 24 |
|  | 5.55 | R Superior Parietal Lobule (7) |  | 18 | -62 | 60 |
|  | 4.69 | R Inferior Parietal Lobule (40) | R Area 2 | 38 | -44 | 52 |
|  | 4.45 | R Angular Gyrus (7) |  | 28 | -62 | 50 |
|  | 4.41 | N/A (7) |  | 24 | -56 | 54 |
|  | 4.22 | R Inferior Parietal Lobule (7) | R Area hIP3 (IPS) | 32 | -54 | 46 |
|  | 4.05 | R SupraMarginal Gyrus (40) | R Area PFt (IPL) | 58 | -34 | 42 |
|  | 3.99 | R Postcentral Gyrus (S1) | R Area 2 | 48 | -32 | 56 |
| 518 | 8.04 | R Cerebelum (VI) | R Lobule VI (Hem) | 34 | -56 | -28 |
| 245 | 5.15 | L Cerebelum (VI) | L Lobule VI (Hem) | -30 | -62 | -24 |
|  | 5.01 | L Cerebelum (VI) | L Lobule VI (Hem) | -34 | -56 | -30 |
| 223 | 6.18 | R Putamen |  | 24 | 0 | 2 |
| 153 | 5.72 | R Insula Lobe (48) |  | 34 | 20 | 4 |

### Section 6: Conjunction results (img>rest)

#### Table S51: Meta-analysis of Conjunction between Motor and Auditory Imagery

| **Cluster Voxels** | **Z-score** | **Macroanatomical Location** | **Cytoarchitectonic/ Tractographic Label** | **MNI Coordinates** | | |
| --- | --- | --- | --- | --- | --- | --- |
|  |  |  |  | **x** | **y** | **z** |
| 523 | 5.82 | L Posterior-Medial Frontal (pre-SMA) |  | -4 | 2 | 66 |
|  | 5.73 | L Posterior-Medial Frontal (pre-SMA) |  | 0 | 10 | 58 |
| 234 | 7.09 | R Precentral Gyrus (PMd) |  | 52 | 2 | 48 |
| 206 | 6.88 | L Precentral Gyrus (PMd) |  | -48 | -4 | 50 |
| 175 | 5.24 | L IFG (p. Opercularis) (PMv) | L Area 44 | -50 | 8 | 18 |
|  | 4.14 | L IFG (p. Opercularis) (PMv) | L Area 44 | -58 | 4 | 18 |
| 115 | 4.15 | L MCC (SMA) |  | 0 | 16 | 42 |
|  | 4.03 | L MCC (24) |  | -4 | 18 | 40 |
|  | 3.40 | L MCC (pre-SMA) |  | -4 | 10 | 44 |
|  | 3.33 | L Superior Medial Gyrus (pre-SMA) |  | -4 | 22 | 46 |
| 113 | 4.53 | L Inferior Parietal Lobule (40) | L Area hIP1 (IPS) | -32 | -50 | 44 |
|  | 4.50 | N/A (40) | L Area hIP1 (IPS) | -36 | -48 | 42 |
|  | 3.84 | L Inferior Parietal Lobule (7) | L Area hIP3 (IPS) | -32 | -56 | 52 |
|  | 3.72 | L Inferior Parietal Lobule | L Area hIP3 (IPS) | -34 | -56 | 56 |
| 94 | 4.68 | L Superior Temporal Gyrus (48) | L Area PFcm (IPL) | -54 | -40 | 24 |
| 93 | 4.32 | L Insula Lobe (48) |  | -34 | 20 | 4 |
|  | 4.01 | L Insula Lobe (48) |  | -38 | 14 | 0 |
| 50 | 4.27 | L Temporal Pole (38) |  | -52 | 8 | -4 |

#### Table S52: Meta-analysis of Conjunction between Motor and Visual Imagery

| **Cluster Voxels** | **Z-score** | **Macroanatomical Location** | **Cytoarchitectonic/ Tractographic Label** | **MNI Coordinates** | | |
| --- | --- | --- | --- | --- | --- | --- |
|  |  |  |  | **x** | **y** | **z** |
| 157 | 4.55 | N/A (PMd) |  | 34 | -2 | 50 |
| 114 | 4.50 | L Superior Parietal Lobule (7) |  | -16 | -66 | 54 |

### Section 7: Contrast results (img>rest)

#### Table S53: Meta-analysis of Contrast_Auditory_greater_than_Visual

| **Cluster Voxels** | **Z-score** | **Macroanatomical Location** | **Cytoarchitectonic/ Tractographic Label** | **MNI Coordinates** | | |
| --- | --- | --- | --- | --- | --- | --- |
|  |  |  |  | **x** | **y** | **z** |
| 321 | 4.48 | R Posterior-Medial Frontal (pre-SMA) |  | 6 | 6 | 62 |
|  | 1.72 | R Posterior-Medial Frontal (pre-SMA) |  | 4 | 2 | 52 |
| 224 | 2.78 | L IFG (p. Orbitalis) (48) |  | -42 | 16 | -2 |
|  | 2.70 | N/A (48) | L Area 44 | -50 | 12 | 2 |
|  | 2.68 | L IFG (p. Orbitalis) (47) |  | -42 | 20 | 2 |
|  | 2.64 | L IFG (p. Orbitalis) (48) |  | -38 | 18 | 0 |
|  | 2.58 | L Temporal Pole (38) |  | -52 | 10 | -2 |
|  | 2.48 | L Temporal Pole (38) | L Area TE 3 | -54 | 6 | -2 |
| 219 | 3.72 | L Superior Temporal Gyrus (A1-42) |  | -60 | -44 | 24 |
|  | 3.54 | L Superior Temporal Gyrus (A1-42) |  | -58 | -42 | 20 |
|  | 3.35 | L Superior Temporal Gyrus (A1-42) | L Area PFcm (IPL) | -54 | -40 | 20 |
| 193 | 3.54 | L Precentral Gyrus (M1) |  | -52 | -8 | 48 |
|  | 3.43 | L Precentral Gyrus (PMd) |  | -52 | 0 | 44 |
|  | 3.35 | L Precentral Gyrus (PMd) |  | -50 | -2 | 48 |
| 176 | 3.43 | R Precentral Gyrus (PMd) |  | 56 | 0 | 46 |
| 117 | 2.46 | L IFG (p. Triangularis) (48) |  | -50 | 18 | 24 |
|  | 2.33 | L IFG (p. Triangularis) (48) |  | -50 | 16 | 14 |
| 58 | 2.24 | L MCC (24) |  | -2 | 16 | 38 |
|  | 2.04 | L MCC (SMA) |  | -8 | 20 | 42 |
| 13 | 1.86 | R Temporal Pole (38) |  | 58 | 14 | -4 |

#### Table S54: Meta-analysis of Contrast_Motor_greater_than_Auditory

| **Cluster Voxels** | **Z-score** | **Macroanatomical Location** | **Cytoarchitectonic/ Tractographic Label** | **MNI Coordinates** | | |
| --- | --- | --- | --- | --- | --- | --- |
|  |  |  |  | **x** | **y** | **z** |
| 1507 | 8.13 | L Middle Frontal Gyrus (PMd) |  | -24 | -8 | 54 |
|  | 7.38 | L Posterior-Medial Frontal (SMA) |  | -12 | -2 | 56 |
|  | 7.06 | N/A (SMA) |  | -12 | -6 | 58 |
|  | 6.38 | N/A (SMA) |  | -14 | -4 | 60 |
|  | 6.26 | N/A (PMd) |  | -16 | -4 | 56 |
|  | 3.72 | L MCC (SMA) |  | -4 | -8 | 54 |
|  | 3.54 | N/A (SMA) |  | -12 | -12 | 66 |
|  | 2.58 | L Superior Frontal Gyrus (PMd) |  | -20 | 6 | 68 |
| 1484 | 8.13 | L Inferior Parietal Lobule (40) | L Area 2 | -40 | -42 | 54 |
|  | 6.29 | L Inferior Parietal Lobule (2) | L Area 2 | -44 | -38 | 50 |
|  | 4.71 | L Precuneus (5) |  | -16 | -58 | 64 |
|  | 3.83 | L Precuneus (7) |  | -14 | -62 | 64 |
|  | 3.72 | L Superior Parietal Lobule |  | -16 | -60 | 60 |
|  | 3.54 | L Precuneus (7) |  | -10 | -62 | 54 |
|  | 3.43 | L Precuneus (7) |  | -10 | -62 | 58 |
|  | 2.95 | L Inferior Parietal Lobule | L Area PF (IPL) | -60 | -36 | 48 |
|  | 2.58 | L SupraMarginal Gyrus (2) | L PFop (IPL) | -66 | -28 | 32 |
|  | 2.52 | N/A | L Area PF (IPL) | -66 | -34 | 40 |
|  | 2.49 | L SupraMarginal Gyrus | L Area PFt (IPL) | -66 | -30 | 40 |
| 427 | 3.24 | N/A (PMd) |  | 24 | -8 | 56 |
|  | 3.16 | N/A (PMd) |  | 28 | -12 | 54 |
|  | 3.12 | N/A (PMd) |  | 32 | -12 | 50 |
| 347 | 3.01 | N/A (S1) | R Area 3a | 34 | -30 | 46 |
|  | 2.91 | R Postcentral Gyrus (S1) | R Area 3b | 40 | -32 | 52 |
|  | 2.81 | R Postcentral Gyrus (S1) | R Area 4p | 36 | -34 | 54 |
|  | 2.15 | N/A (40) | R Area 2 | 34 | -42 | 54 |
| 292 | 4.79 | R Superior Parietal Lobule (7) | R Area 7A (SPL) | 22 | -60 | 60 |
| 226 | 3.19 | L Insula Lobe (48) |  | -42 | -2 | 6 |
|  | 2.99 | L Insula Lobe (pre-SMA) |  | -46 | 0 | 6 |
|  | 2.28 | N/A (48) |  | -30 | 10 | 10 |
| 224 | 2.97 | L IFG (p. Opercularis) (PMv) | L Area 44 | -60 | 10 | 30 |
|  | 2.93 | L IFG (p. Opercularis) (PMv) | L Area 44 | -56 | 6 | 32 |
|  | 2.85 | L IFG (p. Opercularis) (PMv) | L Area 44 | -52 | 4 | 32 |
| 89 | 2.39 | R SupraMarginal Gyrus (48) | R Area PFop (IPL) | 60 | -26 | 30 |
|  | 2.23 | R SupraMarginal Gyrus (2) | R Area PFcm (IPL) | 58 | -34 | 34 |
|  | 2.16 | R SupraMarginal Gyrus (40) | R Area PF (IPL) | 58 | -38 | 36 |
| 55 | 2.18 | R IFG (p. Opercularis) (PMv) |  | 50 | 4 | 30 |
| 25 | 2.04 | L Putamen (48) |  | -28 | 0 | 12 |
|  | 1.90 | L Putamen (48) |  | -30 | 0 | 8 |

#### Table S55: Meta-analysis of Contrast_Auditory_greater_than_Motor

| **Cluster Voxels** | **Z-score** | **Macroanatomical Location** | **Cytoarchitectonic/ Tractographic Label** | **MNI Coordinates** | | |
| --- | --- | --- | --- | --- | --- | --- |
|  |  |  |  | **x** | **y** | **z** |
| 541 | 5.82 | L Posterior-Medial Frontal (pre-SMA) |  | -4 | 2 | 66 |
|  | 5.73 | L Posterior-Medial Frontal (pre-SMA) |  | 0 | 10 | 58 |
| 284 | 5.97 | L IFG (p. Opercularis) (PMv) | L Area 44 | -50 | 10 | 20 |
|  | 4.14 | L IFG (p. Opercularis) (PMv) | L Area 44 | -58 | 4 | 18 |
| 259 | 4.66 | L IFG (p. Triangularis) (48) |  | -36 | 20 | 4 |
|  | 4.33 | L IFG (p. Orbitalis) (48) |  | -40 | 16 | 2 |
|  | 4.29 | L IFG (p. Orbitalis) (48) |  | -40 | 16 | -4 |
|  | 4.27 | L Temporal Pole (38) |  | -52 | 8 | -4 |
|  | 3.72 | L IFG (p. Orbitalis) (47) |  | -38 | 22 | -4 |
| 256 | 7.18 | R Precentral Gyrus (PMd) |  | 54 | 2 | 46 |
|  | 3.29 | R Middle Frontal Gyrus (PMd) |  | 40 | 4 | 58 |
| 209 | 7.29 | L Precentral Gyrus (PMd) |  | -50 | -2 | 50 |
| 182 | 4.73 | L Superior Temporal Gyrus (48) | L Area PFcm (IPL) | -54 | -40 | 24 |
|  | 4.65 | L Superior Temporal Gyrus (48) |  | -60 | -40 | 24 |
| 162 | 4.87 | L Inferior Parietal Lobule (40) | L Area hIP1 (IPS) | -32 | -52 | 44 |
|  | 4.50 | N/A (40) | L Area hIP1 (IPS) | -36 | -48 | 42 |
|  | 3.88 | L Inferior Parietal Lobule (7) | L Area hIP3 (IPS) | -34 | -56 | 52 |
|  | 3.72 | L Inferior Parietal Lobule (7) | L Area 7A (SPL) | -34 | -58 | 58 |
| 160 | 4.30 | L MCC (SMA) |  | 2 | 16 | 42 |
|  | 4.03 | L MCC (24) |  | -4 | 18 | 40 |
|  | 3.40 | L MCC (pre-SMA) |  | -4 | 10 | 44 |
|  | 3.33 | L Superior Medial Gyrus (pre-SMA) |  | -4 | 22 | 46 |
| 66 | 3.63 | R Insula Lobe (48) |  | 48 | 8 | 2 |
|  | 3.50 | N/A (48) |  | 56 | 12 | 2 |

#### Table S56: Meta-analysis of Contrast_Motor_greater_than_Visual

| **Cluster Voxels** | **Z-score** | **Macroanatomical Location** | **Cytoarchitectonic/ Tractographic Label** | **MNI Coordinates** | | |
| --- | --- | --- | --- | --- | --- | --- |
|  |  |  |  | **x** | **y** | **z** |
| 2085 | 8.13 | L Posterior-Medial Frontal (SMA) |  | -6 | -2 | 54 |
|  | 4.74 | R Posterior-Medial Frontal (SMA) |  | 12 | -2 | 64 |
|  | 4.59 | R Posterior-Medial Frontal (pre-SMA) |  | 10 | 2 | 66 |
|  | 4.27 | R Posterior-Medial Frontal (pre-SMA) |  | 12 | 4 | 64 |
|  | 3.72 | L Posterior-Medial Frontal (SMA) |  | 0 | -6 | 60 |
|  | 3.54 | R Posterior-Medial Frontal (SMA) |  | 6 | -2 | 64 |
|  | 3.35 | L Precentral Gyrus (PMv) |  | -56 | 0 | 42 |
|  | 3.29 | L Posterior-Medial Frontal (pre-SMA) |  | 0 | 8 | 68 |
|  | 3.19 | L Precentral Gyrus (PMv) |  | -56 | 0 | 38 |
|  | 2.88 | L Precentral Gyrus (PMd) |  | -50 | -6 | 48 |
|  | 2.74 | L Superior Frontal Gyrus (PMd) |  | -16 | 2 | 68 |
| 992 | 3.54 | L Inferior Parietal Lobule (2) | L Area 2 | -36 | -44 | 58 |
|  | 2.89 | L SupraMarginal Gyrus (48) | L Area PFop (IPL) | -58 | -32 | 26 |
|  | 2.54 | L SupraMarginal Gyrus (2) | L Area PFop (IPL) | -64 | -28 | 30 |
|  | 2.37 | L Inferior Parietal Lobule (2) | L Area PFt (IPL) | -46 | -32 | 42 |
|  | 1.98 | L SupraMarginal Gyrus (S1) | L Area 2 | -52 | -28 | 48 |
| 419 | 3.72 | L Temporal Pole (PMv) | L Area 44 | -54 | 8 | 4 |
|  | 3.43 | N/A (48) | L Area 44 | -50 | 10 | 4 |
|  | 1.70 | L Insula Lobe (48) |  | -40 | 14 | 4 |
| 316 | 3.33 | R Postcentral Gyrus (S1) | R Area 4p | 40 | -26 | 44 |
|  | 3.12 | R Postcentral Gyrus (S1) | R Area 3b | 42 | -28 | 48 |
|  | 2.25 | R Rolandic Operculum (48) | R Area PFcm (IPL) | 54 | -32 | 26 |
|  | 2.24 | R Rolandic Operculum (48) | R Area PFcm (IPL) | 56 | -34 | 28 |
|  | 2.19 | R SupraMarginal Gyrus (48) | R Area PFcm (IPL) | 58 | -36 | 30 |
|  | 2.10 | R SupraMarginal Gyrus (40) | R Area PFcm (IPL) | 50 | -32 | 40 |
|  | 1.98 | R SupraMarginal Gyrus (40) | R Area PFcm (IPL) | 58 | -36 | 36 |
|  | 1.92 | R SupraMarginal Gyrus (40) | R Area PF (IPL) | 62 | -38 | 36 |
| 312 | 3.06 | R IFG (p. Opercularis) (PMv) | R Area 44 | 52 | 8 | 12 |
| 168 | 2.64 | L Putamen (48) |  | -24 | 10 | 8 |
|  | 2.07 | L Putamen |  | -24 | -4 | 0 |
| 137 | 2.11 | N/A (19) |  | 22 | -58 | -26 |
|  | 2.08 | R Cerebelum (Crus 1) | R Lobule VI (Hem) | 34 | -58 | -28 |
|  | 2.04 | R Cerebelum (VI) | R Lobule VI (Hem) | 32 | -60 | -26 |
|  | 2.00 | R Cerebelum (Crus 1) | R Lobule VI (Hem) | 36 | -50 | -30 |
|  | 2.00 | N/A (37) | R Lobule VI (Hem) | 26 | -58 | -28 |
| 128 | 2.58 | R Putamen |  | 28 | -4 | 6 |
|  | 2.54 | R Putamen |  | 24 | 0 | 10 |
| 108 | 2.42 | R Precentral Gyrus (PMd) |  | 48 | -4 | 44 |
| 49 | 2.15 | N/A (7) | R Area 7PC (SPL) | 28 | -52 | 54 |
|  | 2.07 | N/A (7) |  | 30 | -54 | 50 |
|  | 2.02 | R Inferior Parietal Lobule (40) | R Area hIP3 (IPS) | 34 | -48 | 52 |
|  | 1.84 | R Superior Parietal Lobule (5) | R Area 7PC (SPL) | 22 | -56 | 62 |
|  | 1.78 | R Superior Parietal Lobule (7) | Area 7A (SPL) | 20 | -62 | 68 |

#### Table S57: Meta-analysis of Contrast_Visual_greater_than_Motor

| **Cluster Voxels** | **Z-score** | **Macroanatomical Location** | **Cytoarchitectonic/ Tractographic Label** | **MNI Coordinates** | | |
| --- | --- | --- | --- | --- | --- | --- |
|  |  |  |  | **x** | **y** | **z** |
| 157 | 4.55 | N/A (PMd) |  | 34 | -2 | 50 |
| 114 | 4.50 | L Superior Parietal Lobule (7) |  | -16 | -66 | 54 |

#### Table S58: Meta-analysis of Contrast_Auditory_greater_than_Tactile

| **Cluster Voxels** | **Z-score** | **Macroanatomical Location** | **Cytoarchitectonic/ Tractographic Label** | **MNI Coordinates** | | |
| --- | --- | --- | --- | --- | --- | --- |
|  |  |  |  | **x** | **y** | **z** |
| 99 | 4.05 | R Precentral Gyrus (PMd) |  | 48 | 2 | 54 |
|  | 3.43 | R Precentral Gyrus (PMd) |  | 52 | -4 | 46 |
|  | 3.36 | R Middle Frontal Gyrus (PMd) |  | 50 | 10 | 46 |
|  | 3.12 | R Precentral Gyrus (PMd) |  | 48 | 2 | 48 |
|  | 3.04 | R Precentral Gyrus (PMd) |  | 52 | 6 | 44 |
|  | 2.95 | R Precentral Gyrus (PMd) |  | 56 | 4 | 42 |
|  | 2.93 | R Precentral Gyrus (PMd) |  | 50 | -2 | 50 |
|  | 2.89 | R Precentral Gyrus (PMd) |  | 52 | 0 | 44 |
|  | 2.78 | R Precentral Gyrus (PMd) |  | 54 | 2 | 50 |
|  | 2.52 | R Precentral Gyrus (PMv) |  | 52 | 2 | 40 |
|  | 2.30 | R Middle Frontal Gyrus (PMd) |  | 48 | 6 | 56 |
| 14 | 2.95 | L SupraMarginal Gyrus (48) | L Area PF (IPL) | -62 | -44 | 30 |
|  | 2.72 | L SupraMarginal Gyrus (40) | L Area PFm (IPL) | -60 | -50 | 32 |
|  | 2.29 | L SupraMarginal Gyrus (48) | L Area PFcm (IPL) | -54 | -44 | 32 |
|  | 2.22 | L SupraMarginal Gyrus | L Area PFm (IPL) | -64 | -52 | 32 |

#### Table S59: Meta-analysis of Contrast_Motor_greather_than_Tactile

| **Cluster Voxels** | **Z-score** | **Macroanatomical Location** | **Cytoarchitectonic/ Tractographic Label** | **MNI Coordinates** | | |
| --- | --- | --- | --- | --- | --- | --- |
|  |  |  |  | **x** | **y** | **z** |
| 576 | 5.64 | L Precentral Gyrus (PMd) |  | -32 | 0 | 52 |
|  | 3.96 | N/A (pre-SMA) |  | -22 | -16 | 56 |
|  | 3.85 | L Precentral Gyrus (M1) |  | -32 | -16 | 58 |
|  | 3.72 | L Middle Frontal Gyrus (PMd) |  | -30 | -4 | 60 |
|  | 3.43 | L Precentral Gyrus (PMd) |  | -34 | -8 | 52 |
|  | 3.35 | L Middle Frontal Gyrus (PMd) |  | -32 | 0 | 62 |
|  | 3.33 | N/A (PMd) |  | -16 | -8 | 64 |
|  | 3.24 | L Middle Frontal Gyrus (PMd) |  | -26 | -8 | 60 |
|  | 3.19 | N/A (PMd) |  | -20 | -8 | 50 |
|  | 3.16 | N/A (PMd) |  | -26 | -4 | 50 |
|  | 3.12 | L Precentral Gyrus (PMd) |  | -34 | -10 | 56 |
| 513 | 5.08 | R Superior Frontal Gyrus (PMd) |  | 26 | -6 | 64 |
|  | 3.72 | R IFG (p. Opercularis) (PMv) | R Area 44 | 50 | 10 | 16 |
|  | 3.68 | R Middle Frontal Gyrus (PMd) |  | 38 | 4 | 56 |
|  | 3.54 | R Precentral Gyrus (PMd) |  | 52 | -2 | 46 |
|  | 3.43 | R Precentral Gyrus (PMd) |  | 42 | -4 | 46 |
|  | 3.29 | R Precentral Gyrus (PMd) |  | 52 | -6 | 48 |
|  | 3.27 | R Precentral Gyrus (PMv) |  | 54 | 2 | 32 |
|  | 3.24 | R Superior Frontal Gyrus (PMd) |  | 22 | -4 | 62 |
|  | 3.16 | R IFG (p. Opercularis) (PMv) |  | 54 | 8 | 18 |
|  | 3.12 | N/A (PMd) | R Area 44 | 26 | -4 | 54 |
|  | 3.06 | R IFG (p. Opercularis) (PMv) | R Area 45 | 52 | 14 | 16 |
| 35 | 3.35 | R Posterior-Medial Frontal (pre-SMA) |  | 10 | 12 | 54 |
|  | 3.15 | R MCC (pre-SMA) |  | 12 | 14 | 50 |
|  | 2.81 | R Posterior-Medial Frontal (pre-SMA) |  | 12 | 18 | 56 |
|  | 2.77 | R MCC (pre-SMA) |  | 10 | 6 | 50 |
|  | 2.44 | R Posterior-Medial Frontal (pre-SMA) |  | 10 | 2 | 54 |
|  | 2.04 | R Posterior-Medial Frontal (SMA) |  | 10 | -2 | 54 |
| 24 | 3.16 | R Superior Parietal Lobule (7) | R Area 7A (SPL) | 22 | -64 | 62 |
|  | 2.67 | R Precuneus (5) |  | 14 | -60 | 64 |
|  | 2.62 | R Superior Parietal Lobule (7) | R Area 7A (SPL) | 16 | -64 | 66 |
|  | 2.14 | R Superior Parietal Lobule (5) | R Area 5L (SPL) | 20 | -58 | 66 |
|  | 1.94 | R Precuneus (5) |  | 14 | -58 | 58 |
|  | 1.93 | R Precuneus (5) |  | 14 | -62 | 58 |
|  | 1.92 | R Superior Parietal Lobule (5) |  | 18 | -60 | 62 |
| 15 | 2.73 | L IFG (p. Opercularis) (PMv) | L Area 44 | -50 | 6 | 32 |
|  | 2.58 | L IFG (p. Opercularis) (PMv) |  | -44 | 6 | 32 |
|  | 2.51 | L IFG (p. Opercularis) (PMv) | L Area 44 | -46 | 2 | 32 |
|  | 2.31 | L Precentral Gyrus (PMv) | L Area 44 | -54 | 0 | 32 |
|  | 1.80 | L Precentral Gyrus (PMv) |  | -50 | -2 | 32 |
| 13 | 2.78 | N/A (48) |  | 32 | 16 | 6 |
|  | 2.68 | R Insula Lobe (48) |  | 38 | 18 | 8 |
|  | 2.36 | R Insula Lobe (48) |  | 38 | 22 | 8 |

### Section 8: Main results (img>rest_control)

#### Table S60: Meta-analysis of Auditory Imagery (n=35)

| **Cluster Voxels** | **Z-score** | **Macroanatomical Location** | **Cytoarchitectonic/ Tractographic Label** | **MNI Coordinates** | | |
| --- | --- | --- | --- | --- | --- | --- |
|  |  |  |  | **x** | **y** | **z** |
| 887 | 5.95 | L Posterior-Medial Frontal (pre-SMA) |  | 0 | 10 | 58 |
|  | 5.55 | L Posterior-Medial Frontal (pre-SMA) |  | -6 | 6 | 62 |
|  | 4.39 | L MCC (SMA) |  | 2 | 16 | 42 |
|  | 4.31 | L MCC (24) |  | -4 | 20 | 38 |
|  | 4.16 | L MCC (pre-SMA) |  | -6 | 10 | 44 |
|  | 3.69 | L Posterior-Medial Frontal (pre-SMA) |  | 0 | 20 | 62 |
|  | 3.17 | L Superior Medial Gyrus (pre-SMA) |  | -4 | 22 | 46 |
| 377 | 5.67 | L IFG (p. Orbitalis) (48) |  | -40 | 18 | 0 |
|  | 4.31 | L Temporal Pole (38) |  | -52 | 6 | -6 |
|  | 3.83 | N/A (48) |  | -50 | 12 | 0 |
|  | 3.29 | L IFG (p. Triangularis) (S1) | L Area44 | -50 | 12 | 6 |
| 297 | 5.77 | L IFG (p. Opercularis) (PMv) | L Area44 | -50 | 10 | 20 |
|  | 3.92 | L IFG (p. Opercularis) (PMv) | L Area44 | -58 | 4 | 18 |
| 242 | 6.91 | R Precentral Gyrus (PMd) |  | 54 | 2 | 46 |
|  | 3.33 | R Middle Frontal Gyrus (PMd) |  | 42 | 4 | 58 |
| 195 | 7.01 | L Precentral Gyrus (PMd) |  | -50 | -2 | 50 |
| 142 | 4.69 | L Inferior Parietal Lobule (40) | L Area hIP1 (IPS) | -32 | -52 | 44 |
|  | 4.29 | N/A (40) | L Area hIP1 (IPS) | -36 | -48 | 42 |
|  | 4.03 | L Inferior Parietal Lobule (7) | L Area hIP3 (IPS) | -34 | -58 | 56 |
|  | 3.88 | L Inferior Parietal Lobule (7) | L Area hIP1 (IPS) | -32 | -54 | 50 |

#### Table S61: Meta-analysis of Visual Imagery (n=56)

| **Cluster Voxels** | **Z-score** | **Macroanatomical Location** | **Cytoarchitectonic/ Tractographic Label** | **MNI Coordinates** | | |
| --- | --- | --- | --- | --- | --- | --- |
|  |  |  |  | **x** | **y** | **z** |
| 670 | 6.00 | L Superior Parietal Lobule (7) |  | -20 | -68 | 52 |
|  | 4.52 | L Inferior Parietal Lobule (7) | L Area hIP3 (IPS) | -30 | -60 | 48 |
|  | 4.09 | L Inferior Parietal Lobule (40) | L Area hIP2 (IPS) | -52 | -44 | 46 |
|  | 3.96 | L Inferior Parietal Lobule (40) | L Area PFt (IPL) | -42 | -40 | 46 |
| 265 | 4.53 | R Superior Occipital Gyrus (7) |  | 24 | -72 | 40 |
|  | 4.34 | N/A (7) |  | 20 | -64 | 52 |
| 209 | 5.14 | L Inferior Temporal Gyrus (37) |  | -52 | -56 | -10 |
|  | 3.45 | L Inferior Occipital Gyrus (19) | L Area FG2 | -46 | -70 | -12 |
| 167 | 4.87 | R Superior Frontal Gyrus (PMd) |  | 26 | -4 | 60 |
|  | 4.56 | N/A (PMd) |  | 34 | -2 | 50 |
| 140 | 4.44 | R Insula Lobe (48) |  | 46 | 12 | -4 |
| 127 | 4.59 | L IFG (p. Opercularis) (PMv) | L Area 44 | -48 | 8 | 26 |

#### Table S62: Meta-analysis of Motor Imagery (n=208)

| **Cluster Voxels** | **Z-score** | **Macroanatomical Location** | **Cytoarchitectonic/ Tractographic Label** | **MNI Coordinates** | | |
| --- | --- | --- | --- | --- | --- | --- |
|  |  |  |  | **x** | **y** | **z** |
| 10020 | 8.13 | L Temporal Pole (48) |  | -52 | 8 | 0 |
|  | 7.83 | R IFG (p. Opercularis) (PMv) | L Area 44 | 56 | 12 | 12 |
|  | 7.81 | R IFG (p. Opercularis) (PMv) | L Area 44 | 56 | 10 | 22 |
|  | 7.48 | R Precentral Gyrus (PMd) |  | 52 | 2 | 46 |
|  | 7.29 | L Precentral Gyrus (PMd) |  | -46 | -4 | 46 |
|  | 6.47 | R IFG (p. Opercularis) (PMv) |  | 54 | 8 | 34 |
|  | 6.32 | L Insula Lobe (48) |  | -32 | 16 | 6 |
|  | 5.88 | R Insula Lobe (48) |  | 36 | 20 | 4 |
|  | 4.25 | R IFG (p. Opercularis) (47) |  | -38 | 18 | -6 |
|  | 4.22 | L Superior Medial Gyrus (32) |  | -8 | 28 | 38 |
| 3225 | 8.13 | L Inferior Parietal Lobule (40) |  | -36 | -42 | 44 |
|  | 7.19 | L Superior Parietal Lobule (7) | L Area 7A (SPL) | -18 | -64 | 56 |
|  | 6.25 | L Inferior Parietal Lobule (7) | L Area hIP3 (IPS) | -30 | -54 | 58 |
|  | 6.14 | L Superior Temporal Gyrus (48) | L Area PFcm (IPL) | -52 | -36 | 24 |
|  | 4.79 | L SupraMarginal Gyrus (48) | L Area PFop (IPL) | -62 | -26 | 32 |
| 1646 | 7.17 | R SupraMarginal Gyrus (40) | R Area 2 | 40 | -36 | 46 |
|  | 5.95 | R Superior Parietal Lobule (7) |  | 18 | -62 | 60 |
|  | 5.68 | R Superior Temporal Gyrus (48) | R Area PFcm (IPL) | 60 | -32 | 22 |
|  | 4.78 | R Inferior Parietal Lobule (40) | R Area hIP3 (IPS) | 36 | -46 | 50 |
|  | 4.65 | R Angular Gyrus (7) | R Area hIP3 (IPS) | 28 | -62 | 52 |
|  | 4.47 | R Angular Gyrus (7) | R Area hIP1 (IPS) | 32 | -54 | 44 |
|  | 4.46 | R SupraMarginal Gyrus (40) |  | 58 | -34 | 44 |
|  | 4.44 | R Postcentral Gyrus (S1) | R Area 2 | 48 | -32 | 56 |
| 615 | 8.13 | R Cerebelum (Crus 1) | R Lobule VIIa crusI (Hem) | 34 | -58 | -30 |
| 417 | 6.60 | N/A | L Lobule VI (Hem) | -34 | -56 | -32 |
|  | 6.50 | L Cerebelum (VI) | L Lobule VI (Hem) | -30 | -62 | -24 |
|  | 6.44 | L Cerebelum (Crus 1) | L Lobule VI (Hem) | -32 | -60 | -28 |
| 336 | 6.78 | R Putamen |  | 26 | 0 | 2 |

#### Table S63: Meta-analysis of Tactile Imagery (n=6)

| **Cluster Voxels** | **Z-score** | **Macroanatomical Location** | **Cytoarchitectonic/ Tractographic Label** | **MNI Coordinates** | | |
| --- | --- | --- | --- | --- | --- | --- |
|  |  |  |  | **x** | **y** | **z** |
| 119 | 4.90 | N/A (40) | L Area hIP1 (IPS) | 34 | -46 | 42 |
| 104 | 5.32 | L Posterior-Medial Frontal (PMd) |  | -8 | 12 | 56 |

### Section 9: Conjunction results (img> rest_control)

#### Table S64: Meta-analysis of Conjunction between Motor and Auditory Imagery

| **Cluster Voxels** | **Z-score** | **Macroanatomical Location** | **Cytoarchitectonic/ Tractographic Label** | **MNI Coordinates** | | |
| --- | --- | --- | --- | --- | --- | --- |
|  |  |  |  | **x** | **y** | **z** |
| 811 | 5.95 | L Posterior-Medial Frontal (pre-SMA) |  | 0 | 10 | 58 |
|  | 5.55 | L Posterior-Medial Frontal (pre-SMA) |  | -6 | 6 | 62 |
|  | 4.39 | L MCC (SMA) |  | 2 | 16 | 42 |
|  | 4.16 | L MCC (pre-SMA) |  | -6 | 10 | 44 |
|  | 4.13 | L MCC (24) |  | -4 | 20 | 40 |
|  | 3.17 | L Posterior-Medial Frontal (pre-SMA) |  | -4 | 22 | 46 |
| 266 | 5.25 | L Insula Lobe (48) |  | -34 | 20 | 4 |
|  | 4.13 | L Temporal Pole (38) |  | -52 | 6 | -4 |
|  | 4.09 | L IFG (p. Orbitalis) (47) |  | -40 | 18 | -6 |
|  | 3.83 | N/A (48) |  | -50 | 12 | 0 |
|  | 3.29 | L IFG (p. Triangularis) (S1) | L Area 44 | -50 | 12 | 6 |
| 231 | 6.91 | R Precentral Gyrus (PMd) |  | 54 | 2 | 46 |
|  | 3.33 | R Middle Frontal Gyrus (PMd) |  | 42 | 4 | 58 |
| 196 | 5.39 | L IFG (p. Opercularis) (PMv) | L Area 44 | -50 | 8 | 18 |
|  | 3.92 | L IFG (p. Opercularis) (PMv) | L Area 44 | -58 | 4 | 18 |
| 193 | 6.65 | L Precentral Gyrus (PMd) |  | -48 | -4 | 50 |
| 127 | 4.53 | L Inferior Parietal Lobule (40) | L Area hIP1 (IPS) | -32 | -50 | 44 |
|  | 4.50 | N/A (40) | L Area hIP1 (IPS) | -36 | -48 | 42 |
|  | 3.84 | L Inferior Parietal Lobule (7) | L Area hIP3 (IPS) | -32 | -56 | 52 |
|  | 3.72 | L Inferior Parietal Lobule (7) | L Area hIP1 (IPS) | -34 | -54 | 50 |
| 79 | 4.33 | L Superior Temporal Gyrus (48) | L Area PFcm (IPL) | -58 | -38 | 24 |

#### Table S65: Meta-analysis of Conjunction between Motor and Visual Imagery

| **Cluster Voxels** | **Z-score** | **Macroanatomical Location** | **Cytoarchitectonic/ Tractographic Label** | **MNI Coordinates** | | |
| --- | --- | --- | --- | --- | --- | --- |
|  |  |  |  | **x** | **y** | **z** |
| 425 | 5.86 | L Superior Parietal Lobule (7) |  | -20 | -66 | 52 |
|  | 3.96 | L Inferior Parietal Lobule (40) | L Area PFt (IPL) | -42 | -40 | 46 |
|  | 3.84 | L Inferior Parietal Lobule (7) | L Area hIP3 (IPS) | -30 | -60 | 54 |
|  | 3.71 | L Inferior Parietal Lobule (7) | L Area hIP1 (IPS) | -34 | -56 | 44 |
|  | 3.69 | L Inferior Parietal Lobule (40) | L Area hIP2 (IPS) | -50 | -42 | 46 |
| 167 | 4.87 | R Superior Frontal Gyrus (PMd) |  | 26 | -4 | 60 |
|  | 4.56 | N/A (PMd) |  | 34 | -2 | 50 |
| 96 | 4.33 | N/A (7) |  | 20 | -64 | 54 |
| 96 | 4.59 | L IFG (p. Opercularis) (PMv) | L Area 44 | -48 | 8 | 26 |
| 21 | 3.70 | R Insula Lobe (48) |  | 46 | 16 | 4 |
|  | 3.60 | R Insula Lobe (48) |  | 46 | 12 | 2 |

#### Table S66: Meta-analysis of Conjunction between Auditory and Visual Imagery

| **Cluster Voxels** | **Z-score** | **Macroanatomical Location** | **Cytoarchitectonic/ Tractographic Label** | **MNI Coordinates** | | |
| --- | --- | --- | --- | --- | --- | --- |
|  |  |  |  | **x** | **y** | **z** |
| 59 | 4.33 | L IFG (p. Opercularis) (PMv) |  | -48 | 10 | 24 |
| 40 | 3.58 | L Inferior Parietal Lobule (7) | L Area hIP3 (IPS) | -32 | -58 | 56 |
|  | 3.46 | L Inferior Parietal Lobule (7) | L Area hIP3 (IPS) | -32 | -56 | 48 |
|  | 3.40 | L Inferior Parietal Lobule (7) | L Area hIP1 (IPS) | -34 | -54 | 44 |
|  | 3.35 | L Inferior Parietal Lobule (40) | L Area hIP1 (IPS) | -38 | -46 | 44 |

#### Table S67: Meta-analysis of Conjunction between Auditory and Motor and Visual Imagery

| **Cluster Voxels** | **Z-score** | **Macroanatomical Location** | **Cytoarchitectonic/ Tractographic Label** | **MNI Coordinates** | | |
| --- | --- | --- | --- | --- | --- | --- |
|  |  |  |  | **x** | **y** | **z** |
| 43 | 4.31 | L IFG (p. Opercularis) (PMv) | L Area 44 | -48 | 8 | 24 |
| 40 | 3.58 | L Inferior Parietal Lobule (7) | L Area hIP3 (IPS) | -32 | -58 | 56 |
|  | 3.46 | L Inferior Parietal Lobule (7) | L Area hIP3 (IPS) | -32 | -56 | 48 |
|  | 3.40 | L Inferior Parietal Lobule (7) | L Area hIP1 (IPS) | -34 | -54 | 44 |
|  | 3.35 | L Inferior Parietal Lobule (40) | L Area hIP1 (IPS) | -38 | -46 | 44 |

#### Table S68: Meta-analysis of Conjunction between Motor and Tactile Imagery

| **Cluster Voxels** | **Z-score** | **Macroanatomical Location** | **Cytoarchitectonic/ Tractographic Label** | **MNI Coordinates** | | |
| --- | --- | --- | --- | --- | --- | --- |
|  |  |  |  | **x** | **y** | **z** |
| 99 | 5.32 | L Posterior-Medial Frontal (pre-SMA) |  | -8 | 12 | 56 |
| 98 | 4.90 | N/A (40) | L Area hIP1 (IPS) | -34 | -46 | 42 |

#### Table S69: Meta-analysis of Conjunction between Motor and Tactile and Auditory Imagery

| **Cluster Voxels** | **Z-score** | **Macroanatomical Location** | **Cytoarchitectonic/ Tractographic Label** | **MNI Coordinates** | | |
| --- | --- | --- | --- | --- | --- | --- |
|  |  |  |  | **x** | **y** | **z** |
| 53 | 4.43 | L Posterior-Medial Frontal (pre-SMA) |  | -6 | 10 | 58 |
| 46 | 4.25 | N/A (40) | L Area hIP1 (IPS) | -34 | -48 | 42 |

#### Table S70: Meta-analysis of Conjunction between Auditory and Tactile Imagery

| **Cluster Voxels** | **Z-score** | **Macroanatomical Location** | **Cytoarchitectonic/ Tractographic Label** | **MNI Coordinates** | | |
| --- | --- | --- | --- | --- | --- | --- |
|  |  |  |  | **x** | **y** | **z** |
| 53 | 4.43 | L Posterior-Medial Frontal (pre-SMA) |  | -6 | 10 | 58 |
| 48 | 4.25 | N/A (40) | L Area hIP1 (IPS) | -34 | -48 | 42 |

### Section 10: Contrast results (img> rest_control)

#### Table S71: Meta-analysis of Contrast_Auditory_greater_than_Motor

| **Cluster Voxels** | **Z-score** | **Macroanatomical Location** | **Cytoarchitectonic/ Tractographic Label** | **MNI Coordinates** | | |
| --- | --- | --- | --- | --- | --- | --- |
|  |  |  |  | **x** | **y** | **z** |
| 887 | 6.24 | L Posterior-Medial Frontal (pre-SMA) |  | 0 | 10 | 58 |
|  | 5.77 | L Posterior-Medial Frontal (pre-SMA) |  | -6 | 6 | 62 |
|  | 4.61 | L MCC (SMA) |  | 2 | 16 | 42 |
|  | 4.47 | L MCC (24) |  | -4 | 20 | 38 |
|  | 4.03 | L MCC (pre-SMA) |  | -6 | 10 | 44 |
|  | 3.51 | L Superior Medial Gyrus (pre-SMA) |  | -4 | 22 | 46 |
| 376 | 5.67 | L IFG (p. Opercularis) (48) |  | -40 | 18 | 0 |
|  | 4.13 | L Temporal Pole (38) |  | -52 | 6 | -4 |
|  | 3.83 | N/A (48) |  | -50 | 12 | 0 |
|  | 3.29 | L IFG (p. Triangularis) (S1) | L Area 44 | -50 | 12 | 6 |
| 297 | 5.77 | L IFG (p. Opercularis) (PMv) | L Area 44 | -50 | 10 | 20 |
|  | 3.92 | L IFG (p. Opercularis) (PMv) | L Area 44 | -58 | 4 | 18 |
| 233 | 6.91 | R Precentral Gyrus (PMd) |  | 54 | 2 | 46 |
|  | 3.33 | R Middle Frontal Gyrus (PMd) |  | 42 | 4 | 58 |
| 205 | 5.13 | L Superior Temporal Gyrus (48) |  | -60 | -40 | 24 |
| 195 | 7.01 | L Precentral Gyrus (PMd) |  | -50 | -2 | 50 |
| 142 | 4.69 | L Inferior Parietal Lobule (40) |  | -32 | -52 | 44 |
|  | 4.29 | L Inferior Parietal Lobule (40) |  | -36 | -48 | 42 |
|  | 4.03 | N/A (7) |  | -34 | -58 | 56 |
|  | 3.88 | L Inferior Parietal Lobule (7) |  | -32 | -54 | 50 |

#### Table S72: Meta-analysis of Contrast_Motor_greater_than_Auditory

| **Cluster Voxels** | **Z-score** | **Macroanatomical Location** | **Cytoarchitectonic/ Tractographic Label** | **MNI Coordinates** | | |
| --- | --- | --- | --- | --- | --- | --- |
|  |  |  |  | **x** | **y** | **z** |
| 1822 | 8.13 | N/A (PMd) |  | -24 | -8 | 50 |
|  | 7.73 | L Posterior-Medial Frontal (SMA) |  | -8 | -4 | 54 |
|  | 7.41 | L Posterior-Medial Frontal (SMA) |  | -10 | -6 | 56 |
|  | 2.22 | R Posterior-Medial Frontal (SMA) |  | 10 | -12 | 64 |
| 938 | 8.13 | L Postcentral Gyrus (40) | L Area 2 | -40 | -36 | 48 |
|  | 3.72 | L Inferior Parietal Lobule (40) | L Area 2 | -44 | -42 | 54 |
|  | 2.39 | L SupraMarginal Gyrus | L Area PFt (IPL) | -66 | -32 | 40 |
|  | 2.38 | L SupraMarginal Gyrus (2) | L Area PFt (IPL) | -60 | -26 | 46 |
|  | 2.18 | L SupraMarginal Gyrus | L Area PF (IPL) | -64 | -36 | 42 |
|  | 1.74 | L SupraMarginal Gyrus (2) | L Area PFop (IPL) | -60 | -30 | 30 |
| 501 | 5.40 | L Precuneus (5) |  | -16 | -58 | 62 |
|  | 3.72 | L Precuneus (5) |  | -14 | -60 | 56 |
|  | 3.54 | L Superior Parietal Lobule (7) | L Area 7A (SPL) | -22 | -60 | 64 |
| 430 | 3.43 | L Insula Lobe (48) |  | -44 | -2 | 6 |
|  | 3.40 | L Insula Lobe (48) |  | -38 | 0 | 10 |
|  | 2.54 | N/A (48) |  | -32 | 8 | 8 |
| 420 | 3.16 | R Precentral Gyrus (PMd) |  | 28 | -10 | 60 |
|  | 3.04 | N/A (PMd) |  | 22 | -8 | 54 |
| 286 | 2.88 | R Superior Parietal Lobule (7) | R Area hIP3 (IPS) | 28 | -62 | 56 |
|  | 2.76 | R Superior Parietal Lobule (7) | R Area hIP3 (IPS) | 24 | -64 | 58 |
|  | 2.67 | R Superior Parietal Lobule (7) | R Area 7A (SPL) | 22 | -64 | 64 |
|  | 2.61 | R Superior Parietal Lobule | R Area 7A (SPL) | 18 | -64 | 68 |
|  | 2.38 | R Precuneus (7) |  | 10 | -60 | 64 |
| 244 | 2.99 | R Postcentral Gyrus (S1) | R Area 4a | 42 | -30 | 60 |
|  | 2.85 | R Postcentral Gyrus (S1) | R Area 3b | 44 | -32 | 56 |
| 220 | 3.01 | L IFG (p. Opercularis) (PMv) | L Area 44 | -60 | 10 | 30 |
| 67 | 2.28 | R IFG (p. Opercularis) (PMv) | R Area 44 | 54 | 4 | 30 |
| 27 | 1.99 | R Putamen |  | 30 | -6 | -2 |
|  | 1.88 | N/A (48) |  | 26 | -4 | -4 |

#### Table S73: Meta-analysis of Contrast_Motor_greater_than_Visual

| **Cluster Voxels** | **Z-score** | **Macroanatomical Location** | **Cytoarchitectonic/ Tractographic Label** | **MNI Coordinates** | | |
| --- | --- | --- | --- | --- | --- | --- |
|  |  |  |  | **x** | **y** | **z** |
| 2791 | 8.13 | L Temporal Pole (PMv) | L Area 44 | -54 | 6 | 4 |
|  | 6.78 | L Superior Frontal Gyrus (SMA) |  | -14 | -2 | 60 |
|  | 3.72 | L Posterior-Medial Frontal (pre-SMA) |  | -8 | 6 | 50 |
|  | 3.54 | L Superior Frontal Gyrus (PMd) |  | -16 | 0 | 66 |
|  | 3.44 | L Superior Frontal Gyrus (PMd) |  | -18 | 0 | 70 |
|  | 3.19 | N/A (SMA) |  | -62 | 12 | 14 |
|  | 2.89 | L Precentral Gyrus (M1) |  | -50 | -10 | 54 |
|  | 2.35 | L Precentral Gyrus (PMd) |  | -32 | -6 | 68 |
|  | 2.35 | L Precentral Gyrus (PMd) |  | -34 | -4 | 66 |
|  | 2.19 | L Precentral Gyrus (PMd) |  | -52 | 8 | 44 |
|  | 2.07 | L Precentral Gyrus (PMv) |  | -58 | 2 | 40 |
| 1037 | 3.72 | L Inferior Parietal Lobule (40) |  | -36 | -46 | 54 |
|  | 3.54 | L Inferior Parietal Lobule (40) | L Area hIP3 (IPS) | -38 | -50 | 56 |
|  | 3.29 | L SupraMarginal Gyrus (48) | L Area PFop (IPL) | -58 | -32 | 28 |
|  | 3.19 | L SupraMarginal Gyrus (2) | L Area PFop (IPL) | -66 | -28 | 30 |
|  | 2.51 | L Inferior Parietal Lobule (S1) | L Area PFop (IPL) | -54 | -28 | 50 |
|  | 2.31 | L SupraMarginal Gyrus (2) | L Area PFt (IPL) | -46 | -32 | 38 |
|  | 1.94 | N/A (40) | L Area 1 | -36 | -34 | 40 |
|  | 1.93 | N/A (40) | L Area PFt (IPL) | -32 | -34 | 42 |
| 430 | 3.12 | R Postcentral Gyrus (S1) | R Area 3b | 42 | -26 | 44 |
|  | 3.09 | R Postcentral Gyrus (S1) | R Area 3b | 40 | -28 | 48 |
|  | 2.89 | N/A (S1) | R Area 3a | 36 | -28 | 42 |
|  | 2.59 | R Superior Temporal Gyrus (48) | R Area PFcm (IPL) | 56 | -34 | 26 |
|  | 2.49 | R SupraMarginal Gyrus (48) | R Area OP1 [SII] | 56 | -30 | 24 |
|  | 2.45 | R Postcentral Gyrus (S1) | R Area 3b | 44 | -28 | 56 |
|  | 2.44 | R SupraMarginal Gyrus (48) | R Area PFcm (IPL) | 60 | -36 | 30 |
|  | 2.37 | R SupraMarginal Gyrus (40) | R Area PF (IPL) | 62 | -38 | 34 |
|  | 2.33 | R SupraMarginal Gyrus (40) | R Area PFcm (IPL) | 56 | -34 | 40 |
|  | 2.30 | R Superior Temporal Gyrus (48) | R Area PFcm (IPL) | 54 | -34 | 22 |
|  | 2.21 | R SupraMarginal Gyrus (40) | R Area PF (IPL) | 60 | -36 | 38 |
| 318 | 3.54 | R IFG (p. Opercularis) (PMv) | R Area 44 | 54 | 6 | 12 |
| 288 | 2.89 | R Cerebelum (Crus 1) | R Lobule VI (Hem) | 34 | -58 | -28 |
|  | 2.68 | R Cerebelum (Crus 1) | R Lobule VI (Hem) | 36 | -50 | -30 |
|  | 2.63 | N/A |  | 22 | -58 | -26 |
| 233 | 2.48 | L Putamen |  | -28 | 4 | 8 |
|  | 2.42 | L Putamen (48) |  | -30 | -8 | 6 |
|  | 2.32 | L Pallidum |  | -24 | -10 | 6 |
|  | 1.83 | L Insula Lobe (48) |  | -32 | 14 | 6 |
|  | 1.73 | L Insula Lobe (48) |  | -36 | 16 | 2 |
| 208 | 2.89 | R Precentral Gyrus (PMd) |  | 50 | -4 | 44 |
| 197 | 2.24 | L Cerebelum (VI) | L Lobule VI (Hem) | -24 | -62 | -26 |
|  | 2.22 | L Cerebelum (VI) | L Lobule VI (Hem) | -26 | -58 | -26 |
|  | 2.20 | L Cerebelum (VI) | L Lobule VI (Hem) | -30 | -64 | -22 |
|  | 2.20 | L Cerebelum (VI) | L Lobule VI (Hem) | -32 | -58 | -24 |
|  | 2.04 | L Cerebelum (Crus 1) | L Lobule VIIa crusI (Hem) | -40 | -54 | -26 |
| 136 | 3.01 | R Putamen |  | 28 | -4 | 8 |
|  | 2.83 | R Putamen |  | 30 | -8 | 0 |
| 17 | 1.94 | R Middle Frontal Gyrus (PMd) |  | 40 | -4 | 60 |
|  | 1.80 | R Precentral Gyrus (M1) |  | 34 | -12 | 58 |

#### Table S74: Meta-analysis of Contrast_Visual_greater_than_Motor

| **Cluster Voxels** | **Z-score** | **Macroanatomical Location** | **Cytoarchitectonic/ Tractographic Label** | **MNI Coordinates** | | |
| --- | --- | --- | --- | --- | --- | --- |
|  |  |  |  | **x** | **y** | **z** |
| 666 | 6.00 | L Superior Parietal Lobule (7) |  | -20 | -68 | 52 |
|  | 4.10 | L Inferior Parietal Lobule (7) | L Area hIP3 (IPS) | -30 | -60 | 52 |
|  | 3.96 | L Inferior Parietal Lobule (40) | L Area PFt (IPL) | -42 | -40 | 46 |
|  | 3.88 | L Inferior Parietal Lobule (7) | L Area hIP3 (IPS) | -34 | -58 | 46 |
|  | 3.81 | L Inferior Parietal Lobule (40) | L Area PFt (IPL) | -52 | -42 | 46 |
| 238 | 4.34 | N/A (7) |  | 20 | -64 | 52 |
|  | 2.55 | R Middle Occipital Gyrus (19) |  | 28 | -74 | 36 |
| 167 | 4.87 | R Superior Frontal Gyrus (PMd) |  | 26 | -4 | 60 |
|  | 4.56 | N/A (PMd) |  | 34 | -2 | 50 |
| 133 | 3.90 | R Insula Lobe (48) |  | 46 | 14 | 2 |
|  | 3.72 | N/A |  | 48 | 12 | -2 |
| 127 | 4.59 | L IFG (p. Opercularis) (PMv) | L Area 44 | -48 | 8 | 26 |
| 97 | 2.79 | L Inferior Temporal Gyrus (37) |  | -52 | -60 | -4 |
|  | 2.75 | L Inferior Temporal Gyrus (37) |  | -50 | -56 | -4 |
|  | 2.62 | L Inferior Temporal Gyrus (37) |  | -48 | -58 | -6 |
|  | 2.52 | L Inferior Temporal Gyrus (37) |  | -50 | -50 | -6 |
|  | 2.28 | L Fusiform Gyrus (37) | L Area FG2 | -48 | -60 | -14 |
|  | 2.18 | L Inferior Occipital Gyrus (19) | L Area FG2 | -46 | -68 | -8 |
|  | 2.16 | L Inferior Temporal Gyrus (37) | L Area FG2 | -50 | -66 | -12 |
|  | 2.14 | L Inferior Occipital Gyrus (37) | L Area FG2 | -48 | -64 | -10 |
|  | 2.05 | L Inferior Occipital Gyrus (19) | L Area FG2 | -46 | -72 | -10 |

#### Table S75: Meta-analysis of Contrast_Auditory_greater_than_Visual

| **Cluster Voxels** | **Z-score** | **Macroanatomical Location** | **Cytoarchitectonic/ Tractographic Label** | **MNI Coordinates** | | |
| --- | --- | --- | --- | --- | --- | --- |
|  |  |  |  | **x** | **y** | **z** |
| 582 | 4.31 | R Posterior-Medial Frontal (pre-SMA) |  | 4 | 6 | 64 |
|  | 3.72 | L Posterior-Medial Frontal (pre-SMA) |  | -2 | 10 | 64 |
|  | 3.54 | R Posterior-Medial Frontal (pre-SMA) |  | 8 | 6 | 58 |
|  | 3.06 | L Posterior-Medial Frontal (pre-SMA) |  | -4 | 8 | 54 |
|  | 2.52 | L MCC (24) |  | -2 | 14 | 40 |
|  | 2.47 | L MCC (pre-SMA) |  | -4 | 12 | 44 |
|  | 2.35 | L MCC (pre-SMA) |  | -6 | 10 | 46 |
|  | 2.32 | L MCC (pre-SMA) |  | -10 | 10 | 46 |
| 332 | 5.67 | L IFG (p. Orbitalis) (48) |  | -40 | 18 | 0 |
|  | 2.83 | L Temporal Pole (48) |  | -54 | 10 | 0 |
|  | 2.40 | L IFG (p. Triangularis) (S1) |  | -52 | 12 | 6 |
| 232 | 5.09 | L Superior Temporal Gyrus (48) | L Area PF (IPL) | -60 | -42 | 26 |
| 200 | 3.72 | R Precentral Gyrus (PMd) |  | 54 | 2 | 46 |
| 174 | 5.11 | L Precentral Gyrus (PMd) |  | -54 | -2 | 50 |
|  | 3.72 | L Precentral Gyrus (S1) |  | -48 | -8 | 52 |
|  | 3.54 | L Precentral Gyrus (SMA) |  | -50 | -8 | 48 |
| 150 | 3.35 | L IFG (p. Triangularis) (48) |  | -52 | 16 | 12 |

#### Table S76: Meta-analysis of Contrast_Auditory_greater_than_Gustatory

| **Cluster Voxels** | **Z-score** | **Macroanatomical Location** | **Cytoarchitectonic/ Tractographic Label** | **MNI Coordinates** | | |
| --- | --- | --- | --- | --- | --- | --- |
|  |  |  |  | **x** | **y** | **z** |
| 95 | 3.63 | R MCC (pre-SMA) |  | 4 | 8 | 50 |
|  | 3.54 | L MCC (pre-SMA) |  | -2 | 22 | 40 |
|  | 3.35 | L MCC (32) |  | -6 | 18 | 38 |
|  | 3.01 | L MCC (32) |  | -8 | 10 | 42 |
|  | 2.97 | L MCC (24) |  | -4 | 20 | 34 |
|  | 2.88 | L MCC (24) |  | -2 | 12 | 40 |
|  | 2.85 | R MCC (SMA) |  | 4 | 18 | 42 |
|  | 2.75 | L MCC (pre-SMA) |  | -2 | 8 | 46 |
|  | 2.64 | L Superior Medial Gyrus (pre-SMA) |  | 0 | 12 | 48 |
|  | 2.60 | L Superior Medial Gyrus (pre-SMA) |  | -6 | 12 | 48 |
|  | 2.43 | L MCC (pre-SMA) |  | -8 | 8 | 46 |
| 63 | 3.43 | L SupraMarginal Gyrus (48) | L Area PFcm (IPL) | -56 | -44 | 30 |
|  | 3.35 | L Superior Temporal Gyrus (22) | L Area TE 3 | -64 | -40 | 20 |
|  | 3.24 | L SupraMarginal Gyrus (40) | L Area PF (IPL) | -60 | -46 | 32 |
|  | 3.12 | L Superior Temporal Gyrus (40) | L Area PFcm (IPL) | -58 | -48 | 28 |
|  | 3.04 | L Superior Temporal Gyrus |  | -66 | -42 | 24 |
|  | 2.97 | L SupraMarginal Gyrus (40) | L Area PFm (IPL) | -60 | -52 | 34 |
|  | 2.74 | L Superior Temporal Gyrus | L Area PFcm (IPL) | -58 | -38 | 26 |
|  | 2.67 | L Superior Temporal Gyrus (A1-42) |  | -54 | -44 | 22 |
|  | 2.62 | L Superior Temporal Gyrus (A1-42) | L Area PF (IPL) | -60 | -44 | 26 |
|  | 2.54 | L Superior Temporal Gyrus (A1-42) | L Area PFcm (IPL) | -58 | -38 | 20 |
|  | 2.41 | L Superior Temporal Gyrus (A1-42) |  | -58 | -42 | 20 |
| 21 | 2.73 | N/A (40) | L Area hIP1 (IPS) | -32 | -50 | 42 |
|  | 2.45 | N/A (7) |  | -28 | -50 | 44 |
|  | 2.29 | N/A (40) | L Area hIP1 (IPS) | -38 | -48 | 40 |
|  | 2.21 | L Inferior Parietal Lobule (7) | L Area hIP1 (IPS) | -32 | -56 | 46 |
| 20 | 3.43 | R Precentral Gyrus (PMd) |  | 52 | 4 | 52 |
|  | 2.86 | R Precentral Gyrus (PMd) |  | 46 | 2 | 52 |
|  | 2.54 | R Middle Frontal Gyrus (PMd) |  | 46 | 8 | 54 |
|  | 2.23 | R Middle Frontal Gyrus (PMd) |  | 50 | 8 | 52 |

#### Table S77: Meta-analysis of Contrast_Auditory_greather_than_Tactile

| **Cluster Voxels** | **Z-score** | **Macroanatomical Location** | **Cytoarchitectonic/ Tractographic Label** | **MNI Coordinates** | | |
| --- | --- | --- | --- | --- | --- | --- |
|  |  |  |  | **x** | **y** | **z** |
| 27 | 2.27 | R MCC (24) |  | 2 | 16 | 38 |
|  | 1.85 | R MCC (pre-SMA) |  | 4 | 16 | 44 |

#### Table S78: Meta-analysis of Contrast_Tactile_greather_than_Auditory

| **Cluster Voxels** | **Z-score** | **Macroanatomical Location** | **Cytoarchitectonic/ Tractographic Label** | **MNI Coordinates** | | |
| --- | --- | --- | --- | --- | --- | --- |
|  |  |  |  | **x** | **y** | **z** |
| 98 | 4.43 | L Posterior-Medial Frontal (pre-SMA) |  | -6 | 10 | 58 |
|  | 3.12 | L Posterior-Medial Frontal (pre-SMA) |  | -8 | 12 | 50 |
|  | 2.66 | L Posterior-Medial Frontal (pre-SMA) |  | -6 | 16 | 52 |
|  | 2.37 | L Posterior-Medial Frontal (pre-SMA) |  | -10 | 16 | 52 |
| 80 | 3.54 | N/A (40) | L Area hIP1 (IPS) | -32 | -52 | 42 |

#### Table S79: Meta-analysis of Contrast_Motor_greather_than_Tactile

| **Cluster Voxels** | **Z-score** | **Macroanatomical Location** | **Cytoarchitectonic/ Tractographic Label** | **MNI Coordinates** | | |
| --- | --- | --- | --- | --- | --- | --- |
|  |  |  |  | **x** | **y** | **z** |
| 293 | 3.72 | R Precentral Gyrus (PMv) |  | 58 | 8 | 40 |
|  | 3.29 | R IFG (p. Opercularis) (PMv) | R Area 44 | 56 | 8 | 24 |
|  | 3.26 | R IFG (p. Opercularis) (PMv) | R Area 44 | 52 | 4 | 26 |
|  | 3.24 | R IFG (p. Triangularis) (PMv) | R Area 45 | 56 | 14 | 30 |
|  | 3.12 | R IFG (p. Opercularis) (PMv) | R Area 44 | 56 | 4 | 18 |
|  | 3.04 | R IFG (p. Opercularis) (PMv) | R Area 45 | 54 | 16 | 18 |
|  | 2.93 | R Precentral Gyrus (PMv) |  | 48 | 2 | 36 |
|  | 2.89 | R IFG (p. Opercularis) (PMv) | R Area 44 | 56 | 4 | 22 |
|  | 2.88 | R IFG (p. Opercularis) (PMv) | R Area 45 | 54 | 14 | 34 |
|  | 2.85 | R IFG (p. Opercularis) (PMv) | R Area 44 | 52 | 6 | 22 |
|  | 2.78 | R IFG (p. Opercularis) (PMv) | R Area 44 | 52 | 8 | 16 |
| 174 | 2.88 | L Posterior-Medial Frontal (SMA) |  | -8 | -14 | 68 |
|  | 2.57 | L Posterior-Medial Frontal (SMA) |  | -4 | -14 | 66 |
|  | 2.25 | N/A (SMA) |  | -14 | -10 | 64 |
|  | 1.85 | L Posterior-Medial Frontal (PMd) |  | -16 | -2 | 70 |
| 74 | 3.43 | R Precentral Gyrus (PMd) |  | 26 | -8 | 54 |
|  | 3.12 | R Superior Frontal Gyrus (PMd) |  | 26 | -10 | 60 |
|  | 2.78 | R Superior Frontal Gyrus (PMd) |  | 24 | -8 | 58 |
|  | 2.53 | N/A (PMd) |  | 20 | -8 | 60 |
|  | 2.52 | R Superior Frontal Gyrus (PMd) |  | 22 | -10 | 62 |
|  | 2.49 | R Superior Frontal Gyrus (SMA) |  | 16 | -10 | 68 |
|  | 2.32 | R Superior Frontal Gyrus (PMd) |  | 22 | -6 | 68 |
|  | 2.26 | R Superior Frontal Gyrus (PMd) |  | 20 | -8 | 66 |
|  | 2.23 | R Superior Frontal Gyrus (PMd) |  | 26 | -4 | 68 |
| 24 | 2.89 | R Putamen (48) |  | 32 | -4 | 4 |
|  | 2.48 | N/A |  | 28 | -8 | -2 |
|  | 2.46 | R Putamen (48) |  | 32 | -6 | 0 |
|  | 2.19 | R Putamen |  | 28 | -10 | 2 |
|  | 2.17 | R Putamen (48) |  | 28 | 0 | 4 |
|  | 1.99 | R Putamen (48) |  | 28 | -2 | -4 |

#### Table S80: Meta-analysis of Contrast_Tactile_greather_than_Motor

| **Cluster Voxels** | **Z-score** | **Macroanatomical Location** | **Cytoarchitectonic/ Tractographic Label** | **MNI Coordinates** | | |
| --- | --- | --- | --- | --- | --- | --- |
|  |  |  |  | **x** | **y** | **z** |
| 119 | 4.90 | N/A (40) | L Area hIP1 (IPS) | -34 | -46 | 42 |
| 104 | 5.32 | L Posterior-Medial Frontal (pre-SMA) |  | -8 | 12 | 56 |

#### Table S81: Meta-analysis of Contrast_Tactile_greather_than_Visual

| **Cluster Voxels** | **Z-score** | **Macroanatomical Location** | **Cytoarchitectonic/ Tractographic Label** | **MNI Coordinates** | | |
| --- | --- | --- | --- | --- | --- | --- |
|  |  |  |  | **x** | **y** | **z** |
| 119 | 4.22 | N/A (40) | L Area hIP1 (IPS) | -34 | -48 | 40 |
| 103 | 3.54 | L Posterior-Medial Frontal (pre-SMA) |  | -6 | 10 | 52 |

#### Table S82: Meta-analysis of Contrast_Tactile_greather_than_Gustatory

| **Cluster Voxels** | **Z-score** | **Macroanatomical Location** | **Cytoarchitectonic/ Tractographic Label** | **MNI Coordinates** | | |
| --- | --- | --- | --- | --- | --- | --- |
|  |  |  |  | **x** | **y** | **z** |
| 119 | 4.67 | N/A (40) | L Area hIP1 (IPS) | -34 | -48 | 42 |
|  | 3.95 | N/A (40) |  | -32 | -42 | 38 |
|  | 3.87 | N/A (40) |  | -34 | -46 | 38 |
|  | 3.85 | L Inferior Parietal Lobule (40) | L Area hIP1 (IPS) | -34 | -46 | 46 |
|  | 3.06 | N/A (40) |  | -32 | -44 | 42 |

### Section 11: Sub-analyses Visual Imagery

#### Table S83: Meta-analysis of Visual Motion Imagery (n=25)

| **Cluster Voxels** | **Z-score** | **Macroanatomical Location** | **Cytoarchitectonic/ Tractographic Label** | **MNI Coordinates** | | |
| --- | --- | --- | --- | --- | --- | --- |
|  |  |  |  | **x** | **y** | **z** |
| 471 | 5.75 | N/A (PMd) |  | -26 | -6 | 52 |
| 193 | 6.18 | R Middle Frontal Gyrus (PMd) |  | 28 | -4 | 58 |
| 127 | 4.26 | L IFG (p. Triangularis) (PMv) | L Area 45 | -54 | 12 | 32 |
|  | 3.87 | L IFG (p. Opercularis) (PMv) | L Area 44 | -46 | 2 | 32 |
|  | 3.75 | L Precentral Gyrus (PMv) | L Area 44 | -50 | 2 | 28 |
|  | 3.34 | L Precentral Gyrus (PMv) |  | -44 | 0 | 38 |
| 103 | 4.88 | R SupraMarginal Gyrus (40) | R Area 2 | 44 | -38 | 46 |

#### Table S84: Meta-analysis of Visual Form Imagery (n=53)

| **Cluster Voxels** | **Z-score** | **Macroanatomical Location** | **Cytoarchitectonic/ Tractographic Label** | **MNI Coordinates** | | |
| --- | --- | --- | --- | --- | --- | --- |
|  |  |  |  | **x** | **y** | **z** |
| 283 | 5.55 | L Superior Parietal Lobule |  | -22 | -68 | 50 |
|  | 4.42 | L Inferior Parietal Lobule (7) | L Area hIP3 (IPS) | -30 | -60 | 48 |
|  | 3.63 | L Inferior Parietal Lobule (40) | L Area Hip2 (IPS) | -52 | -44 | 46 |
|  | 3.57 | L Inferior Parietal Lobule (40) | L Area PFt (IPL) | -42 | -40 | 46 |
| 453 | 6.08 | L Inferior Temporal Gyrus (37) |  | -52 | -58 | -8 |
|  | 3.66 | L Inferior Occipital Gyrus (19) | L Area FG2 | -46 | -70 | -12 |
|  | 3.60 | L Fusiform Gyrus (37) | L Area FG3 | -34 | -52 | -14 |
| 243 | 4.85 | R Superior Occipital Gyrus (7) |  | 24 | -72 | 40 |
|  | 3.48 | N/A (7) |  | 22 | -62 | 50 |
| 199 | 5.41 | L IFG (p. Opercularis) (PMv) |  | -46 | 8 | 28 |
| 177 | 5.13 | R Inferior Temporal Gyrus (37) |  | 52 | -56 | -10 |
| 125 | 4.20 | R MCC (pre-SMA) |  | 2 | 22 | 42 |
|  | 3.72 | L Posterior-Medial Frontal (pre-SMA) |  | -4 | 18 | 52 |
| 110 | 4.22 | L Precentral Gyrus (PMd) |  | -32 | 2 | 48 |

### Section 12: List of included papers

#### Table S85: Included papers

| **Author, year** | **Imagery** | **Subject** | **Task** | **Foci** | **DOI** |
| --- | --- | --- | --- | --- | --- |
| Abidi,2021 | Motor | 14 | Imagine walking | 2 | 10.1002/jmri.27335 |
| Abidi,2021 | Motor | 14 | Imagine walking | 3 | 10.1002/jmri.27335 |
| Alkadhi,2005 | Motor | 8 | Imagine leg movement | 19 | 10.1093/cercor/bhh116 |
| Allali,2014 | Motor | 14 | Imagine leg movement | 46 | 10.1093/gerona/glt207 |
| Amemiya,2016 | Motor | 19 | Imagine arm movement | 13 | 10.1016/j.cortex.2016.01.017 |
| Amemiya,2021 | Motor | 16 | Imagine walking | 27 | 10.1007/s11682-020-00275-w |
| Baeck,2012 | Motor | 18 | Imagine whole body movement | 14 | 10.1016/j.bbr.2012.06.001 |
| Bagarinao,2018 | Motor | 22 | Imagine hand movement | 15 | 10.3389/fnhum.2018.00158 |
| Bagarinao,2020 | Motor | 30 | Imagine hand movement | 10 | 10.3389/fnins.2020.00623 |
| Bagarinao,2020 | Motor | 30 | Imagine hand movement | 21 | 10.3389/fnins.2020.00623 |
| Bagarinao,2020 | Motor | 30 | Imagine hand movement | 18 | 10.3389/fnins.2020.00623 |
| Bakker,2008 | Motor | 16 | Imagine leg movement | 9 | 10.1016/j.neuroimage.2008.03.020 |
| Bastepe-Gray,2020 | Motor | 7 | Imagine finger movement | 7 | 10.1016/j.jchemneu.2020.101748 |
| Bastepe-Gray,2020 | Motor | 7 | Imagine finger movement | 4 | 10.1016/j.jchemneu.2020.101748 |
| Bastepe-Gray,2020 | Motor | 7 | Imagine finger movement | 6 | 10.1016/j.jchemneu.2020.101748 |
| Bastepe-Gray,2020 | Motor | 7 | Imagine finger movement | 12 | 10.1016/j.jchemneu.2020.101748 |
| Bastepe-Gray,2020 | Motor | 7 | Imagine finger movement | 12 | 10.1016/j.jchemneu.2020.101748 |
| Bastepe-Gray,2020 | Motor | 7 | Imagine finger movement | 2 | 10.1016/j.jchemneu.2020.101748 |
| Baumann,2022 | Motor | 18 | Imagine writing | 4 | 10.3389/fnhum.2022.829576 |
| Baumann,2022 | Motor | 18 | Imagine writing | 2 | 10.3389/fnhum.2022.829576 |
| Baumann,2022 | Motor | 18 | Imagine writing | 4 | 10.3389/fnhum.2022.829576 |
| Berman,2012 | Motor | 15 | Imagine arm movement | 25 | 10.1016/j.neuroimage.2011.07.035 |
| Beudel,2011 | Motor | 7 | Imagine leg movement | 16 | 10.1002/hbm.21044 |
| Beudel,2011 | Motor | 7 | Imagine leg movement | 29 | 10.1002/hbm.21044 |
| Bezzola,2012 | Motor | 22 | Imagine whole body movement | 4 | 10.3389/fnhum.2012.00067 |
| Bhatt,2018 | Motor | 10 | Imagine walking | 8 | 10.3389/fnbeh.2018.00203 |
| Bhatt,2018 | Motor | 10 | Imagine walking | 1 | 10.3389/fnbeh.2018.00203 |
| Bhatt,2018 | Motor | 10 | Imagine walking | 10 | 10.3389/fnbeh.2018.00203 |
| Binkofski,2000 | Motor | 6 | Imagine arm movement | 14 | 10.1002/1097-0193(200012)11:4<273::aid-hbm40>3.0.co;2-0 |
| Blumen,2014 | Motor | 33 | Imagine leg movement | 9 | 10.1002/hbm.22461 |
| Boecker,2002 | Motor | 6 | Imagine arm movement | 11 | 10.1006/nimg.2002.1139 |
| Boly,2007 | Motor | 12 | Imagine faces | 24 | 10.1016/j.neuroimage.2007.02.047 |
| Bonda,1995 | Motor | 16 | Imagine arm movement | 21 | 10.1073/pnas.92.24.11180 |
| Bonda,1996 | Motor | 14 | Imagine arm movement | 14 | 10.1152/jn.1996.76.3.2042 |
| Boraxbekk,2016 | Motor | 56 | Imagine finger movement | 7 | 10.1016/j.neuropsychologia.2016.07.019 |
| Burianova,2013 | Motor | 14 | Imagine arm movement | 11 | 10.1016/j.neuroimage.2013.01.001 |
| Chang,2011 | Motor | 18 | Imagine whole body movement | 18 | 10.1002/nbm.1600 |
| Corradi-Dell'Acqua,2009 | Motor | 17 | Imagine hand rotation | 6 | 10.1523/JNEUROSCI.4861-08.2009 |
| Corradi-Dell'Acqua,2009 | Motor | 17 | Imagine hand rotation | 11 | 10.1523/JNEUROSCI.4861-08.2009 |
| Creem,2001 | Motor | 12 | Imagine whole body movement | 15 | 10.3758/cabn.1.3.239 |
| Creem,2001 | Motor | 12 | Imagine whole body movement | 2 | 10.3758/cabn.1.3.239 |
| Creem-Regehr,2007 | Motor | 13 | Imagine grasping movement | 18 | 10.1017/S1355617707071093 |
| Creem-Regehr,2007 | Motor | 13 | Imagine grasping movement | 26 | 10.1017/S1355617707071093 |
| Creem-Regehr,2007 | Motor | 13 | Imagine grasping movement | 9 | 10.1017/S1355617707071093 |
| Creem-Regehr,2007b | Motor | 13 | Imagine arm movement | 3 | 10.1016/j.neuroimage.2006.11.057 |
| Creem-Regehr,2007b | Motor | 13 | Imagine arm movement | 7 | 10.1016/j.neuroimage.2006.11.057 |
| Creem-Regehr,2007b | Motor | 13 | Imagine arm movement | 14 | 10.1016/j.neuroimage.2006.11.057 |
| Crémers,2012 | Motor | 18 | Imagine leg movement | 39 | 10.1002/hbm.21255 |
| Crotti,2022 | Motor | 51 | Imagine squeezing movement | 6 | 10.1002/jnr.25003 |
| Crotti,2022 | Motor | 51 | Imagine squeezing movement | 4 | 10.1002/jnr.25003 |
| de Lange,2005 | Motor | 6 | Imagine arm movement | 7 | 10.1162/0898929052880039 |
| de Lange,2005 | Motor | 17 | Imagine arm movement | 4 | 10.1162/0898929052880039 |
| de Lange,2005 | Motor | 17 | Imagine arm movement | 6 | 10.1162/0898929052880039 |
| de Vries,2009 | Motor | 10 | Imagine arm movement | 12 | 10.1016/j.brainres.2009.06.006 |
| Decety,1994 | Motor | 6 | Imagine arm movement | 86 | 10.1038/371600a0 |
| Deen,2010 | Motor | 15 | Imagine whole body movement | 13 | 10.1016/j.neuropsychologia.2010.01.028 |
| Dieber,1998 | Motor | 10 | Imagine arm movement | 8 | 10.1006/nimg.1997.0314 |
| Diers,2015 | Motor | 20 | Imagine hand movement | 17 | 10.1016/j.brainres.2014.11.001 |
| Enzinger,2008 | Motor | 5 | Imagine leg and arm movement | 20 | 10.1007/s00221-008-1465-y |
| Errante,2019 | Motor | 12 | Imagine grasping movement | 9 | 10.3389/fneur.2019.00837 |
| Ferraye,2014 | Motor | 20 | Imagine whole body movement | 14 | 10.1371/journal.pone.0091183 |
| Flanagin,2009 | Motor | 9 | Imagine leg or whole body movement | 14 | 10.1111/j.1749-6632.2009.03844.x |
| Fleming,2010 | Motor | 15 | Imagine arm movement | 35 | 10.1007/s00221-009-2062-4 |
| Geers,2021 | Motor | 30 | Imagine grasping movement | 10 | 10.1038/s41598-021-86719-9 |
| Gerardin,2000 | Motor | 8 | Imagine arm movement | 20 | 10.1093/cercor/10.11.1093 |
| Gerardin,2000 | Motor | 8 | Imagine arm movement | 15 | 10.1093/cercor/10.11.1093 |
| Grafton,1996 | Motor | 7 | Imagine arm movement | 12 | 10.1007/BF00227183 |
| Gu,2021 | Motor | 16 | Imagine hand movement | 29 | 10.3233/NRE-210185 |
| Gu,2021 | Motor | 16 | Imagine hand movement | 44 | 10.3233/NRE-210185 |
| Guillot,2008 | Motor | 13 | Imagine arm movement | 35 | 10.1016/j.neuroimage.2008.03.042 |
| Guillot,2008 | Motor | 15 | Imagine arm movement | 26 | 10.1016/j.neuroimage.2008.03.042 |
| Guillot,2008 | Motor | 13 | Imagine arm movement | 10 | 10.1016/j.neuroimage.2008.03.042 |
| Guillot,2008 | Motor | 15 | Imagine arm movement | 4 | 10.1016/j.neuroimage.2008.03.042 |
| Guillot,2009 | Motor | 13 | Imagine arm movement | 87 | 10.1002/hbm.20658 |
| Guillot,2009 | Motor | 13 | Imagine arm movement | 18 | 10.1002/hbm.20658 |
| Halder,2011 | Motor | 8 | Imagine limbs movement | 25 | 10.1016/j.neuroimage.2011.01.021 |
| Halder,2011 | Motor | 9 | Imagine limbs movement | 24 | 10.1016/j.neuroimage.2011.01.021 |
| Hamada,2018 | Motor | 26 | Imagine hand rotation | 10 | 10.1007/s11682-017-9821-9 |
| Hamada,2018 | Motor | 26 | Imagine hand rotation | 13 | 10.1007/s11682-017-9821-9 |
| Hanakawa,2008 | Motor | 13 | Imagine arm movement | 12 | 10.1093/cercor/bhn036 |
| Hanakawa,2017 | Motor | 38 | Imagine finger movement | 10 | 10.1523/ENEURO.0200-17.2017 |
| Hanawaka,2003 | Motor | 10 | Imagine finger movement | 3 | 10.1152/jn.00132.2002 |
| Harrigton,2007 | Motor | 11 | Imagine arm movement | 35 | 10.1002/hbm.20286 |
| Harrigton,2007 | Motor | 11 | Imagine arm movement | 16 | 10.1002/hbm.20286 |
| Harrington,2009 | Motor | 8 | Imagine arm movement | 38 | 10.1016/j.cortex.2007.10.015 |
| Harrington,2009 | Motor | 8 | Imagine arm movement | 8 | 10.1016/j.cortex.2007.10.015 |
| Harris,2014 | Motor | 12 | Imagine arm movement | 7 | 10.1371/journal.pone.0093681 |
| He,2021 | Motor | 20 | Imagine drawing | 12 | 10.3389/fnhum.2021.706425 |
| He,2021 | Motor | 20 | Imagine drawing | 6 | 10.3389/fnhum.2021.706425 |
| Herholz,2016 | Motor | 14 | Imagine arm movement | 52 | 10.1093/cercor/bhv138 |
| Hernandez-Martin,2020 | Motor | 8 | Imagine finger movement | 11 | 10.1364/BOE.399907 |
| Higuchi,2007 | Motor | 8 | Imagine arm movement | 10 | 10.1016/s0010-9452(08)70460-x |
| Ionta,2010 | Motor | 12 | Imagine leg movement | 5 | 10.1002/hbm.20898 |
| Iseki,2008 | Motor | 16 | Imagine leg movement | 11 | 10.1016/j.neuroimage.2008.03.010 |
| Jahn,2004 | Motor | 13 | Imagine leg movement | 13 | 10.1016/j.neuroimage.2004.05.017 |
| Jahn,2004 | Motor | 13 | Imagine leg movement | 17 | 10.1016/j.neuroimage.2004.05.017 |
| Jahn,2004 | Motor | 13 | Imagine leg movement | 11 | 10.1016/j.neuroimage.2004.05.017 |
| Jahn,2008 | Motor | 26 | Imagine whole body movement | 21 | 10.1016/j.neuroimage.2007.09.047 |
| Jahn,2008 | Motor | 26 | Imagine whole body movement | 21 | 10.1016/j.neuroimage.2007.09.047 |
| Jahn,2008 | Motor | 26 | Imagine whole body movement | 31 | 10.1016/j.neuroimage.2007.09.047 |
| Jiang,2015 | Motor | 11 | Imagine leg movement | 35 | 10.1016/j.bandc.2015.04.005 |
| Jiang,2015 | Motor | 11 | Imagine leg movement | 90 | 10.1016/j.bandc.2015.04.005 |
| Johnson,2002 | Motor | 8 | Imagine arm movement | 7 | 10.1006/nimg.2002.1265 |
| Johnson,2002 | Motor | 8 | Imagine arm movement | 13 | 10.1006/nimg.2002.1265 |
| Johnson,2012 | Motor | 12 | Imagine arm movement | 12 | 10.1111/j.1552-6569.2010.00529.x |
| Johnson,2012 | Motor | 12 | Imagine arm movement | 24 | 10.1111/j.1552-6569.2010.00529.x |
| Johnson,2012 | Motor | 12 | Imagine arm movement | 24 | 10.1111/j.1552-6569.2010.00529.x |
| Kilintari,2016 | Motor | 14 | Imagine arm movement | 23 | 10.1016/j.brainres.2016.06.009 |
| Kilintari,2016 | Motor | 14 | Imagine arm movement | 12 | 10.1016/j.brainres.2016.06.009 |
| Kim,2023 | Motor | 18 | Imagine grasping movement | 3 | 10.1007/s10548-023-00956-x |
| Kim,2023 | Motor | 18 | Imagine grasping movement | 5 | 10.1007/s10548-023-00956-x |
| Klaus,2020 | Motor | 20 | Imagine self rotation | 7 | 10.1162/jocn_a_01496 |
| Klaus,2020 | Motor | 20 | Imagine self rotation | 8 | 10.1162/jocn_a_01496 |
| Klaus,2020 | Motor | 20 | Imagine self rotation | 3 | 10.1162/jocn_a_01496 |
| Klaus,2020 | Motor | 20 | Imagine self rotation | 4 | 10.1162/jocn_a_01496 |
| Kober,2019 | Motor | 11 | Imagine swallowing | 6 | 10.1007/s00455-019-09985-w |
| Kober,2019 | Motor | 11 | Imagine swallowing | 4 | 10.1007/s00455-019-09985-w |
| Kosslyn,1998 | Motor | 12 | Imagine arm movement | 9 | 10.1111/1469-8986.3520151 |
| Kosslyn,1998 | Motor | 12 | Imagine arm movement | 4 | 10.1111/1469-8986.3520151 |
| Kraeutner,2020 | Motor | 38 | Imagine dart trowing | 13 | 10.1038/s41598-020-78120-9 |
| Kraft,2015 | Motor | 15 | Imagine grip force task | 4 | 10.3233/NRE-151221 |
| Kuhtz-Buschbeck,2003 | Motor | 12 | Imagine arm movement | 24 | 10.1111/j.1460-9568.2003.03066.x |
| Kuhtz-Buschbeck,2003 | Motor | 12 | Imagine arm movement | 18 | 10.1111/j.1460-9568.2003.03066.x |
| Kurby,2013 | Motor | 28 | Imagine movement | 2 | 10.1016/j.bandl.2013.07.003 |
| Kurby,2013 | Motor | 28 | Imagine movement | 1 | 10.1016/j.bandl.2013.07.003 |
| la Fougère,2010 | Motor | 16 | Imagine leg movement | 20 | 10.1016/j.neuroimage.2009.12.060 |
| Labriffe,2017 | Motor | 18 | Imagine leg movement | 19 | 10.3389/fnhum.2017.00106 |
| Lacourse,2005 | Motor | 54 | Imagine arm movement | 15 | 10.1016/j.neuroimage.2005.04.025 |
| Lafleur,2002 | Motor | 9 | Imagine leg movement | 12 | 10.1006/nimg.2001.1048 |
| Lafleur,2002 | Motor | 9 | Imagine leg movement | 6 | 10.1006/nimg.2001.1048 |
| Lebon,2018 | Motor | 48 | Imagine finger movement | 3 | 10.1002/hbm.23956 |
| Lebon,2018 | Motor | 48 | Imagine finger movement | 4 | 10.1002/hbm.23956 |
| Lee,2019 | Motor | 18 | Imagine grasping movement | 3 | 10.1038/s41598-019-49254-2 |
| Lee,2019 | Motor | 18 | Imagine grasping movement | 5 | 10.1038/s41598-019-49254-2 |
| Lee,2019 | Motor | 18 | Imagine grasping movement | 10 | 10.1038/s41598-019-49254-2 |
| Lorey,2009 | Motor | 20 | Imagine arm movement | 15 | 10.1007/s00221-008-1693-1 |
| Lorey,2009 | Motor | 20 | Imagine arm movement | 6 | 10.1007/s00221-008-1693-1 |
| Lorey,2009 | Motor | 20 | Imagine arm movement | 11 | 10.1007/s00221-008-1693-1 |
| Lorey,2010 | Motor | 23 | Imagine arm movement | 13 | 10.1016/j.neuroimage.2009.11.038 |
| Lorey,2010 | Motor | 23 | Imagine arm movement | 17 | 10.1016/j.neuroimage.2009.11.038 |
| Lorey,2011 | Motor | 23 | Imagine arm movement | 21 | 10.1371/journal.pone.0020368 |
| Lorey,2014 | Motor | 18 | Imagine hand and foot movement | 22 | 10.1002/hbm.22246 |
| Lorey,2014 | Motor | 18 | Imagine hand and foot movement | 17 | 10.1002/hbm.22246 |
| Lu,2016 | Motor | 10 | Imagine arm movement | 12 | 10.4103/1673-5374.180756 |
| Lui,2018 | Motor | 16 | Imagine arm movement | 14 | 10.1080/17470910701458551 |
| Ma,2022 | Motor | 21 | Imagine hand movement | 30 | 10.3389/fnins.2022.806406 |
| Madkhali,2022 | Motor | 4 | Imagine arm movement | 13 | 10.1155/2022/8744982 |
| Maidan,2016 | Motor | 20 | Imagine leg movement | 17 | 10.1016/j.parkreldis.2016.01.025 |
| Maillet,2015 | Motor | 8 | Imagine leg movement | 20 | 10.1002/hbm.22679 |
| Malouin,2003 | Motor | 6 | Imagine leg and whole body movement | 37 | 10.1002/hbm.10103 |
| Malouin,2003 | Motor | 6 | Imagine leg and whole body movement | 2 | 10.1002/hbm.10103 |
| Malouin,2003 | Motor | 6 | Imagine leg and whole body movement | 5 | 10.1002/hbm.10103 |
| Marchesotti,2017 | Motor | 12 | Imagine arm movement | 22 | 10.1002/hbm.23566 |
| Marins,2015 | Motor | 14 | Imagine arm movement | 21 | 10.3389/fnbeh.2015.00341 |
| Meister,2004 | Motor | 12 | Imagine arm movement | 12 | 10.1016/j.cogbrainres.2003.12.005 |
| Mizuguchi,2013 | Motor | 19 | Imagine arm movement | 13 | 10.1016/j.neures.2013.03.012 |
| Mizuguchi,2013 | Motor | 19 | Imagine arm movement | 18 | 10.1016/j.neures.2013.03.012 |
| Mizuguchi,2013 | Motor | 19 | Imagine arm movement | 6 | 10.1016/j.neures.2013.03.012 |
| Mizuguchi,2014 | Motor | 16 | Imagine arm movement | 8 | 10.3389/fnhum.2014.00810 |
| Mizuguchi,2014 | Motor | 16 | Imagine arm movement | 10 | 10.3389/fnhum.2014.00810 |
| Mizuguchi,2014 | Motor | 16 | Imagine arm movement | 8 | 10.3389/fnhum.2014.00810 |
| Mizuguchi,2014b | Motor | 17 | Imagine hand and foot movement | 49 | 10.1016/j.neulet.2014.08.025 |
| Mizuguchi,2016 | Motor | 19 | Imagine whole body movement | 13 | 10.1016/j.neuroscience.2015.12.013 |
| Mizuguchi,2019 | Motor | 34 | Imagine finger movement | 23 | 10.1016/j.neures.2018.12.005 |
| Mochizuki,2014 | Motor | 30 | Imagine coordination exercices | 7 | 10.1080/17461391.2014.893019 |
| Mochizuki,2014 | Motor | 30 | Imagine coordination exercices | 3 | 10.1080/17461391.2014.893019 |
| Müller,2012 | Motor | 16 | Imagine arm movement | 4 | 10.1016/j.bbr.2012.03.013 |
| Müller,2012 | Motor | 16 | Imagine arm movement | 6 | 10.1016/j.bbr.2012.03.013 |
| Müller,2013 | Motor | 16 | Imagine arm movement | 6 | 10.1016/j.neuroimage.2012.10.073 |
| Müller,2013 | Motor | 16 | Imagine arm movement | 4 | 10.1016/j.neuroimage.2012.10.073 |
| Munzert,2008 | Motor | 10 | Imagine movement sequences | 5 | 10.1007/s00221-008-1376-y |
| Naito,2002 | Motor | 10 | Imagine arm movement | 8 | 10.1523/JNEUROSCI.22-09-03683.2002 |
| Naito,2002 | Motor | 10 | Imagine arm movement | 4 | 10.1523/JNEUROSCI.22-09-03683.2002 |
| Norton,2023 | Motor | 14 | Imagine playing tennis | 3 | 10.1016/j.ijchp.2022.100347 |
| Norton,2023 | Motor | 14 | Imagine playing tennis | 13 | 10.1016/j.ijchp.2022.100347 |
| Nyberg,2006 | Motor | 16 | Imagine finger movement | 12 | 10.1016/j.neuropsychologia.2005.08.006 |
| Olivetti,2009 | Motor | 9 | Imagine whole body movement | 8 | 10.1016/j.actpsy.2009.06.009 |
| Olivetti,2009 | Motor | 9 | Imagine whole body movement | 7 | 10.1016/j.actpsy.2009.06.009 |
| Olivetti,2009 | Motor | 9 | Imagine whole body movement | 4 | 10.1016/j.actpsy.2009.06.009 |
| Olsson,2008 | Motor | 12 | Imagine whole body movement | 18 | 10.2174/1874440000802010005 |
| Olsson,2012 | Motor | 8 | Imagine whole body movement | 8 | 10.1016/j.expneurol.2012.03.022 |
| Olsson,2012b | Motor | 18 | Imagine arm movement | 9 | 10.3389/fnhum.2012.00255 |
| Olsson,2012b | Motor | 18 | Imagine arm movement | 2 | 10.3389/fnhum.2012.00255 |
| Onuki,2015 | Motor | 16 | Imagine arm movement | 10 | 10.1093/cercor/bht221 |
| Osborne,2015 | Motor | 15 | Imagine arm movement | 3 | 10.3389/fnhum.2015.00493 |
| Oullier,2005 | Motor | 15 | Imagine arm movement | 21 | 10.1093/cercor/bhh198 |
| Oullier,2005 | Motor | 15 | Imagine arm movement | 12 | 10.1093/cercor/bhh198 |
| Palmiero,2009 | Motor | 9 | Imagine whole body movement | 6 | 10.1007/s10339-009-0324-5 |
| Papageorgiou,2013 | Motor | 24 | Imagine face movement | 10 | 10.1073/pnas.1210738110 |
| Patel,2018 | Motor | 10 | Imagine slipping, walking | 10 | 10.3389/fneur.2018.01181 |
| Patel,2018 | Motor | 10 | Imagine slipping, walking | 7 | 10.3389/fneur.2018.01181 |
| Piefke,2009 | Motor | 14 | Imagine face or hand movement | 8 | 10.1002/hbm.20514 |
| Pilgramm,2013 | Motor | 20 | Imagine sueezing or pointing movement | 12 | 10.1002/hbm.23015 |
| Romero-Romo,2010 | Motor | 6 | Imagine leg movement | 8 | 10.1177/197140091002300605 |
| Rousseau,2021 | Motor | 26 | Imagine hand movement | 11 | 10.1093/cercor/bhaa376 |
| Ruby,2001 | Motor | 10 | Imagine arm movement | 9 | 10.1038/87510 |
| Ruby,2001 | Motor | 10 | Imagine arm movement | 7 | 10.1038/87510 |
| Ruby,2001 | Motor | 10 | Imagine arm movement | 6 | 10.1038/87510 |
| Ruffieux,2018 | Motor | 15 | Imagine standing or balance exercices | 6 | 10.3389/fnbeh.2018.00010 |
| Ruppert-Junck,2023 | Motor | 10 | Imagine walking | 3 | 10.1016/j.heliyon.2023.e14741 |
| Sacco,2006 | Motor | 10 | Imagine leg movement | 29 | 10.1016/j.neuroimage.2006.05.018 |
| Sacco,2006 | Motor | 10 | Imagine leg movement | 9 | 10.1016/j.neuroimage.2006.05.018 |
| Sacheli,2017 | Motor | 24 | Imagine walking | 50 | 10.1002/hbm.23725 |
| Sacheli,2020 | Motor | 43 | Imagine walking | 28 | 10.1002/hbm.24919 |
| Sakreida,2018 | Motor | 28 | Imagine arm movement | 23 | 10.1093/cercor/bhw414 |
| Sauvage,2011 | Motor | 8 | Imagine arm movement | 16 | 10.1007/s11682-011-9118-3 |
| Sauvage,2013 | Motor | 12 | Imagine leg movement | 57 | 10.1016/j.neurad.2012.10.001 |
| Sauvage,2013 | Motor | 12 | Imagine leg movement | 5 | 10.1016/j.neurad.2012.10.001 |
| Sauvage,2015 | Motor | 35 | Imagine leg movement | 26 | 10.1016/j.neurad.2014.04.001 |
| Seiler,2015 | Motor | 18 | Imagine arm movement | 28 | 10.1123/jsep.2014-0303 |
| Seiler,2022 | Motor | 16 | Imagine arm and trunc movement | 38 | 10.1123/jsep.2021-0229 |
| Seiler,2022 | Motor | 16 | Imagine arm and trunc movement | 35 | 10.1123/jsep.2021-0229 |
| Seitz,1997 | Motor | 8 | Imagine arm movement | 6 | 10.1111/j.1460-9568.1997.tb01407.x |
| Servos,2002 | Motor | 10 | Imagine arm movement | 2 | 10.1093/cercor/12.7.772 |
| Sharma,2009 | Motor | 13 | Imagine arm movement | 14 | 10.1002/ana.21810 |
| Sharma,2014 | Motor | 31 | Imagine arm movement | 28 | 10.1371/journal.pone.0088443 |
| Simmonds,2014 | Motor | 17 | Imagine face movement | 24 | 10.1523/JNEUROSCI.0336-14.2014 |
| Simos,2017 | Motor | 21 | Imagine arm movement | 29 | 10.1016/j.neuroimage.2017.03.036 |
| Snijders,2011 | Motor | 21 | Imagine leg movement | 6 | 10.1093/brain/awq324 |
| Snijders,2011 | Motor | 21 | Imagine leg movement | 20 | 10.1093/brain/awq324 |
| Soliman,2017 | Motor | 34 | Imagine face movement | 44 | 10.1097/WNR.0000000000000758 |
| Stephan,1995 | Motor | 6 | Imagine arm movement | 25 | 10.1152/jn.1995.73.1.373 |
| Stephan,1995 | Motor | 6 | Imagine arm movement | 3 | 10.1152/jn.1995.73.1.373 |
| Szameitat,2007 | Motor | 15 | Imagine arm and whole body movement | 11 | 10.1016/j.neuroimage.2006.09.033 |
| Szameitat,2007 | Motor | 15 | Imagine arm and whole body movement | 12 | 10.1016/j.neuroimage.2006.09.033 |
| Szameitat,2007 | Motor | 15 | Imagine arm and whole body movement | 13 | 10.1016/j.neuroimage.2006.09.033 |
| Szameitat,2007 | Motor | 15 | Imagine arm and whole body movement | 5 | 10.1016/j.neuroimage.2006.09.033 |
| Szameitat,2007 | Motor | 15 | Imagine arm and whole body movement | 7 | 10.1016/j.neuroimage.2006.09.033 |
| Szameitat,2007b | Motor | 17 | Imagine arm movement | 47 | 10.1111/j.1460-9568.2007.05920.x |
| Szameitat,2012 | Motor | 17 | Imagine arm movement | 11 | 10.1371/journal.pone.0038506 |
| Szameitat,2012b | Motor | 21 | Imagine arm movement | 8 | 10.1016/j.neuroimage.2012.05.009 |
| Tanaka,2017 | Motor | 41 | Imagine playing music | 7 | 10.3389/fnhum.2017.00606 |
| Taube,2015 | Motor | 16 | Imagine balance or static exercices | 17 | 10.1016/j.cortex.2014.09.022 |
| Taube,2015 | Motor | 16 | Imagine balance or static exercices | 27 | 10.1016/j.cortex.2014.09.022 |
| Tian,2016 | Motor | 18 | Imagine face movement | 11 | 10.1016/j.cortex.2016.01.002 |
| Tomasino,2007 | Motor | 15 | Imagine whole body movement | 10 | 10.1016/j.neuroimage.2007.03.035 |
| Tomasino,2013 | Motor | 20 | Imagine whole body movement | 16 | 10.1016/j.brainres.2013.09.048 |
| Tomasino,2013 | Motor | 20 | Imagine whole body movement | 37 | 10.1016/j.brainres.2013.09.048 |
| Ueno,2010 | Motor | 15 | Imagine arm movement | 8 | 10.1007/s11682-009-9087-y |
| van der Meulen,2014 | Motor | 20 | Imagine leg movement | 34 | 10.1002/hbm.22192 |
| van Elk,2012 | Motor | 18 | Imagine arm movement | 8 | 10.1007/s00221-012-3016-9 |
| Villiger,2013 | Motor | 14 | Imagine foot movement | 12 | 10.1371/journal.pone.0072403 |
| Villiger,2013 | Motor | 14 | Imagine foot movement | 8 | 10.1371/journal.pone.0072403 |
| Vingerhoets,2002 | Motor | 12 | Imagine arm movement | 6 | 10.1006/nimg.2002.1290 |
| Vrana,2015 | Motor | 14 | Imagine daily life movement | 13 | 10.1371/journal.pone.0142391 |
| Vrana,2015 | Motor | 14 | Imagine daily life movement | 1 | 10.1371/journal.pone.0142391 |
| Vry,2012 | Motor | 20 | Imagine wrist movement | 9 | 10.1007/s00221-012-3079-7 |
| Wagner,2008 | Motor | 12 | Imagine walking | 12 | 10.1007/s00221-008-1520-8 |
| Wagner,2008 | Motor | 12 | Imagine walking | 10 | 10.1007/s00221-008-1520-8 |
| Wagner,2008 | Motor | 12 | Imagine walking | 12 | 10.1007/s00221-008-1520-8 |
| Wagner,2008 | Motor | 12 | Imagine walking | 15 | 10.1007/s00221-008-1520-8 |
| Wagner,2008 | Motor | 12 | Imagine walking | 7 | 10.1007/s00221-008-1520-8 |
| Wagner,2008 | Motor | 12 | Imagine walking | 4 | 10.1007/s00221-008-1520-8 |
| Wagner,2008 | Motor | 12 | Imagine walking | 10 | 10.1007/s00221-008-1520-8 |
| Wang,2009 | Motor | 21 | Imagine leg movement | 32 | 10.1007/s00702-009-0269-y |
| Wang,2010 | Motor | 10 | Imagine finger movement | 6 | 10.1016/j.mri.2010.02.008 |
| Wang,2019 | Motor | 20 | Imagine hand movement | 10 | 10.3389/fnagi.2019.00312 |
| Wei,2010 | Motor | 12 | Imagine gymnastics and diving | 16 | 10.1016/j.brainres.2009.08.014 |
| Wei,2010 | Motor | 12 | Imagine gymnastics and diving | 10 | 10.1016/j.brainres.2009.08.014 |
| Willems,2009 | Motor | 32 | Imagine arm movement | 14 | 10.3389/neuro.09.039.2009 |
| Willems,2009 | Motor | 32 | Imagine arm movement | 22 | 10.3389/neuro.09.039.2009 |
| Willems,2010 | Motor | 20 | Imagine arm and whole body movement | 9 | 10.1162/jocn.2009.21386 |
| Wolbers,2003 | Motor | 13 | Imagine arm movement | 6 | 10.1093/cercor/13.4.392 |
| Wriessnegger,2014 | Motor | 23 | Imagine whole body movement | 8 | 10.3389/fnhum.2014.00469 |
| Wriessnegger,2016 | Motor | 21 | Imagine whole body movement | 4 | 10.1016/j.bandc.2016.08.008 |
| Wriessnegger,2016 | Motor | 21 | Imagine whole body movement | 10 | 10.1016/j.bandc.2016.08.008 |
| Wutte,2012 | Motor | 19 | Imagine leg movement | 20 | 10.1016/j.bbr.2011.09.042 |
| Wutte,2012 | Motor | 19 | Imagine leg movement | 5 | 10.1016/j.bbr.2011.09.042 |
| Yang,2014 | Motor | 20 | Imagine arm movement | 11 | 10.1007/s00221-013-3753-4 |
| Yang,2014 | Motor | 20 | Imagine arm movement | 6 | 10.1007/s00221-013-3753-4 |
| Yang,2014 | Motor | 20 | Imagine arm movement | 6 | 10.1007/s00221-013-3753-4 |
| Yang,2014 | Motor | 20 | Imagine arm movement | 6 | 10.1007/s00221-013-3753-4 |
| Zabicki,2017 | Motor | 20 | Imagine hand movement | 8 | 10.1093/cercor/bhw257 |
| Zanardi,2016 | Motor | 14 | Imagine hand rotation | 33 | 10.1016/j.bandc.2016.01.002 |
| Zapparoli,2013 | Motor | 24 | Imagine arm movement | 9 | 10.1007/s00221-012-3331-1 |
| Zapparoli,2014 | Motor | 30 | Imagine arm movement | 24 | 10.1007/s00221-014-4065-z |
| Zapparoli,2014 | Motor | 30 | Imagine arm movement | 19 | 10.1007/s00221-014-4065-z |
| Zapparoli,2016 | Motor | 46 | Imagine arm movement | 47 | 10.1111/ejn.13130 |
| Zapparoli,2016 | Motor | 46 | Imagine arm movement | 39 | 10.1111/ejn.13130 |
| Zapparoli,2019 | Motor | 22 | Imagine grasping movement | 93 | 10.1093/cercor/bhy314 |
| Zapparoli,2020 | Motor | 29 | Imagine walking | 23 | 10.1002/hbm.25123 |
| Zhang,2014 | Motor | 32 | Imagine finger movement | 2 | 10.1371/journal.pone.0085489 |
| Zich,2015 | Motor | 24 | Imagine arm movement | 7 | 10.1016/j.neuroimage.2015.04.020 |
| Zwergal,2012 | Motor | 20 | Imagine leg and whole body movement | 35 | 10.1016/j.neurobiolaging.2010.09.022 |
| Anderson,2019 | Visual | 14 | Imagine flickering circle | 8 | 10.1016/j.neuroimage.2019.06.057 |
| Banca,2015 | Visual | 20 | Imagine dot moving | 13 | 10.1088/1741-2560/12/6/066003 |
| Banca,2015 | Visual | 20 | Imagine dot moving | 4 | 10.1088/1741-2560/12/6/066003 |
| Beauregard,2009 | Visual | 15 | Imagine light | 10 | 10.1016/j.resuscitation.2009.05.006 |
| Boly,2007 | Visual | 12 | Imagine faces | 7 | 10.1016/j.neuroimage.2007.02.047 |
| Bonino,2015 | Visual | 10 | Imagine clock | 12 | 10.1016/j.neuropsychologia.2015.01.004 |
| Bray,2010 | Visual | 44 | Imagine rewards | 13 | 10.1152/jn.01030.2009 |
| Christian,2015 | Visual | 33 | Imagine pain scenario | 7 | 10.1162/jocn_a_00754 |
| Ciarlo,2022 | Visual | 18 | Imagine objects | 10 | 10.1088/1741-2552/ac6f81 |
| Costantini,2011 | Visual | 13 | Imagine images | 5 | 10.1016/j.neuropsychologia.2011.01.034 |
| Creem,2001 | Visual | 12 | Imagine self rotation | 15 | 10.3758/cabn.1.3.239 |
| Creem,2001 | Visual | 12 | Imagine self rotation | 13 | 10.3758/cabn.1.3.239 |
| Creem-Regehr,2007 | Visual | 23 | Imagine object manipulation | 18 | 10.1017/S1355617707071093 |
| Creem-Regehr,2007 | Visual | 23 | Imagine object manipulation | 26 | 10.1017/S1355617707071093 |
| Daselaar,2010 | Visual | 15 | Imagine images | 18 | 10.1016/j.neuroimage.2010.04.239 |
| de Borst,2012 | Visual | 3 | Imagine scene | 38 | 10.1016/j.neuroimage.2011.12.005 |
| Delamillieure,2010 | Visual | 7 | Imagine objects | 3 | 10.1016/j.brainresbull.2009.11.014 |
| D'Esposito,1997 | Visual | 7 | Imagine objects | 3 | 10.1016/s0028-3932(96)00121-2 |
| Diekhof,2011 | Visual | 9 | Imagine faces | 20 | 10.1016/j.neuroimage.2010.08.034 |
| Diekhof,2011 | Visual | 9 | Imagine faces | 9 | 10.1016/j.neuroimage.2010.08.034 |
| Dijkstra,2021 | Visual | 35 | Imagine images | 11 | 10.1523/ENEURO.0228-21.2021 |
| Dijkstra,2021 | Visual | 35 | Imagine images | 7 | 10.1523/ENEURO.0228-21.2021 |
| Direito,2020 | Visual | 10 | Imagine dot moving | 15 | 10.3389/fnhum.2020.578119 |
| Ganis,2004 | Visual | 15 | Imagine objects | 60 | 10.1016/j.cogbrainres.2004.02.012 |
| Ganis,2004 | Visual | 15 | Imagine objects | 26 | 10.1016/j.cogbrainres.2004.02.012 |
| Garbarini,2020 | Visual | 17 | Naming definition task | 4 | 10.1016/j.neuropsychologia.2019.107275 |
| Gardini,2005 | Visual | 15 | Imagine images | 7 | 10.1016/j.neuroimage.2005.04.032 |
| Gardini,2005 | Visual | 15 | Imagine images | 17 | 10.1016/j.neuroimage.2005.04.032 |
| Gardini,2009 | Visual | 13 | Imagine images | 23 | 10.1007/s00426-008-0175-1 |
| Gonsalves,2004 | Visual | 11 | Imagine images | 8 | 10.1111/j.0956-7976.2004.00736.x |
| Greening,2022 | Visual | 13 | Imagine fear scenario | 13 | 10.1038/s41598-022-05019-y |
| Han,2022 | Visual | 49 | Imagine objects | 12 | 10.1111/ejn.15654 |
| Hauk,2008 | Visual | 21 | Imagine action and object | 5 | 10.1111/j.1460-9568.2008.06143.x |
| Hayashi,2014 | Visual | 16 | Imagine word | 31 | 10.1016/j.neures.2013.10.007 |
| Hemati,2018 | Visual | 14 | Imagine animals | 124 | 10.3389/fnhum.2018.00515 |
| Hohenfeld,2017 | Visual | 16 | Imagine footpath | 18 | 10.3389/fneur.2017.00384 |
| Holland,2013 | Visual | 23 | Recall autobiographical memories | 21 | 10.1162/jocn_a_00289 |
| Holland,2013 | Visual | 23 | Recall autobiographical memories | 11 | 10.1162/jocn_a_00289 |
| Holland,2013 | Visual | 23 | Recall autobiographical memories | 34 | 10.1162/jocn_a_00289 |
| Huijbers,2011 | Visual | 21 | Imagine images | 6 | 10.1016/j.neuropsychologia.2011.02.051 |
| Ishai,2000 | Visual | 9 | Imagine house, face and chair | 10 | 10.1016/s0896-6273(00)00168-9 |
| Ishai,2000 | Visual | 9 | Imagine house, face and chair | 16 | 10.1016/s0896-6273(00)00168-9 |
| Izadifar,2022 | Visual | 1 | Imagine musical composition | 6 | 10.1002/pchj.522 |
| Kawamichi,2007 | Visual | 14 | Imagine object rotation | 30 | 10.1016/j.brainres.2007.01.082 |
| Kleider-Offutt,2019 | Visual | 28 | Imagine hair cut | 13 | 10.1007/s00221-019-05492-4 |
| Kosslyn,1993 | Visual | 7 | Imagine letter | 7 | 10.1162/jocn.1993.5.3.263 |
| Kosslyn,1993 | Visual | 7 | Imagine letter | 13 | 10.1162/jocn.1993.5.3.263 |
| Kosslyn,2005 | Visual | 16 | Imagine figure | 7 | 10.3758/cabn.5.1.41 |
| Kosslyn,2005 | Visual | 16 | Imagine figure | 10 | 10.3758/cabn.5.1.41 |
| Lacey,2010 | Visual | 8 | Imagine objects | 17 | 10.1016/j.neuroimage.2009.10.081 |
| Lacey,2010 | Visual | 8 | Imagine objects | 24 | 10.1016/j.neuroimage.2009.10.081 |
| Lazard,2011 | Visual | 10 | Imagine color | 11 | 10.1016/j.neuropsychologia.2011.04.025 |
| Lee,2014 | Visual | 14 | Imagine word | 6 | 10.1002/hbm.22512 |
| Logie,2011 | Visual | 21 | Imagine mental rotation | 2 | 10.1016/j.neuropsychologia.2011.07.011 |
| Mazard,2002 | Visual | 6 | Imagine shapes | 49 | 10.1162/089892902317236821 |
| Mellet,1996 | Visual | 9 | Imagine objects | 17 | 10.1523/JNEUROSCI.16-20-06504.1996 |
| Mellet,1996 | Visual | 9 | Imagine objects | 11 | 10.1523/JNEUROSCI.16-20-06504.1996 |
| Mellet,1998 | Visual | 8 | Imagine objects or animals | 15 | 10.1523/JNEUROSCI.16-20-06504.1996 |
| Mellet,1998 | Visual | 8 | Imagine objects or animals | 13 | 10.1523/JNEUROSCI.16-20-06504.1996 |
| Mohr,2009 | Visual | 12 | Imagine lines and circles | 3 | 10.1016/j.neuroimage.2009.03.045 |
| Olivetti,2009 | Visual | 9 | Imagine objects | 2 | 10.1016/j.actpsy.2009.06.009 |
| Olivetti,2009 | Visual | 9 | Imagine objects | 16 | 10.1016/j.actpsy.2009.06.009 |
| Roland,1995 | Visual | 11 | Imagine color | 5 | 10.1093/cercor/5.1.79 |
| Roland,1995 | Visual | 11 | Imagine color | 7 | 10.1093/cercor/5.1.79 |
| Roland,1995 | Visual | 11 | Imagine color | 10 | 10.1093/cercor/5.1.79 |
| Sasaoka,2014 | Visual | 23 | Imagine clock | 20 | 10.1162/jocn_a_00493 |
| Schicke,2006 | Visual | 11 | Imagine disc flying | 12 | 10.1111/j.1460-9568.2006.04720.x |
| Schicke,2006 | Visual | 11 | Imagine disc flying | 12 | 10.1111/j.1460-9568.2006.04720.x |
| Schienle,2008 | Visual | 24 | Imagine images | 14 | 10.1097/WNR.0b013e3282f85e10 |
| Schienle,2008 | Visual | 24 | Imagine images | 19 | 10.1097/WNR.0b013e3282f85e10 |
| Servos,2002 | Visual | 10 | Imagine objects | 8 | 10.1093/cercor/12.7.772 |
| Seurinck,2011 | Visual | 16 | Imagine mental rotation | 6 | 10.1162/jocn.2010.21525 |
| Thompson,2009 | Visual | 16 | Imagine mental rotation (images) | 9 | 10.1111/j.1467-9280.2009.02440.x |
| Tomasino,2022 | Visual | 19 | Imagine scene | 8 | 10.1002/hbm.25839 |
| Trojano,2000 | Visual | 7 | Imagine clock | 7 | 10.1093/cercor/10.5.473 |
| Trojano,2000 | Visual | 7 | Imagine clock | 10 | 10.1093/cercor/10.5.473 |
| Trojano,2000 | Visual | 7 | Imagine clock | 14 | 10.1093/cercor/10.5.473 |
| Weiler,2010 | Visual | 17 | Imagine past and future event | 6 | 10.1016/j.bbr.2010.04.013 |
| Weiler,2010 | Visual | 17 | Imagine past and future event | 18 | 10.1016/j.bbr.2010.04.013 |
| Whittingstall,2014 | Visual | 18 | Imagine spatial navigation | 14 | 10.1016/j.cortex.2013.02.004 |
| Yi,2008 | Visual | 9 | Imagine scene | 11 | 10.1162/jocn.2008.20094 |
| Zeman,2010 | Visual | 10 | Imagine faces | 20 | 10.1016/j.neuropsychologia.2009.08.024 |
| Zvyagintsev,2013 | Visual | 15 | Imagine objects | 11 | 10.1111/ejn.12140 |
| Zvyagintsev,2013 | Visual | 15 | Imagine objects | 8 | 10.1111/ejn.12140 |
| Boly,2007 | Auditory | 12 | Imagine song | 4 | 10.1016/j.neuroimage.2007.02.047 |
| Daselaar,2010 | Auditory | 15 | Imagine sound | 22 | 10.1016/j.neuroimage.2010.04.239 |
| Endestad,2020 | Auditory | 1 | Imagine melody | 10 | 10.3389/fnhum.2020.576888 |
| Grandchamp,2019 | Auditory | 24 | Imagine voice | 33 | 10.3389/fpsyg.2019.02019 |
| Halpern,2004 | Auditory | 10 | Imagine instrument | 7 | 10.1016/j.neuropsychologia.2003.12.017 |
| Herholz,2012 | Auditory | 10 | Imagine song | 2 | 10.1162/jocn_a_00216 |
| Herholz,2012 | Auditory | 10 | Imagine song | 5 | 10.1162/jocn_a_00216 |
| Herholz,2012 | Auditory | 10 | Imagine song | 9 | 10.1162/jocn_a_00216 |
| Huijbers,2011 | Auditory | 21 | Imagine sound | 6 | 10.1016/j.neuropsychologia.2011.02.051 |
| Izadifar,2022 | Auditory | 1 | Imagine melody | 3 | 10.1002/pchj.522 |
| Kim,2008 | Auditory | 23 | Imagine speech | 6 | 10.1016/j.neulet.2008.09.019 |
| Kim,2008 | Auditory | 23 | Imagine speech | 15 | 10.1016/j.neulet.2008.09.019 |
| Kleber,2007 | Auditory | 16 | Imagine song | 13 | 10.1016/j.neuroimage.2007.02.053 |
| Kleider-Offutt,2019 | Auditory | 28 | Imagine voice | 9 | 10.1007/s00221-019-05492-4 |
| Kleider-Offutt,2019 | Auditory | 28 | Imagine voice | 2 | 10.1007/s00221-019-05492-4 |
| Kurby,2013 | Auditory | 28 | Imagine sound | 13 | 10.1016/j.bandl.2013.07.003 |
| Kurby,2013 | Auditory | 28 | Imagine sound | 9 | 10.1016/j.bandl.2013.07.003 |
| Lazard,2011 | Auditory | 10 | Imagine sound | 9 | 10.1016/j.neuropsychologia.2011.04.025 |
| Lazard,2011 | Auditory | 10 | Imagine sound | 3 | 10.1016/j.neuropsychologia.2011.04.025 |
| Lazard,2013 | Auditory | 10 | Imagine sound | 10 | 10.1002/hbm.21504 |
| Lazard,2013 | Auditory | 10 | Imagine sound | 4 | 10.1002/hbm.21504 |
| Leaver,2009 | Auditory | 20 | Imagine melody | 4 | 10.1523/JNEUROSCI.4921-08.2009 |
| Leaver,2009 | Auditory | 20 | Imagine melody | 8 | 10.1523/JNEUROSCI.4921-08.2009 |
| Leclerc,2019 | Auditory | 53 | Imagine word | 11 | 10.1093/chemse/bjz055 |
| Lu,2021 | Auditory | 24 | Imagine poem | 2 | 10.1016/j.neuroimage.2021.117724 |
| Lu,2021 | Auditory | 24 | Imagine poem | 12 | 10.1016/j.neuroimage.2021.117724 |
| Lu,2023 | Auditory | 30 | Imagine poem | 8 | 10.1093/cercor/bhac519 |
| Lu,2023 | Auditory | 30 | Imagine poem | 6 | 10.1093/cercor/bhac519 |
| Neef,2016 | Auditory | 32 | Imagine melody | 57 | 10.1016/j.neuroimage.2016.08.030 |
| Neef,2016 | Auditory | 32 | Imagine melody | 32 | 10.1016/j.neuroimage.2016.08.030 |
| Olivetti,2009 | Auditory | 9 | Imagine sound | 3 | 10.1016/j.actpsy.2009.06.009 |
| Olivetti,2009 | Auditory | 9 | Imagine sound | 5 | 10.1016/j.actpsy.2009.06.009 |
| Olivetti,2009 | Auditory | 9 | Imagine sound | 8 | 10.1016/j.actpsy.2009.06.009 |
| Palmiero,2009 | Auditory | 9 | Imagine whole body movement | 2 | 10.1007/s10339-009-0324-5 |
| Rudner,2005 | Auditory | 12 | Imagine word | 10 | 10.1016/j.bandl.2004.05.010 |
| Shergill,2001 | Auditory | 8 | Imagine word | 12 | 10.1017/s003329170100335x |
| Shergill,2001 | Auditory | 8 | Imagine word | 9 | 10.1017/s003329170100335x |
| Shergill,2001 | Auditory | 8 | Imagine word | 10 | 10.1017/s003329170100335x |
| Simmonds,2014 | Auditory | 17 | Imagine word | 28 | 10.1523/JNEUROSCI.0336-14.2014 |
| Tian,2016 | Auditory | 18 | Imagine syllabe | 11 | 10.1016/j.cortex.2016.01.002 |
| Tsai,2018 | Auditory | 14 | Imagine word, sound, music | 37 | 10.1016/j.neuropsychologia.2018.07.028 |
| Tsai,2018 | Auditory | 14 | Imagine word, sound, music | 6 | 10.1016/j.neuropsychologia.2018.07.028 |
| Tsai,2018 | Auditory | 14 | Imagine word, sound, music | 36 | 10.1016/j.neuropsychologia.2018.07.028 |
| Yoo,2001 | Auditory | 12 | Imagine sound | 21 | 10.1097/00001756-200110080-00013 |
| Zatorre,2010 | Auditory | 12 | Imagine song | 36 | 10.1162/jocn.2009.21239 |
| Zatorre,2010 | Auditory | 12 | Imagine song | 21 | 10.1162/jocn.2009.21239 |
| Zvyagintsev,2013 | Auditory | 15 | Imagine melody | 7 | 10.1111/ejn.12140 |
| Zvyagintsev,2013 | Auditory | 15 | Imagine melody | 15 | 10.1111/ejn.12140 |
| Nierhaus,2023 | Tactile | 21 | Imagine braille | 12 | 10.1523/ENEURO.0408-22.2023 |
| Olivetti,2009 | Tactile | 9 | Imagine touching something | 3 | 10.1016/j.actpsy.2009.06.009 |
| Olivetti,2009 | Tactile | 9 | Imagine touching something | 4 | 10.1016/j.actpsy.2009.06.009 |
| Olivetti,2009 | Tactile | 9 | Imagine touching something | 7 | 10.1016/j.actpsy.2009.06.009 |
| Schmidt,2014 | Tactile | 14 | Imagine vibration on body part | 27 | 10.1016/j.neuroimage.2014.05.014 |
| Schmidt,2019 | Tactile | 19 | Imagine vibration on body part | 28 | 10.3389/fnhum.2019.00010 |
| Tomasino,2022 | Tactile | 19 | Imagine sensation | 13 | 10.1002/hbm.25839 |
| Yoo,2003 | Tactile | 13 | Imagine tactile stimulation | 11 | 10.1097/00001756-200303240-00011 |
| Yoo,2003 | Tactile | 13 | Imagine tactile stimulation | 6 | 10.1097/00001756-200303240-00011 |
| Avery,2023 | Gustatory | 22 | Imagine taste of substance | 4 | 10.1016/j.pneurobio.2023.102423 |
| Avery,2023 | Gustatory | 22 | Imagine taste of substance | 11 | 10.1016/j.pneurobio.2023.102423 |
| Kikushi,2005 | Gustatory | 13 | Imagine taste of substance | 6 | 10.1097/00001756-200502280-00016 |
| Kobayashi,2004 | Gustatory | 18 | Imagine taste of substance | 23 | 10.1016/j.neuroimage.2004.08.002 |
| Olivetti,2009 | Gustatory | 9 | Imagine taste of substance | 2 | 10.1016/j.actpsy.2009.06.009 |
| Olivetti,2009 | Gustatory | 9 | Imagine taste of substance | 4 | 10.1016/j.actpsy.2009.06.009 |
| Olivetti,2009 | Gustatory | 9 | Imagine taste of substance | 3 | 10.1016/j.actpsy.2009.06.009 |
| Han,2022 | Olfactory | 49 | Imagine smell of substance | 7 | 10.1111/ejn.15654 |
| Leclerc,2019 | Olfactory | 48 | Imagine smell of substance | 17 | 10.1093/chemse/bjz055 |
| Leclerc,2019 | Olfactory | 48 | Imagine smell of substance | 3 | 10.1093/chemse/bjz055 |
| Olivetti,2009 | Olfactory | 9 | Imagine smell of substance | 6 | 10.1016/j.actpsy.2009.06.009 |
| Olivetti,2009 | Olfactory | 9 | Imagine smell of substance | 3 | 10.1016/j.actpsy.2009.06.009 |
| Olivetti,2009 | Olfactory | 9 | Imagine smell of substance | 6 | 10.1016/j.actpsy.2009.06.009 |
| Plaily,2012 | Olfactory | 28 | Imagine smell of substance | 8 | 10.1002/hbm.21207 |
